# Optimal energy and redox metabolism in the cyanobacterium *Synechocystis* sp. PCC 6803

**DOI:** 10.1101/2022.09.14.507938

**Authors:** Amit Kugler, Karin Stensjö

## Abstract

Cyanobacteria represent an attractive platform for the sustainable production of chemicals and fuels. However, the obtained rates, yields, and titers are below those required for commercial application. Carbon metabolism alone cannot achieve maximal accumulation of end-products, since an efficient production of target molecules entails energy and redox balance, in addition to carbon flow. The interplay between cofactor regeneration and heterologous metabolite overproduction in cyanobacteria is not fully explored. Here, we applied stoichiometric metabolic modelling of the cyanobacterium *Synechocystis* sp. PCC 6803, in order to investigate the optimality of energy and redox metabolism, while overproducing bio-alkenes - isobutene, isoprene, ethylene and 1-undecene. Our network-wide analysis indicates that the rate of NADP+ reduction, rather than ATP synthesis, controls ATP/NADPH ratio, and thereby chemical production. The simulation implies that energy and redox balance necessitates gluconeogenesis, and that acetate metabolism via phosphoketolase serves as an efficient carbon- and energy-recycling pathway. Furthermore, we show that an auxiliary pathway, composed of serine, one-carbon and glycine metabolism, supports cellular redox homeostasis and ATP cycling, and that the *Synechocystis* metabolism is controlled by few key reactions carrying a high flux. The study also revealed non-intuitive metabolic pathways to enhance isoprene, ethylene and 1-undecene production. We conclude that metabolism of ATP and NAD(P)H is entwined with carbon and nitrogen metabolism, and cannot be assessed in isolation. We envision that the presented here in-depth metabolic analysis will guide the a priori design of *Synechocystis* as a host strain for an efficient manufacturing of target products.

## Introduction

The biosynthesis of various cell metabolites involves biochemical reactions, which require cofactors, such as adenosine triphosphate (ATP) and reduced nicotinamide adenine dinucleotides (NAD(P)H). ATP is the major energy carrier in the cell, driving numerous metabolic pathways. The pyridine nucleotides NAD(H) and NADP(H) serve as reducing equivalents in redox reactions. While NADH participates mainly in catabolic reactions, NADPH plays a role in anabolic reactions.

Cyanobacteria serve as an attractive platform for the sustainable production of chemicals and fuels, due to the capability of converting sunlight and atmospheric carbon dioxide into organic compounds, and the ease of genetic manipulation. In fact, in the past years, the unicellular cyanobacterium *Synechocystis* sp. PCC 6803 (hereafter *Synechocystis*) has been successfully used as a cell factory for the production of commodity chemicals, including hydrogen,^1^ L-lactic acid,^2^ ethanol^3^ and others.^4^ Besides photoautotrophy, cyanobacteria are able to grow under mixotrophic conditions. Mixotrophic metabolism allows the cells to assimilate CO_2_ in parallel with the consumption of organic carbon, while exploiting light energy. The presence of organic substrates in light growth conditions is considered as a promising cultivation strategy for the commercialization of cyanobacteria.^5,6^ This trophic mode enables enhanced biomass^7^ and chemical yield,^8,9^ and thereby confers industrial applicability. In cyanobacteria, the photosynthetic electron transport chain is the major source for ATP and NADPH generation, when grown under light conditions.^10^ The different glycolytic routes and the TCA cycle generate reducing agent in the form of NADH. Additionally, transhydrogenases catalyse the interconversion between NAD(H) and NADP(H).^11^

An optimal performance of photosynthesis requires that the light-energy conversion by the photosystems and the downstream metabolic pathways are fine-tuned. This, to support the ATP/NADPH output ratio, an important parameter for evaluating the cellular energy economy.^12^ Maintaining intracellular energy and redox state is essential to support cell survival and viability through routine production and degradation of nucleotides, lipids and proteins. The maximum ATP/NADPH ratio that can be generated by the photosynthetic linear electron flow (LEF) is 1.28 (9 ATP and 7 NADPH). Alternative electron flow (AEF) pathways, such as the cyclic electron flow (CEF), generate additional ATP and thus support the demand for downstream biochemical pathways, such as the Calvin–Benson–Bassham (CBB) cycle and photorespiration (1.5–1.67).^13^ In addition, re-oxidation of NAD(P)H to NAD(P)+ and phosphorylation-dephosphorylation of reducing equivalents are important for balancing the ATP/NADPH budget and maintaining redox homeostasis.^14^ An imbalance between the supply and demand for reducing power and chemical energy results in metabolic congestion, and can be one of the rate-limiting factors impeding photosynthetic efficiency.

Despite of the importance of meeting the energy and redox requirement to achieve an efficient cellular performance,^15,16^ metabolic engineering efforts of *Synechocystis* have been mainly focused on channelling the carbon skeleton into the end-product. This is, probably, due to the conception that carbon (i.e., precursor availability) is considered as the main factor that controls microbial growth and bio-based chemical production. In addition, cyanobacteria are characterized by a large NADPH pool,^17,18^ as compared to heterotrophic bacteria,^19^ which favour the exploitation of NADPH-dependent enzymes, a rich source of biocatalysts with high potential for the production of valuable chemicals.

Computational tools, such as constraint-based modelling (CBM), have been widely used for the rational strain optimization. CBM is based on imposing of a set of constraints that govern the operation of a metabolic network at steady state. These include, for example, the stoichiometry of the biochemical reactions, mass balance and thermodynamic laws. Flux balance analysis (FBA) is a common approach to calculate, under given constraints, the intracellular flux distributions within the stoichiometric network, by optimizing an objective function.^20^ As such, FBA can be used to predict the maximum (or minimum) growth rate or the production of desired metabolites, as well as to identify genetic interventions that force carbon flux towards chosen compound. In the past decade, a few model-driven approaches have been applied in *Synechocystis* to guide metabolic engineering for the production of n-butanol, 1,3-propanediol and limonene.^21^

Alkenes are commercially valuable platform chemicals, traditionally used as detergents, lubricants and rubbers. In addition, they are compatible hydrocarbon fuels, due to their high energy content.^22^ The development of gas-to-liquid technologies, enabling the oligomerization of small gaseous substrates into liquid chemicals, together with the environmental concerns associated with the use of fossil fuels and carbon dioxide emissions,^23^ are driving the necessity for an environmentally-friendly, yet economically-feasible, manufacturing process of bio-alkenes.^24–26^

In this work, we addressed the question of how the model cyanobacterium *Synechocystis* balances its anabolic-catabolic processes, with respect to central cofactor metabolites (ATP, NADPH and NADH), during growth under photoautotrophic and mixotrophic conditions. To this end, we employed genome-scale metabolic modelling, and analysed the metabolism of *Synechocystis* strains overproducing alkenes as case study.

## Materials and methods

### Genome-scale modelling

The constraint-based analysis of the metabolic network evaluates the steady-state flux distribution, which satisfies the flux balance condition for any internal metabolites. By specifying cellular objective to be optimized, the metabolic model can be formulated as a linear programming (LP) problem (Eq. 1-3).

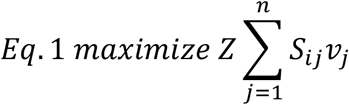

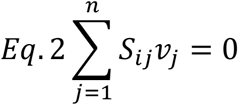

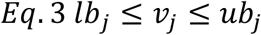

Where *S*_*ij*_ refers to the stoichiometric coefficient of metabolite i participating in reaction j, and the *v* _*j*_ refers to the vector of reaction flux (mmol/gDW/h; gDW, gram dry weight) of reaction j at steady state. The flux *v* _*j*_ is the j-th component of an n-dimensional flux vector v, where n is the total number of fluxes. lb and ub correspond to the lower bound and upper bound, respectively, of the estimated rate for the reaction j.

Parsimonious FBA (pFBA)^27^ was employed on COBRApy package^28^ using the commercial solver Gurobi 9.5.2 (Gurobi Optimization, Inc., Houston, TX, United States) in Python 3.8. The iJN678 genome-scale model of *Synechocystis* sp. PCC 6803 metabolism,^29^ downloaded from the BiGG database,^30^ was used for the computational analysis. The metabolic network was updated in accordance with the literature (Supplementary Table 1), and include the tricarboxylic acid (TCA) cycle shunt reactions,^31–33^ the phosphoketolase reactions,^34,35^ the Entner–Doudoroff (ED) pathway,^36^ the light-independent L-serine biosynthesis pathway,^37^ modifications in tyrosine biosynthesis,^38^ a modification to transhydrogenase^11^ and modifications in the electron-transport chain reactions.^39,40^ In addition, non-growth associated ATP maintenance (NGAM) reaction was added to the model. The NGAM coefficient^41^ is represented by a pseudo-reaction that degrades ATP into ADP and orthophosphate (Pi) (Eq. 4). For the evaluation of heterologous alkene(s) production in *Synechocystis*, four biosynthesis pathways^42–45^ were separately implemented into the iJN678 model (Supplementary Tables 2-5), yielding iJN678_isobutene, iJN678_isoprene, iJN678_ethylene and iJN678_1-undecene.

Due to mass balance considerations and unknown mechanism of RnKICD activity, we choose to simulate the isobutene production by including the M3K reaction in the metabolic network of iJN678_isobutene model, as described in.^46,47^ In each reconstructed model, a cytoplasmatic sink for the alkene was introduced and set as the objective function. In addition, in order to model a more realistic scenario, that is, to enable high production rate of end-products whilst sustain biomass synthesis, the minimum flux through the biomass objective function^48^ was set to 10% of the maximal growth rate. Autotrophic metabolism was simulated by constraining the bicarbonate uptake rate to 3.7 mmol/gDW/h and the photon uptake rate to 45 mmol/gDW/h. Mixotrophic metabolism was simulated by constraining the carbon dioxide, bicarbonate, glucose and photon uptake rate to 0, 3.7, 0.38 and 45 mmol/gDW/h, respectively. Visualization of the metabolic reaction network and the obtained fluxes was done using the Escher web-tool^49^.

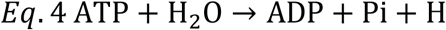

To explore the flux solution space, flux variability analysis (FVA)^50^ was performed (Supplementary Data 1-10). For FVA, the fraction of the optimum was set to 95%, which accounts for 5% variation around the best-known objective value.

The sensitivity of the FBA solution was indicated by shadow prices (Eq. 5).^41^ The shadow prices are calculated in the dual solution to the linear programming problem, and can be used to interpret shifts from one optimal flux distribution to another. These indicate how much the corresponding metabolite would increase or decrease the objective function by a one unit.

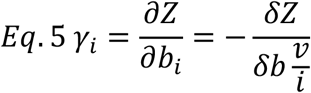

Where Z refers to the objective function of FBA, and *b*_*i*_ denotes the additional availability for the metabolite i.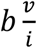defines the violation of a mass balance constraint and is equivalent to an uptake reaction.

The flux-sum analysis was used to quantify the cofactors turnover rates among the metabolic network.^51^ The flux-sum values for a metabolite represent the sum of flux in all the reactions that consume or produce it, multiplied by the corresponding stoichiometric coefficients (Eq. 6).

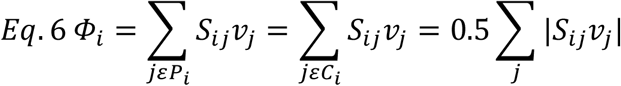

Where *S*_*ij*_ refers to the stoichiometric coefficient of metabolite i participating in reaction j, and *v* _*j*_ refers to the flux (mmol/gDW/h) of reaction j. *j* denotes the set of reactions producing metabolite i, and *j* denotes the set of reactions consuming metabolite i. Under the steady-state conditions, the consumption and production rates for any metabolite are equal. Thus, the turnover rate of metabolite i is half the absolute sum of consumption and the generation rates.

We then weighted the cellular metabolic costs by multiplying the shadow prices with the total flux passing through each cofactor metabolite (calculated by flux-sum) (Eq. 7). The rationale behind doing this is that, cofactors are interconnected with immense number of biochemical reactions, which affect the extent of utilization.

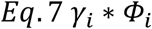

The theoretical maximum yield is expressed as gram product produced per gram of substrate consumed. The carbon-conversion efficiency was estimated by calculating carbon-mole (c-mole) produced per carbon-mole of substrate consumed.

## Results and discussion

Engineering metabolism for biotechnological purposes requires that the host organism can support genetic modification being introduced. Modifications to energy and redox metabolism are commonplace when engineering microorganisms, as they induce a cellular imbalance of ATP and NAD(P)H, which often results in decreased growth performance and biosynthetic capacity of the engineered strain.^52^

To enable a system analysis of alkene-overproducing *Synechocystis* strains, we chose four alkenes whose production using cyanobacteria has been previously demonstrated, stem from different metabolic routes and possess different energy and redox requirements: isobutene,^43^ isoprene,^45^ ethylene^44^ and 1-undecene (Fig. 1).^53^ All the selected production pathways start with the conversion of glucose to pyruvate via glycolytic pathways. Isobutene synthesis, that proceeds via the branched-chain amino acids pathway from 2-ketoisocaproate, requires one molecule of NADP^+^, one NAD^+^ and one ATP, and in total releases four molecules of CO_2_. Isoprene production through methyl-erythritol-4-phosphate (MEP) is a derivative of the terpenoid pathway, and consumes three molecules of NADPH and one molecule of ATP, alongside the release of one molecule of CO_2_. Ethylene synthesis is derived from the TCA cycle, for which one molecule of NAD^+^ and one of NADP^+^ are utilized and seven molecules of CO_2_ are released. 1-undecene is a fatty-acid derived alkene, and its biosynthesis pathway is characterized by the elongation and reduction of the aliphatic chain from acetyl-coenzyme A (acetyl-CoA) and malonyl-CoA, in the presence of 10 NADPH molecules (Fig. 1).

**Fig. 1.**
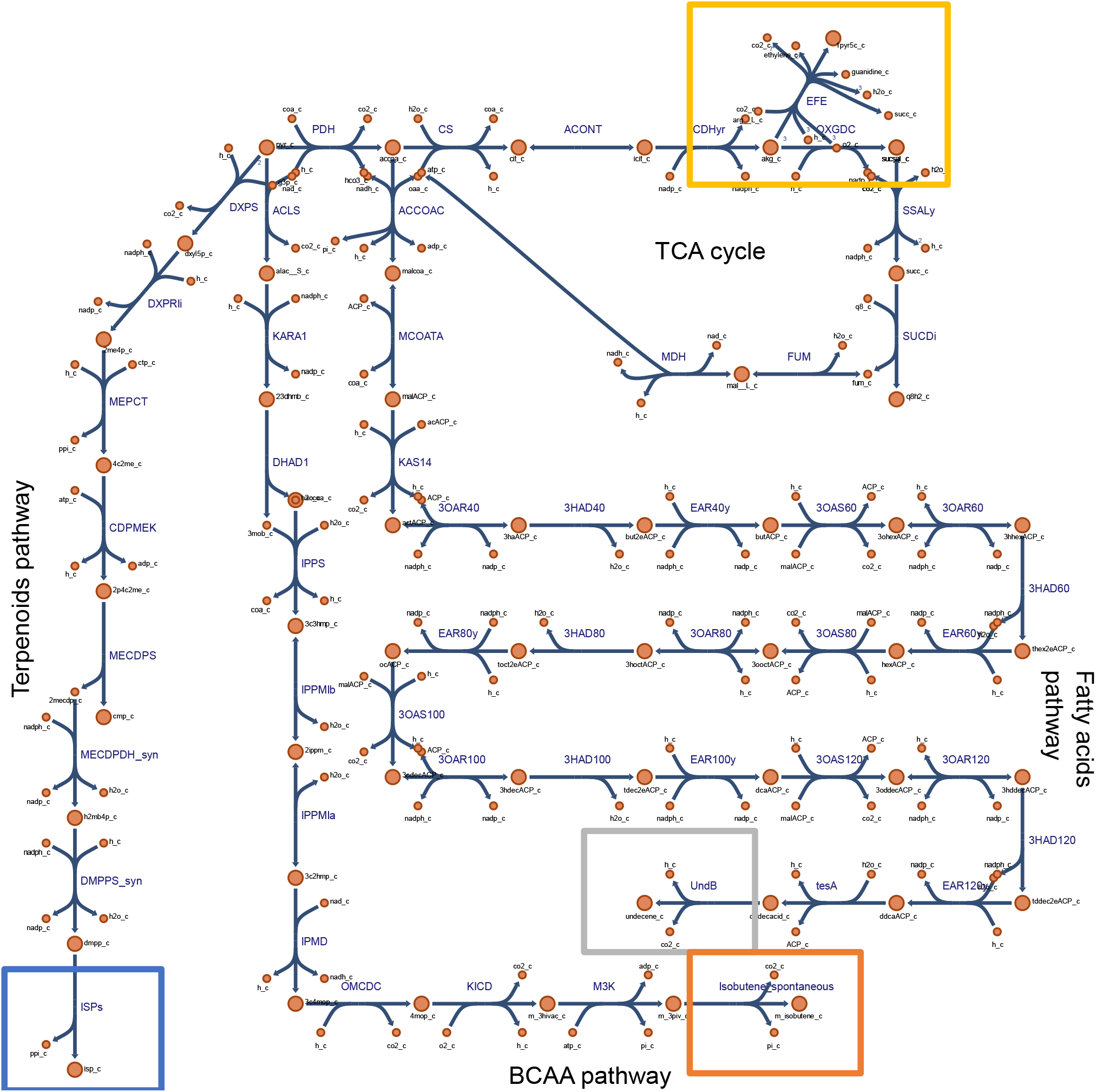
Non-native alkene biosynthesis pathways reconstructed in the iJN678 *Synechocystis* sp. PCC 6803 genome-scale model for cofactor balance analysis. TCA, tricarboxylic acid; BCAA, branched-chain amino acids. Metabolic reactions and metabolites (except heterologous ones) are indicated by their BiGG identifier.^30^

### Metabolite-centric analysis

In contrast to the traditional use of FBA for metabolic engineering, which rely on a reaction-centric approach, we chose to first analyse *Synechocystis* metabolism from the metabolite-centric point of view. In this way, it is possible to predict how the fluxes of reactions surrounding a metabolite of interest are adjusted to satisfy the optimization of the objective function.

We sought to analyse the overall metabolic requirements of ATP and NAD(P)H for synthesizing one mmol of the respective alkene, rather than focusing on the actual molar demands of single biosynthesis pathways. This, to provide an appropriate context and biologically meaningful mechanistic basis for guidance of metabolic engineering approaches. Even though the net accumulation of a metabolite might be zero, the overall turnover rate inside the cell is an indicator of its importance.

Under the steady-state assumption, the overall consumption and production rates of metabolites are equal, and can be evaluated using flux-sum analysis,^51^ which examines the interconversion pattern of a metabolite in the network. On the basis of the flux-sum analysis (Table 1), a minor usage of NADH was observed (turnover of 0.01-0.35 mmol/gDW/h), as compared to that of ATP (turnover of 7.19-8.15 mmol/gDW/h) and NADPH (turnover of 3.87-5.48 mmol/gDW/h), in the *Synechocystis* strains grown photoautotrophically. In addition, all strains exhibited a comparable ATP requirement for biomass and alkene synthesis. A high NADPH turnover rate was determined for isobutene and 1-undecene production (5.48 mmol/gDW/h, respectively), with lower values for isoprene, ethylene and biomass production. NADH turnover rate was the highest for ethylene production (0.49 mmol/gDW/h) and isobutene (0.70 mmol/gDW/h), whereas the lowest for isoprene and 1-undecene (0.01 mmol/gDW/h). A higher NADH turnover rate (1.5-fold) was observed under mixotrophic conditions, as compared to under phototrophic mode. An explanation for these observations is that the introduced biosynthesis pathways are mainly anabolic, and therefore use more NADPH than NADH; under mixotrophic conditions, the regeneration of NADH cofactor is more active. In summary, the calculated usage of NADPH did not correlate with the actual molar of cofactor demanded by the different alkene metabolic pathways. On the other hand, NADH usage did show a correlation with the NADH reliance to drive the metabolic pathways (Fig. 1, Table 1).

**Table 1.**
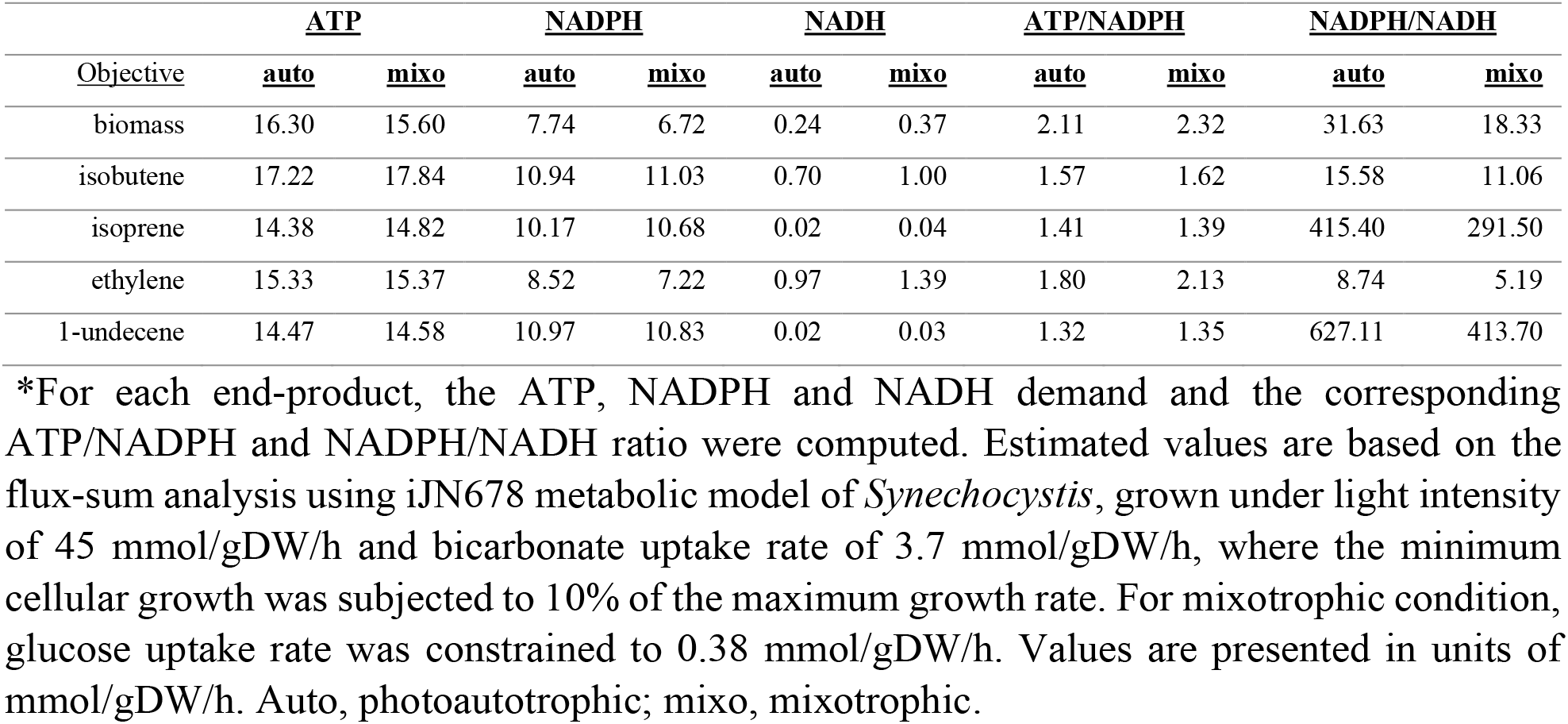
Demand of energy and redox resources for the oversynthesis of alkene end-products in *Synechocystis*, simulated to grow under photoautotrophic or mixotrophic conditions.

| <u>Objective</u> | <u>ATP</u> |  | <u>NADPH</u> |  | <u>NADH</u> |  | <u>ATP/NADPH</u> |  | <u>NADPH/NADH</u> |  |
| --- | --- | --- | --- | --- | --- | --- | --- | --- | --- | --- |
|  | <u>auto</u> | <u>mixo</u> | <u>auto</u> | <u>mixo</u> | <u>auto</u> | <u>mixo</u> | <u>auto</u> | <u>mixo</u> | <u>auto</u> | <u>mixo</u> |
| biomass | 16.30 | 15.60 | 7.74 | 6.72 | 0.24 | 0.37 | 2.11 | 2.32 | 31.63 | 18.33 |
| isobutene | 17.22 | 17.84 | 10.94 | 11.03 | 0.70 | 1.00 | 1.57 | 1.62 | 15.58 | 11.06 |
| isoprene | 14.38 | 14.82 | 10.17 | 10.68 | 0.02 | 0.04 | 1.41 | 1.39 | 415.40 | 291.50 |
| ethylene | 15.33 | 15.37 | 8.52 | 7.22 | 0.97 | 1.39 | 1.80 | 2.13 | 8.74 | 5.19 |
| 1-undecene | 14.47 | 14.58 | 10.97 | 10.83 | 0.02 | 0.03 | 1.32 | 1.35 | 627.11 | 413.70 |

\*For each end-product, the ATP, NADPH and NADH demand and the corresponding ATP/NADPH and NADPH/NADH ratio were computed. Estimated values are based on the flux-sum analysis using iJN678 metabolic model of *Synechocystis*, grown under light intensity of 45 mmol/gDW/h and bicarbonate uptake rate of 3.7 mmol/gDW/h, where the minimum cellular growth was subjected to 10% of the maximum growth rate. For mixotrophic condition, glucose uptake rate was constrained to 0.38 mmol/gDW/h. Values are presented in units of mmol/gDW/h. Auto, photoautotrophic; mixo, mixotrophic.

Since the overall consumption and production rates are equal under the steady-state assumption in FBA, the flux-sum also serves as a proxy of the metabolite pool size in the system. This allowed us to examine the cellular ATP/NADPH budget (Table 1), a vital parameter for efficient photosynthesis-driven metabolism.^12^ For phototrophic growth, it was computed that biomass accumulation requires an ATP/NADPH ratio of above 2. In addition, a lower ATP/NADPH ratio of ca. 1.80 was needed for optimal production of ethylene, followed by isobutene (1.57) and isoprene (1.41). 1-undecene required an ATP/NADPH ratio of 1.31, a ratio slightly above the value provided by the LEF. This means that, the 1-undecene biosynthesis affects the cellular metabolism the least, and is the most energetically balanced pathways. In that respect, the finding of ATP/NADPH ratios below that required for biomass accumulation means that excess ATP must be hydrolysed or channelled into other cellular processes. It should be noted that, the values presented here are slightly different from previous published results,^54,55^ due to differences related to model complexity, as well as to initial configuration, described in the materials and methods section. Under mixotrophic conditions, a similar trend of required ATP/NADPH ratios, but a slightly higher value was determined (Table 1).

The NADPH/NADH ratio was substantially altered (Table 1) between photoautotrophic conditions and mixotrophic conditions. For example, optimization with respect to maximal growth rate resulted in a ratio of 31.63 moles of NADPH per mole of NADH under photoautotrophic conditions, whereas only 18.33 moles of NADPH per mole of NADH under mixotrophic conditions. This is accordance with the higher NADH turnover computed under mixotrophic conditions.

In summary, a relatively solid flux-sum for ATP was observed under both growth conditions. For achieving a higher ATP/NADPH ratio than that provided by the LEF, more ATP is to be supplied, or, alternatively, NADPH must be degenerated. For a lower ratio, ATP is to be dissipated, or, alternatively, additional NADPH must be provided. A previous experimental study presumed that, under steady-state photosynthesis, *Chlamydomonas reinhardtii* is not limited by the availability of ATP, and that the rate of NADPH generation is critical for CO_2_ assimilation.^56^ Consistent with these results, we conclude that, under light-limited condition, the ATP/NADPH ratios are mainly governed by the intracellular levels NADPH, rather than that of ATP.

Mixotrophy has been attributed to a higher biosynthesis performance, as compared to photoautotrophic mode. We subsequently examined the effect of photoautotrophic and mixotrophic metabolism on the theoretical maximum production capability, a parameter of interest from both biotechnological and biochemical aspects. Microbial energy metabolism is one the effectors of product yield, since the biosynthesis of the nucleotides adenosine monophosphate (AMP) and nicotinamide adenine dinucleotide (NAD^+^), the precursors for ATP, NADPH and NADH, is by itself also energetically expensive. The theoretical analysis of production potential revealed that under photoautotrophic conditions, the maximum productivity for isobutene, isoprene and ethylene was comparable (0.65-0.68 mmol/gDW/h), while the value for 1-undecene was lower (0.30 mmol/gDW/h) (Table 2). The maximum theoretical mass yield for isobutene, isoprene and 1-undecene was similar (0.20 gram/gram), with a lower value observed for ethylene (0.15 gram/gram). The determined c-mol yields were comparable for isobutene, isoprene and 1-undecene (0.05 c-mol/c-mol), with a lower value for ethylene (0.03 c-mol/c-mol). Under mixotrophic conditions, similar maximum production patterns were observed, albeit at higher values. The calculated mass yields for isobutene, isoprene, ethylene, and 1-undecene were 0.79, 1.00, 0.38 and 0.95 gram/gram, respectively.

**Table 2.** Theoretical maximum productivity and yield of the studied alkene end-products in *Synechocystis* simulated to grow under photoautotrophic or mixotrophic conditions.

| <u>Objective</u> | <u>Photoautotrophy</u> |  |  | <u>Mixotrophy</u> |  |  |
| --- | --- | --- | --- | --- | --- | --- |
|  | <u>Maximum productivity</u> | <u>Maximum theoretical mass yield</u> | <u>C-mol yield</u> | <u>Maximum productivity</u> | <u>Maximum theoretical mass yield</u> | <u>C-mol yield</u> |
| biomass | 0.08 | 0.02 |  | 0.12 | 0.32 |  |
| isobutene | 0.68 | 0.20 | 0.05 | 0.96 | 0.79 | 0.37 |
| isoprene | 0.67 | 0.20 | 0.05 | 1.00 | 1.00 | 0.45 |
| ethylene | 0.65 | 0.15 | 0.03 | 0.92 | 0.38 | 0.18 |
| 1-undecene | 0.30 | 0.21 | 0.05 | 0.42 | 0.95 | 0.44 |

\*Maximum productivity is expressed as h<sup>-1</sup> or mmol/gDW/h, for biomass and alkenes, respectively. Productivity is presented in units of mmol/gDW/h. Mass yield is presented in units of gram product per gram substrate. Biomass yield is represented in units of gDW biomass per mmol carbon consumed. C-mol yield is presented in units of mol carbon produced per mol carbon consumed. gDW, gram dry-weight; h, hour; C, carbon.

0.37, 0.45, 0.18 and 0.44 c-mol/c-mol were computed for isobutene, isoprene, ethylene, and 1-undecene, respectively (Table 2). In summary, by comparing carbon-source-dependent bioproduction, our computational analysis illustrated that (i) mixotrophic metabolism is advantageous over photoautotrophy for the production of the examined alkenes, (ii) the carbon-conversion efficiency is highest for production of isoprene and 1-undecene, lower for isobutene, and the least for ethylene synthesis. The low yields found can be attributed to the high carbon loss (computed by flux-sum). For example, under photoautotrophic conditions, biomass production releases 3.81 mmol of CO_2_, whereas isobutene and ethylene production results in a release of 5.80 and 5.28 mmol CO_2_, respectively (Supplementary Table 6). This infers that the examined isobutene and ethylene biosynthesis pathways are relatively inefficient for bioproduction (iii) in terms of production rate, 1-undecene biosynthesis is less efficient.

In order to determine which of the studied cofactors imposed the greatest burden on the objective function, we then explored the sensitivity of the objective function to changes in the availability of individual cofactors. We employed flux imbalance analysis approach,^57^ and analyzed the marginal cost of the cofactor metabolites, known as shadow price.^41^ Thereby, a negative value implies that the corresponding metabolite is limiting the objective function, whereas a positive value implies that the intracellular flux of the metabolite is sufficient for reaching the objective function. In this study, the shadow price denotes the usefulness of an individual cofactor metabolite in accelerating the biomass or alkenes production rate. Following weighting, it can be seen that the biosynthesis of energy and redox carriers delimited the in-silico overproduction of all the investigated objective functions (Table 3). For instance, under photoautotrophic conditions, ATP and NADPH were shown to incur the heaviest cost for isobutene synthesis, followed by isoprene, ethylene and 1-undecene. Under mixotrophic conditions, similar trends were observed, where the weighted cost for NADPH displayed a slightly higher value than isobutene. Enhancing the NADH levels were shown to result in a higher increase for ethylene biosynthesis, followed by isobutene, with almost no additional increase in isoprene and 1-undecene production. Interestingly, the addition of NADPH was not found to result in highest production for 1-undecene biosynthesis, a heavily NADPH-dependent metabolic pathway. In summary, the shadow prices did not directly represent the number of cofactors needed for the biosynthesis of the individual alkenes. This analysis implied that a higher cellular availability of ATP and NAD(P)H individually is beneficial for biosynthesis. A higher cofactor availability of NAD(P)H can be achieved by (i) heterologous introduction of non-native oxidoreductases with specificity for another cofactor,^58^ (ii) protein engineering to switch the cofactor-dependency of enzymes,^59^ (iii) knocking out competing cofactor-utilizing pathways and (iiii) increasing the flux in cofactor-producing reactions (e.g. electron transport chain).

**Table 3.** Weighted costs of cofactor metabolites (ATP, NADPH and NADH) in biomass and alkene productivity, under photoautotrophic and mixotrophic conditions.

| Objective | <u>Photoautotrophy</u> |  |  | <u>Mixotrophy</u> |  |  |
| --- | --- | --- | --- | --- | --- | --- |
|  | <u>ATP</u><br><u>Weighted costs</u> | <u>NADPH</u><br><u>Weighted costs</u> | <u>NADH</u><br><u>Weighted costs</u> | <u>ATP</u><br><u>Weighted costs</u> | <u>NADPH</u><br><u>Weighted costs</u> | <u>NADH</u><br><u>Weighted costs</u> |
| biomass | -4.21 | -3.75 | -0.12 | -4.25 | -3.44 | -0.19 |
| isobutene | -40.68 | -48.50 | -3.08 | -41.84 | -48.62 | -4.35 |
| isoprene | -28.76 | -42.71 | -0.10 | -36.33 | -49.19 | -0.17 |
| ethylene | -35.16 | -36.41 | -4.13 | -35.25 | -30.85 | -5.89 |
| 1-undecene | -15.01 | -21.32 | -0.03 | -15.01 | -21.02 | -0.05 |

\*Weighted costs were calculated by multiplying the shadow prices of each cofactor metabolite (ATP, NADPH and NADH) with the its flux-sum. The shadow price values can be found in Supplementary Tables 7-8.

### Reaction-centric analysis

We were then interested to get a more complete picture of the cellular energetics. In order to identify and differentiate the metabolic reactions contributing to the computed turnover rates, we examined the intracellular flux distribution through central metabolic pathways. Using two condition-specific models (photoautotrophic and mixotrophic modes), we investigated the differences in the metabolic fluxes between the environmental conditions. In contract to CO_2_, glucose serves as both carbon and energy source. In addition, by comparing biomass (wild-type) and alkene-overproducing strains, potential reactions whose metabolic fluxes constrain end-product synthesis in *Synechocystis* can be identified. We took into account that the analysis is biased towards the maximization of an objective function, whilst the minimum growth rate was set to 10% of the maximum growth rate.

As mentioned above, the cellular control over the maintenance of redox homeostasis is partially facilitated by the activity of transhydrogenase enzyme (NADTRHD). However, the transfer of reducing equivalents from NAD(H) to NADP(H) is coupled to proton translocation across a membrane.^60^ Thus, the amount of NAD(P)H that can be produced by NADTRHD is limited by this energy requirement. We simulated two scenarios, in which an unconstrained bidirectional NADTRHD is present in the model or not. The model indicated infinite bounds for NADTRHD, computed by FVA (Supplementary Data 11-20). Similar observation was reported earlier, where the transhydrogenase reaction was poorly resolved by ^13^C measurements.^61^ Due to this, only the results from simulations lacking an active NADTRHD is presented; however, we discuss the difference in the flux distribution between active an inactive NADTRHD, where appropriate. At the same time, this allowed us to learn about how the entire metabolic network balances between NADP(H)/NAD(H).

#### Cofactor Energy Metabolism

From the estimated metabolic flux distributions, it can be realized that, under photoautotrophic conditions, the reactions contributing to ATP generation were within photosynthesis, respiration, glycolysis and pyridine metabolism (Table 4). The photosynthetic ATP synthetase (ATPSu) was the major reaction contributing to ATP generation, with over 90% contribution, followed by the pyruvate kinase (PYK) and the respiratory oxidative phosphorylation (ATPS4rpp_1). Notably, ethylene biosynthesis required a higher contribution of the ATPS4rpp_1 reaction with 1.9% influence, whereas isoprene required only 0.09%, and 0.13% was demanded for biomass biosynthesis. The computational solution also noticeably revealed that for all the objective functions, pyruvate is almost exclusively produced through the PYK reaction. Previous studies showed that malic enzyme (ME2) is the main route for synthesizing pyruvate in *Synechocystis*, under light conditions.^7,62^ The reason being that PYK is allosterically-inhibited by ATP.^63^ Fluxomic studies using labelled carbon^62,64^ pinpointed a potential bottleneck at the PYK reaction step. Such bottleneck was associated with a flux diversion into a three-step PYK bypass pathway, which involve phosphoenolpyruvate carboxylase (PPC), malate dehydrogenase (MDH) and ME2, and provide an alternate route for pyruvate formation. Accordingly, a subsequent overproduction of PYK in *Synechococcus elongatus* PCC 7942 led to a significant improvement in isobutyraldehyde production rate.^64^ A non-zero flux through the PPC and MDH reactions appeared in all but one (1-undecene) modeled strains (Fig. 2 and Supplementary Figs 1-9). Based on a pure stochiometric aspect, and in agreement with previous research, our results imply that PYK is a key enzyme in shaping cellular ATP and pyruvate levels in the cell. Further, in addition to PYK, PPC and MDH are good candidates for metabolic engineering of *Synechocystis* for hydrocarbon production.

**Table 4.**
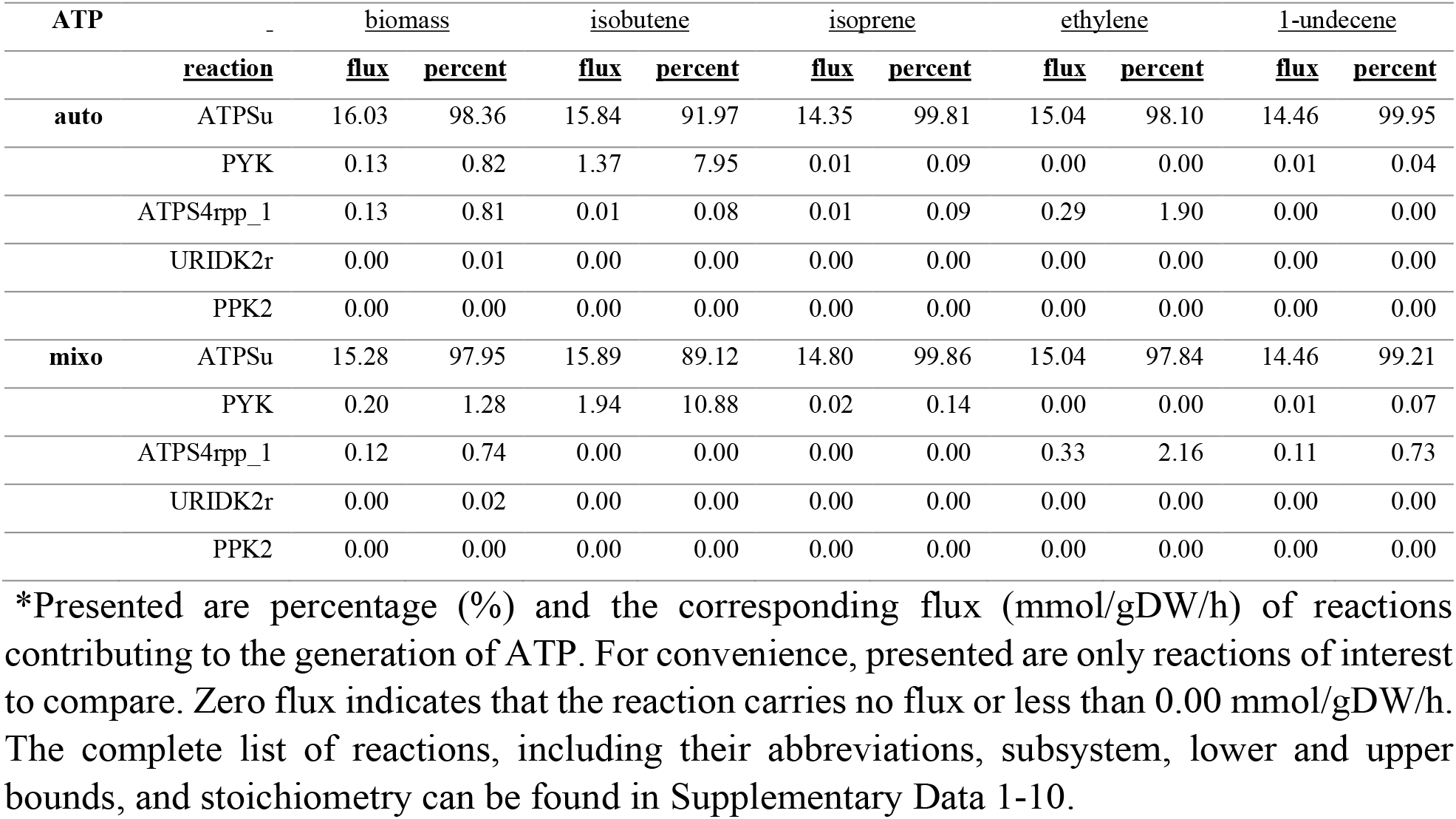
Predicted flux distributions of ATP-producing reactions in *Synechocystis* grown photoautotrophically and mixotrophically, when maximizing biomass and alkenes production.

| ATP | reaction | biomass |  | isobutene |  | isoprene |  | ethylene |  | 1-undecene |  |
| --- | --- | --- | --- | --- | --- | --- | --- | --- | --- | --- | --- |
|  |  | flux | percent | flux | percent | flux | percent | flux | percent | flux | percent |
| auto | ATPSu | 16.03 | 98.36 | 15.84 | 91.97 | 14.35 | 99.81 | 15.04 | 98.10 | 14.46 | 99.95 |
|  | PYK | 0.13 | 0.82 | 1.37 | 7.95 | 0.01 | 0.09 | 0.00 | 0.00 | 0.01 | 0.04 |
|  | ATPS4rpp_1 | 0.13 | 0.81 | 0.01 | 0.08 | 0.01 | 0.09 | 0.29 | 1.90 | 0.00 | 0.00 |
|  | URIDK2r | 0.00 | 0.01 | 0.00 | 0.00 | 0.00 | 0.00 | 0.00 | 0.00 | 0.00 | 0.00 |
|  | PPK2 | 0.00 | 0.00 | 0.00 | 0.00 | 0.00 | 0.00 | 0.00 | 0.00 | 0.00 | 0.00 |
| mixo | ATPSu | 15.28 | 97.95 | 15.89 | 89.12 | 14.80 | 99.86 | 15.04 | 97.84 | 14.46 | 99.21 |
|  | PYK | 0.20 | 1.28 | 1.94 | 10.88 | 0.02 | 0.14 | 0.00 | 0.00 | 0.01 | 0.07 |
|  | ATPS4rpp_1 | 0.12 | 0.74 | 0.00 | 0.00 | 0.00 | 0.00 | 0.33 | 2.16 | 0.11 | 0.73 |
|  | URIDK2r | 0.00 | 0.02 | 0.00 | 0.00 | 0.00 | 0.00 | 0.00 | 0.00 | 0.00 | 0.00 |
|  | PPK2 | 0.00 | 0.00 | 0.00 | 0.00 | 0.00 | 0.00 | 0.00 | 0.00 | 0.00 | 0.00 |

\*Presented are percentage (%) and the corresponding flux (mmol/gDW/h) of reactions contributing to the generation of ATP. For convenience, presented are only reactions of interest to compare. Zero flux indicates that the reaction carries no flux or less than 0.00 mmol/gDW/h. The complete list of reactions, including their abbreviations, subsystem, lower and upper bounds, and stoichiometry can be found in Supplementary Data 1-10.

**Fig. 2.**
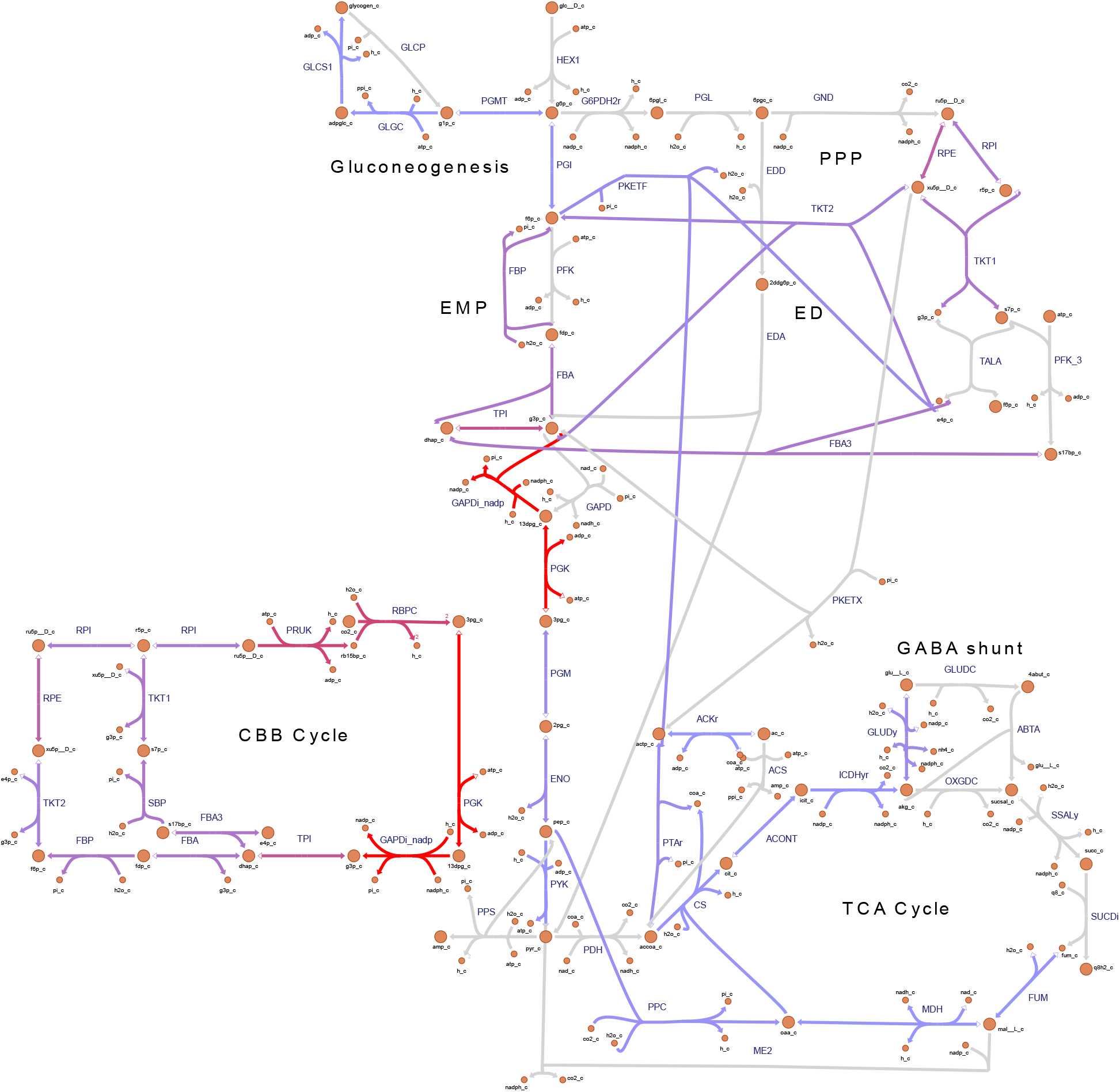
Metabolic flux map of central carbon metabolism for *Synechocystis* sp. PCC 6803 overproducing biomass, simulated to grow under photoautotrophic conditions. Reaction rates (mmol/gDW/h) were predicted using pFBA.^27^ Note that, the colors associated with the fluxes are relative to the other reactions rates presented in the map. Full arrows denote to consumption of a metabolite. Empty arrows denote to production of a metabolite. The map was generated with web-tool^49^. Metabolic reactions and metabolites (except heterologous ones) are indicated by their BiGG identifier.^30^

In all the modelled strains, ATP consumption was dictated by reactions within carbon fixation and glycolysis pathways (Table 5), in agreement with Young et al., 2011^62^. Under photoautotrophic conditions, when maximizing biomass accumulation, phosphoglycerate kinase (PGK), phosphoribulokinase (PRUK), and biomass formation reactions comprised 39.08%, 27.04% and 21.45% of total ATP-consuming reactions, respectively. When maximizing alkenes accumulation, the ATP consumption by PGK comprised 50.77%-58.81% and by PRUK 27.29%-33.52%, with the highest fluxes during isobutene production (Table 5).

**Table 5.**
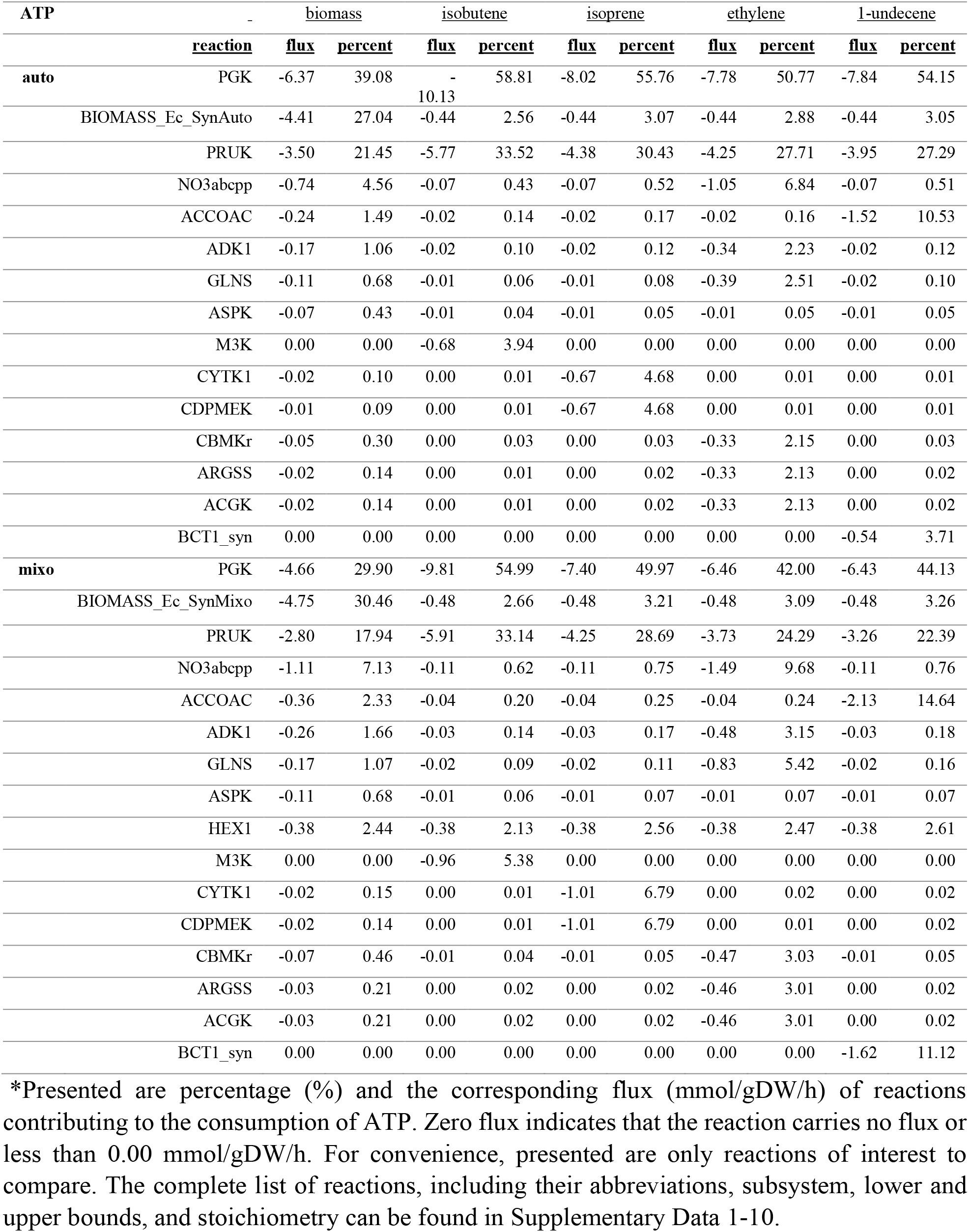
Predicted flux distributions of ATP-consuming reactions in *Synechocystis* grown photoautotrophically, when maximizing biomass and alkenes production.

Under mixotrophic conditions, the same trend as for phototrophic conditions was observed, where ATPSu contributed the most for ATP production, followed by PYK and ATPS4rpp. However, the contribution of PYK was shown to be higher. PYK contribution comprised 1.28%, 10.88%, 0.14% and 0.07% for biomass, isobutene, isoprene and 1-undecnece, respectively. In addition, ATPS4rpp_1 contribution was increases to 2.16% for ethylene biosynthesis. Regarding ATP consumption, the metabolic-flux levels of PGK were reduced to 29.90% for biomass formation, 54.99% for isobutene, 49.97% for isoprene, 42.00% for ethylene and 44.13% for 1-undecene synthesis. PRUK comprised 17.94% for biomass formation, 33.14% for isobutene, 28.69% for isoprene, 24.29% for ethylene and 22.39% for 1-undecene synthesis. Hexokinase (HEX1) comprised 2-2.5% for all objective functions. Nitrate transport (NO3abcpp) was mainly contributed to biomass and ethylene production, with up to 10% under both growth conditions (Tables 4 and 5).

PRUK and PGK control the ATP-dependent CO_2_ fixation and glycolysis/gluconeogenesis pathway, by converting of ribulose-5-phosphate (Ru5P) to ribulose-1,5-diphosphate (RuBP) and 3-phosphoglycerate (3-PGA) to 3-phosphoglycerol phosphate, respectively. Therefore, the enhancement of these enzymes could lead to improved photosynthetic capacity. Indeed, recent studies reported that overproduction of PRUK and PGK resulted in increased production of 2,3-butanediol in *S. elongatus* PCC 7942^9^ and ethylene in *Synechfocystis*.^65^ The increase in PRUK and PGK flux in the in-silico engineered strains is accompanied by additional consumption of ATP, which must be supplemented by ATP-producing reactions. Indeed, an engineering strategy was recently implemented in *S. elongatus* UTEX 2973, using an ATP-producing phosphoenolpyruvate carboxykinase (PPCK) and PRUK, which led to increased CO_2_ fixation rate and malate production.^66^ In direct connection to the shadow price and flux-sum analyses presented here, the computational analysis further pinpoints that, even though the photosynthetic machinery provides a virtually infinite supply of reductants and high-energy adenylates, these pools are rather limited to meet the ATP/NADPH requirement. ATP must be regenerated rapidly and utilized efficiently within the cell, in order to avoid cofactor deficiency and ensure an effective CO_2_-capturing system. Taken together, we suggest that a combinational metabolic engineering, based on a protein overproduction of the CBB enzymes (i.e., PRUK, PGK) and an ATP-producing enzymes (i.e., PYK), which together may form ATP cycling and fuel the CBB cycle, is beneficial for the heterologous production of alkenes, and probably other products, in *Synechocystis*.

The data also revealed specific flux distributions depending on the objective function. For example, the mevalonate-3-kinase (M3K) involved in the isobutene biosynthesis, cytidylate kinase (CYTK1) and 4-(cytidine 5’-diphospho)-2-C-methyl-D-erythritol kinase (CDPMEK), acting in the biosynthesis of isoprene, showed a high flux. In addition, for the accumulation of ethylene, carbamate kinase (CBMKr), argininosuccinate synthase (ARGSS) and N-acetylglutamate kinase (ACGK) presented high flux, probably due to interconnection with nitrogen metabolism. Bicarbonate transport (BCT1_syn) and acetyl-CoA carboxylase (ACCOAC) reactions exhibited a high flux for 1-undecene accumulation, relative to the other alkene biosynthesis pathways. This is probably due to the high amount of carbon assimilated during the elongation process of the fatty acyl chain.

The results presented here manifest the plasticity of certain metabolic pathways to sustain a steady mass flow of ATP against genetically or environmentally perturbed conditions. As revealed by the flux-sum analysis (Table 1), narrow ranges of intracellular ATP pool were observed, even against severe genetic perturbation, such as heterologous overproduction. We therefore investigated in what way and how of the precursors of ATP (ADP, AMP and Pi) are recycled in the metabolic network.

Overall, the reactions contributing to ADP-production and ADP-consumption mirrored those of ATP-consumption and ATP-production, respectively (Supplementary Tables 9-10). This mirroring was not shown for AMP flow. Reactions that dominated AMP production were phosphoribosylpyrophosphate synthetase (PRPPS), argininosuccinate synthase (ARGSS) and adenylsuccinate lyase (ADSL1r), comprising 64% in total (Supplementary Table 11). PRPPS functions at nexus of the pentose phosphate pathway and the de novo synthesis of nucleotides. ADSL1r participated in the salvage biosynthesis pathway for purines. Another key enzyme in buffering the adenylate pool is the adenylate kinase (ADK1) enzyme, which catalyze the interconversion between AMP and ATP with ADP (Fig. 3). The AMP-consuming ADK1 carried a similar flux across the isobutene, isoprene and 1-undecene-producing strains. A noticeable higher value was determined for ethylene and biomass production (Supplementary Table 12). Under mixotrophic conditions, ADK1 was shown to carry a higher flux, as compared to photoautotrophic conditions. Taken together we conclude that the salvage pathway via ADK1 contributes more to the ATP regeneration during biomass and ethylene overproduction, as compared to isobutene, isoprene and 1-undecene production. Enhanced ADK1 flux is presumably for providing more ADP and thereby sustain the increase in respiration rate, required for biomass and ethylene overproduction. An exception to these dominant reactions was observed for ethylene biosynthesis, where ARGSS comprised 95%. ARGSS co-produces AMP and N(omega)-(L-Arginino)succinate. N(omega)-(L-Arginino)succinate is used by ARGSL to generate L-arginine for ethylene overproduction. Hence, the surplus of AMP must be consumed by ADK1.

**Fig. 3.**
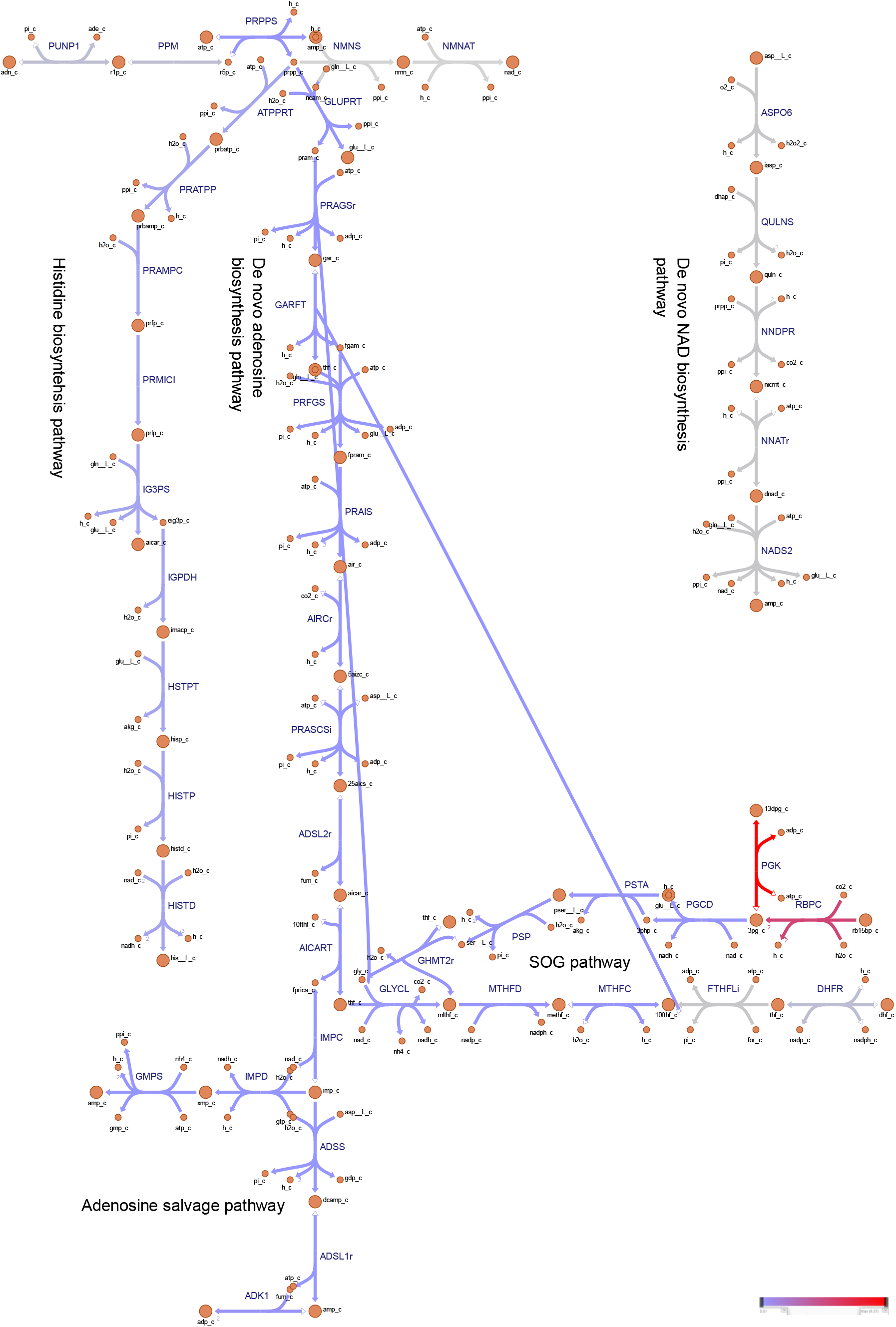
Metabolic flux map of auxiliary pathways enabling cellular energy and redox homeostasis. The auxiliary pathways are composed of the serine, one-carbon cycle, glycine synthesis (SOG) pathway and the biosynthesis of nucleotide precursors. *Synechocystis* sp. PCC 6803 was simulated to overproduce biomass, while grown under photoautotrophic conditions. Reaction rate (mmol/gDW/h) were predicted using pFBA.^27^ Note that, the colors associated with the fluxes are relative to the other reactions rates presented in the map. Full arrows denote to consumption of a metabolite. Empty arrows denote to production of a metabolite. The map was generated with web-tool^49^. Metabolic reactions and metabolites (except heterologous ones) are indicated by their BiGG identifier.^30^

Pi regeneration was dictated by GAPDi_nadp across strains, ranging from 38.39% in the biomass-producing strain to 61.11% in isobutene-producing strain (Supplementary Tables 13-14). Additionally, fructose-bisphosphatase (FBP) and sedoheptulose-bisphosphatase (SBP) significantly contributed to Pi production with 8-12%. Slightly lower values were observed for mixotrophic conditions. Phosphotransacetylase (PTAr) was found also to produce Pi in high extent, with 2% for biomass and 11.31% for 1-undecene production, under photoautotrophic conditions, and with higher flux under mixotrophic growth. The fructose-6-phosphate-utilizing phosphoketolase (PKETF) contributed substantially to Pi recycling, with 2.57%, 4.35%, 0.30%, 6.22% and 11.30% of the total Pi in the cell for biomass, isobutene, isoprene, ethylene and 1-undecene production, under photoautotrophic conditions; slightly higher values were observed under mixotrophic conditions. The xylulose-5-phosphate utilizing phosphoketolase (PKETX) appeared to be a key contributor of Pi for 1-undecene synthesis under mixotrophic conditions, comprising 3.65%. PKETF catalyzes the conversion of fructose-6-phosphate and Pi into acetyl phosphate and D-Erythrose 4-phosphate. Acetyl phosphate, can subsequently be converted into acetyl-CoA by PTAr, or alternatively catalyzed by acetate kinase (ACKr) to form acetate and ATP. Acetate, is, in turn, converted into acetyl-CoA by acetyl-CoA synthetase (ACS). However, the computational modelling reaved that instead, the bidirectional ACKr generates acetyl phosphate from the L-homoserine node (Supplementary Data 21-30). D-erythrose 4-phosphate can be recycled into the pentose phosphate pathways via transketolase and transaldolase enzymes, or, alternatively, into pyruvate via glycolysis. The product of PKETX, glyceraldehyde 3-phosphate (G3P) can be metabolized into the gluconeogenesis and glycolysis pathways. Similar evidence concerning this fine-tuned network for Pi was manifested in *S. elongatus* PCC 7942.^67^ Additionally, product-specific reaction rates that contributed to the flow of Pi were identified. For example, inorganic diphosphatase (PPA) was shown to have a high contribution for isoprene biosynthesis, by pulling out diphosphate – the byproduct of isoprene synthase.

In summary, the stoichiometric analysis implies that acetate metabolism via phosphoketolase serves as efficient a carbon- and energy-recycling pathway for cell maintenance, as previously proposed^34^. In addition, FBP, SBP and ADK1 play a key role in the recycling of Pi and ADP/AMP, and thereby determining the ATP/NADPH ratio. Our results suggest that the ATP/ADP ratio are mainly controlled by the de novo synthesis, whereas for ethylene and biomass accumulation, the salvage pathway to recover nucleotides is used (Supplementary Figs 17-18). This analysis demonstrated the importance of the flexibility in energy- and carbon-converting enzymes, in order to achieve the required ATP/NADPH output ratio.^68,69^ It also emphasizes the essentiality of metabolic shortcuts in central carbon metabolism for attaining effective metabolic states.^70^

#### Cofactor Redox Metabolism

We then analyzed the regeneration of the reducing equivalents NADPH and NADH (Table 6). The theoretical flux analysis of NADPH metabolism revealed that the photosynthetic ferredoxin-NADP^+^ reductase (FNOR) contributed largely to the supply of NADPH, comprising 97.80%, 99.83%, 97.98% and 88.09% of FNOR activity for biomass, isoprene, isobutene and ethylene, respectively. Further, isocitrate dehydrogenase (ICDHyr) in the TCA cycle contributed 1.33% of NADPH generation for biomass synthesis, 11.56% for ethylene and 1.22% for isobutene, whereas only 0.10% for isoprene production. These differences could be explained by the fact that the acetyl-CoA is diverted from the TCA cycle to the synthesis of ethylene. Of note, the flux of ICDHyr, within the oxidative branch of the TCA cycle, in the ethylene-overproducing strain appeared to be significantly higher than that in the wild-type, which is considered to be rigid.^31,71^ The computed ICDHyr flux for the ethylene-overproducing strain was 9.8-fold of that in the wild-type. Similar trends were seen in an in-vivo engineered strain,^72^ in which the ICDHyr showed a 3.8-fold flux in ethylene-producing strain, as compared to wild-type values. The production of ICDHyr caused an enhanced substrate supply and ethylene productivity in *Escherichia coli*.^73^ An increased ethylene production was also documented in *Synechocystis* overexpressing *ppc*.^74^ Taken together, our findings demonstrate that PCC ICDHyr are target candidates for enhancing production of ethylene and possibly other metabolites that stem from the oxidative portion of the TCA cycle.

**Table 6.** Predicted flux distributions of NADPH-producing reactions in *Synechocystis* grown photoautotrophically and mixotrophically, when maximizing biomass and alkenes production.

| NADPH |  | biomass |  | isobutene |  | isoprene |  | ethylene |  | 1-undecene |  |
| --- | --- | --- | --- | --- | --- | --- | --- | --- | --- | --- | --- |
|  | Reaction | flux | percent | flux | percent | flux | percent | flux | percent | flux | percent |
| auto | FNOR | 7.56 | 97.66 | 10.92 | 99.83 | 10.15 | 99.82 | 7.50 | 88.08 | 10.94 | 99.77 |
|  | ICDHyr | 0.10 | 1.33 | 0.01 | 0.09 | 0.01 | 0.10 | 0.98 | 11.56 | 0.01 | 0.09 |
|  | MTHFD | 0.06 | 0.78 | 0.01 | 0.06 | 0.01 | 0.06 | 0.01 | 0.07 | 0.01 | 0.05 |
|  | ME2 | 0.00 | 0.00 | 0.00 | 0.00 | 0.00 | 0.00 | 0.02 | 0.27 | 0.01 | 0.06 |
| mixo | FNOR | 6.44 | 95.96 | 11.00 | 99.75 | 10.65 | 99.75 | 5.79 | 80.28 | 10.79 | 99.65 |
|  | ICDHyr | 0.15 | 2.29 | 0.02 | 0.14 | 0.02 | 0.14 | 1.39 | 19.28 | 0.02 | 0.14 |
|  | MTHFD | 0.09 | 1.34 | 0.01 | 0.08 | 0.01 | 0.08 | 0.01 | 0.13 | 0.01 | 0.08 |
|  | ME2 | 0.00 | 0.00 | 0.00 | 0.00 | 0.00 | 0.00 | 0.02 | 0.28 | 0.01 | 0.10 |

\*Presented are percentage (%) and the corresponding flux (mmol/gDW/h) of reactions contributing to the generation of NADPH. Zero flux indicates that the reaction carries no flux or less than 0.00 mmol/gDW/h. For convenience, presented are only reactions of interest to compare. The complete list of reactions, including their abbreviations, subsystem, lower and upper bounds, and stoichiometry can be found in Supplementary Data 1-10.

Our computational analysis predicted that malic enzyme (ME2) is dispensable, under photoautotrophic conditions. An exception was observed for ethylene and 1-undecene synthesis, for which ME2 showed a non-zero flux of 0.02 and 0.01 mmol/gDW/h, respectively. This was surprising since ME2 converts malate and NADP^+^ into pyruvate and NADPH, while releasing one molecule of CO_2_ in the process. While ME2 flux contributed with 97% and 30% to pyruvate production in the ethylene- and 1-undecene-producing strains under photoautotrophy, the contribution under mixotrophy was 57% and 30% in these strains.

It has been hypothesized that ME2 serves a role in providing intracellular CO_2_ for ribulose 1,5-bisphosphate carboxylase/oxygenase (RuBisCO), in a C4-like pathway.^75^ Considering that the three-step pathway forms part of the reductive TCA cycle, which exhibits a higher flux than the oxidative portion, we thus propose that that the so-called three-step bypass is not necessarily a “bottleneck”, as assumed.^62,64^ The CO_2_ released by ME2 can be further assimilated by PPC, thus resulting in a net zero carbon loss. A direct correlation between the activity of ME2 and the extent of lipid accumulation has been observed in heterotrophs.^76–78^ Since ME2 co-produces CO_2_ and NADPH, ME2 can further act to meet the stoichiometric demands for carbon and redox consumed during fatty acids and ethylene synthesis.

The flux distribution of reactions catalyzing NADPH consumption were investigated next (Table 7). We noted that the anabolic glyceraldehyde-3-phosphate dehydrogenase (GAPDi_nadp) was shown to carry a very high flux of 6.37-10.13 mmol/gDW/h, which corresponds to 82% -93% of total fluxes in the analyzed strains (Table 7, Fig. 2). In addition, a portion of carbon flux was diverted to glycogen synthesis glucose-1-phosphate adenylyltransferase (GLGC) and glycogen synthase (GLCS1) reactions. This suggests that carbon is channeled back to the Embden-Meyerhof-Parnas (EMP) pathway. In *Synechocystis*, inactivation of GLGC led to an increased NADPH pool, and the glycogen metabolism has shown to act as a ATP-buffering system.^79,80^ It has furthermore been reported that glycolytic routes form anaplerotic shunts that replenish metabolites for carbon fixation.^81^ Since the glycolysis/gluconeogenesis pathways share reactions with the CBB cycle, we postulate that, in general, gluconeogenesis functions to maintain energy end redox homeostasis in *Synechocystis*, while simultaneously regenerating substrates to the CBB cycle.

**Table 7.**
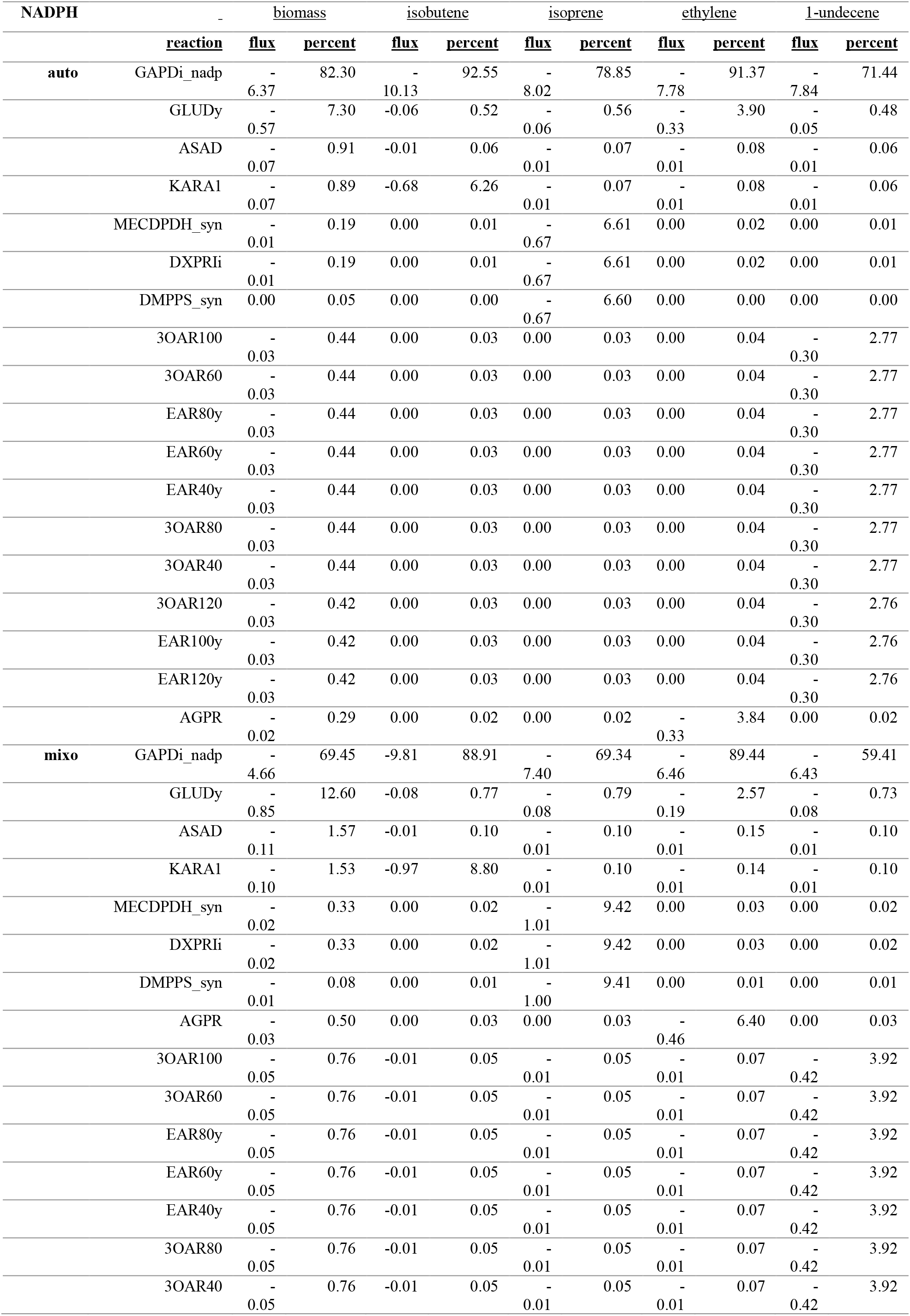
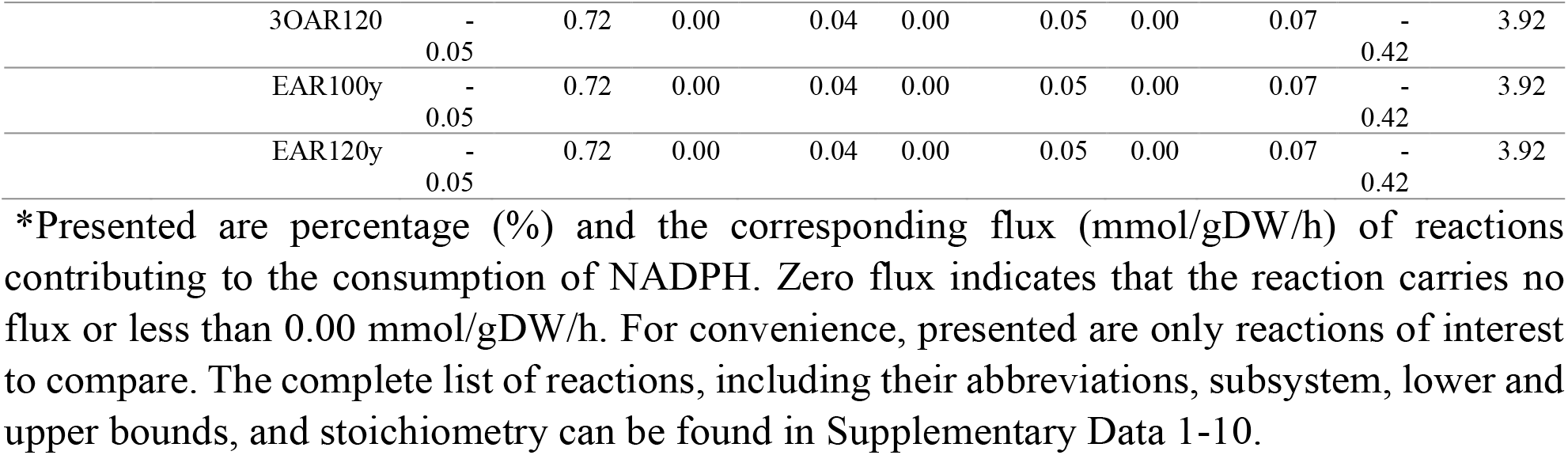
Predicted flux distributions of NADPH-consuming reactions in *Synechocystis* grown photoautotrophically and mixotrophically, when maximizing biomass and alkenes production.

Except for the general importance of GAPDi_nadp, the data revealed other major contributor to NADPH consumption which are more strain and condition specific. In biomass and ethylene-producing strains, L-glutamate dehydrogenase (GLUDy) was comprising 7.30% and 3.90%, respectively. During isobutene overproduction, a high flux was observed for the ketol-acid reductoisomerase (2,3-dihydroxy-3-methylbutanoate) (KARA1), which is involved in branched chain amino acid (BCAA) biosynthesis. A high flux was determined for 3-oxoacyl-[acyl-carrier-protein] reductase (n-C10:0) (3OAR100) during accumulation of 1-undecene, and for MECDPDH_syn, DXPRIi and DMPPS_syn during isoprene biosynthesis. All these reactions are tightly associated with the synthesis of specific alkenes.

In addition, during ethylene production, the N-acetyl-g-glutamyl-phosphate reductase (AGPR) reaction contributed 3.84% and 6.40% for NADPH utilization. AGPR proceeds N-acetylglutamate synthase (ACGS) and the ATP-consuming N-acetylglutamate kinase (ACGK) to from N-acetyl-L-glutamate 5-semialdehyde. N-acetyl-L-glutamate 5-semialdehyde can then be converted back to N-acetyl-L-glutamate via N-acetyl-L-glutamate (ACOTA) and ornithine transacetylase (ORNTAC), while generating 2-oxoglutarate in the process. Alternatively, ORNTAC acts in the reverse direction, consuming N-acetyl-L-glutamate to produce ornithine. Ornithine, in turn, is fed by ornithine carbamoyltransferase (OCBT) to eventually regenerate L-arginine (Fig. 4A, Supplementary Fig. 11). In summary, the analysis identified additional routes for efficient ethylene production, which is interlinked with the urea cycle and L-arginine metabolism, to provide both arginine and 2-oxoglutarate for ethylene-forming enzyme (EFE).

**Fig. 4.**
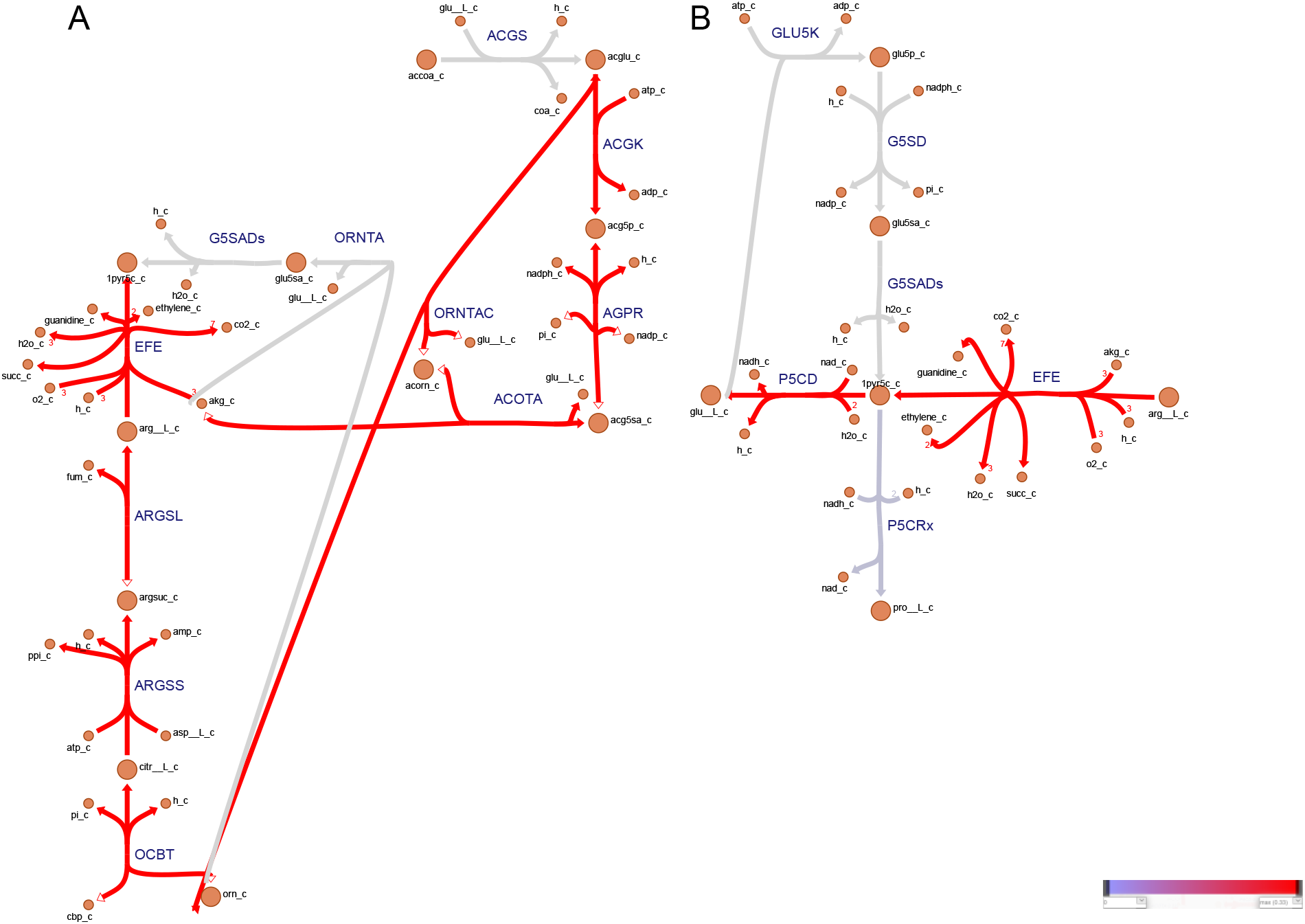
Metabolic flux map through reactions involved in balancing nitrogen metabolism for *Synechocystis* sp. PCC 6803 overproducing ethylene, simulated to grow under photoautotrophic conditions. (A) L-glutamate and L-arginine regeneration through the urea cycle. (B) L-glutamate and L-proline regeneration. Reaction rates (mmol/gDW/h) were predicted using pFBA.^27^ Note that, the colors associated with the fluxes are relative to the other reactions rates presented in the map. Full arrows denote to consumption of a metabolite. Empty arrows denote to production of a metabolite. The map was generated with web-tool^49^. Metabolic reactions and metabolites (except heterologous ones) are indicated by their BiGG identifier.^30^

We noticed that the reductase enzymes (eg. 3OAR100, 3-oxoacyl-[acyl-carrier-protein] reductase (n-C10:0)) along the 1-undecene biosynthesis pathway carried a flux of 0.30 mmol/gDW/h, whereas the DXPRIi (1-deoxy-D-xylulose reductoisomerase) in the isoprene pathway carries a higher flux of 0.67 mmol/gDW/h. These data suggest that, the turnover rate of NAD(P)H is not governed by the length of the metabolic pathway, nor by the actual number of moles shared in a particular pathway, but rather by the capability of individual reactions within the metabolic network to carry a flux.

Provision of the reducing agent NADH was also probed (Table 8). Under both growth conditions, phosphoglycerate dehydrogenase (PGCD) showed the highest contribution to NADH generation in biomass, isoprene and 1-undecene production. For isobutene and ethylene production, malate dehydrogenase (MDH) and 3-isopropylmalate dehydrogenase (IPMD) were predicted to have the most contribution for NADH generation, comprising 97.13% and 65.07%, respectively. In addition, Δ1-pyrroline-5-carboxylate dehydrogenase (P5CD, 33.14%), which acts in the L-arginine and L-proline metabolism, was found to be predominant during ethylene overproduction. Enzymatic reactions participating in the biosynthesis of histidine and purine nucleotides were also adding to NADH generation, histidinol dehydrogenase (HISTD), inosine-5′-monophosphate (IMP) dehydrogenase (IMPD) and glycine cleavage system (GLYCL). HISTD, IMPD and GLYCL reactions stem from the same precursor 5-phospho-alpha-D-ribose 1-diphosphate (PPRP), a pentose phosphate pathway derivative. PRPP is also a common intermediate in the tryptophan biosynthesis branch, which requires PRPP for the production of N-phosphoribosyl anthranilate from anthranilate.

**Table 8.** Predicted flux distributions of NADH-producing reactions in *Synechocystis* grown photoautotrophically and mixotrophically, when maximizing biomass and alkenes production.

| NADH |  | biomass |  | isobutene |  | isoprene |  | ethylene |  | 1-undecene |  |
| --- | --- | --- | --- | --- | --- | --- | --- | --- | --- | --- | --- |
|  | reaction | flux | percent | flux | percent | flux | percent | flux | percent | flux | percent |
| auto | PGCD | 0.10 | 38.97 | 0.01 | 1.36 | 0.01 | 38.98 | 0.01 | 0.98 | 0.01 | 54.53 |
|  | MDH | 0.07 | 28.54 | 0.01 | 0.99 | 0.01 | 28.54 | 0.63 | 65.07 | 0.00 | 0.00 |
|  | IPMD | 0.04 | 17.67 | 0.68 | 97.13 | 0.00 | 17.67 | 0.00 | 0.44 | 0.00 | 24.73 |
|  | HISTD | 0.01 | 5.77 | 0.00 | 0.20 | 0.00 | 5.77 | 0.00 | 0.14 | 0.00 | 8.07 |
|  | IMPD | 0.01 | 5.32 | 0.00 | 0.19 | 0.00 | 5.32 | 0.00 | 0.13 | 0.00 | 7.44 |
|  | GLYCL | 0.01 | 3.68 | 0.00 | 0.13 | 0.00 | 3.66 | 0.00 | 0.09 | 0.00 | 5.14 |
|  | P5CD | 0.00 | 0.00 | 0.00 | 0.00 | 0.00 | 0.00 | 0.32 | 33.14 | 0.00 | 0.00 |
| mixo | PGCD | 0.14 | 38.97 | 0.01 | 1.43 | 0.01 | 38.97 | 0.01 | 1.03 | 0.01 | 54.55 |
|  | MDH | 0.10 | 28.54 | 0.01 | 1.05 | 0.01 | 28.54 | 0.91 | 65.32 | 0.00 | 0.00 |
|  | IPMD | 0.06 | 17.67 | 0.97 | 96.98 | 0.01 | 17.67 | 0.01 | 0.47 | 0.01 | 24.73 |
|  | HISTD | 0.02 | 5.77 | 0.00 | 0.21 | 0.00 | 5.77 | 0.00 | 0.15 | 0.00 | 8.07 |
|  | IMPD | 0.02 | 5.32 | 0.00 | 0.20 | 0.00 | 5.32 | 0.00 | 0.14 | 0.00 | 7.44 |
|  | GLYCL | 0.01 | 3.68 | 0.00 | 0.14 | 0.00 | 3.68 | 0.00 | 0.10 | 0.00 | 5.12 |
|  | P5CD | 0.00 | 0.00 | 0.00 | 0.00 | 0.00 | 0.00 | 0.46 | 32.80 | 0.00 | 0.00 |

\*Presented are percentage (%) and the corresponding flux (mmol/gDW/h) of reactions contributing to the generation of NADH. Zero flux indicates that the reaction carries no flux or less than 0.00 mmol/gDW/h. For convenience presented are only reactions of interest to compare. The complete list of reactions, including their abbreviations, subsystem, lower and upper bounds, and stoichiometry can be found in Supplementary Data 1-10.

In this regard, it should be noted that, since MDH showed to act in the reverse reaction, from L-malate into oxaloacetate, L-malate is originated from fumarase (FUM), for which fumarate can be generated by either adenylsuccinate lyase (ADSL1r) or argininosuccinate lyase (ARGSL). Oxaloacetate is then fueled into the oxidative branch of the TCA cycle by citrate synthase (CS). These metabolic pathways serve a role as a “push-pull” effect to promote carbon distribution towards coenzyme A and enhanced ethylene and 1-undecene synthesis. Interestingly, and less intuitively, the IPMD, a known key enzyme participating in the BCAA biosynthesis pathway, also showed a high contribution for isoprene and 1-undecene biosynthesis. We presume that the degradation of BCAA provides an alternative route for the synthesis of acetyl-CoA - a precursor molecule for terpenoids and lipids in *Synechocystis*. The highly NADH-contributing enzymatic reaction identified in our study could be useful as engineering targets for production under light conditions. Our analysis also strengthens the view that bidirectional reaction steps (e.g., PGK, PGM and MDH) are central to the metabolism of cyanobacteria, capable of operating in different trophic modes of life.

We then explored the flux distribution of the NADH-consuming reactions (Table 9). We observed that, NAD(P)H dehydrogenase (NDH1_2u) in the oxidative phosphorylation had the highest contribution for all but one (1-undecene) investigated strains, and for both trophic modes. NDH1_2u comprised 44.84% for biomass and isoprene, 98.08% for isobutene, and 60.20% of total NADH consumption for ethylene accumulation. We were curious to compare the flux distributions with and without active NADTRHD. The flux distributions in the model including the transhydrogenase reaction (NADTRHD) mirrored those without the bidirectional NADTRHD, with the exception that NADH was then oxidized by NADTRHD (Supplementary Data 11-20). We observed that primary usage of NADH was by NADTRHD across all strains and trophic modes NADTRHD comprised 44.85% for biomass, isoprene and 1-undecene in total, whereas over 98% for isobutene and ethylene (Supplementary Data 11-20). The significance of NADTRHD for increased production of bioproducts has been demonstrated by overexpression of *pntAB* (encoding NADTRHD), which resulted in increased production of 3-hydroxypropionic acid^82^ and sorbitol,^83^ respectively. Thus, the overexpression of the *pntAB* could be beneficial for the production of specific alkene compounds (e.g., isobutene and ethylene).

**Table 9.** Predicted flux distributions of NADH-consuming reactions in *Synechocystis* grown photoautotrophically and mixotrophically, when maximizing biomass and alkenes production.

| NADH |  | biomass |  | isobutene |  | isoprene |  | ethylene |  | 1-undecene |  |
| --- | --- | --- | --- | --- | --- | --- | --- | --- | --- | --- | --- |
|  | reaction | flux | percent | flux | percent | flux | percent | flux | percent | flux | percent |
| auto | NDH1_2u | -0.11 | 44.84 | -0.69 | 98.08 | -0.01 | 44.84 | -0.59 | 60.20 | 0.00 | 0.00 |
|  | HSDxi | -0.05 | 21.52 | -0.01 | 0.75 | -0.01 | 21.52 | -0.01 | 0.54 | -0.01 | 30.11 |
|  | ALAD_L | -0.04 | 15.73 | 0.00 | 0.55 | 0.00 | 15.70 | 0.00 | 0.40 | 0.00 | 22.01 |
|  | P5CRx | -0.02 | 7.95 | 0.00 | 0.28 | 0.00 | 7.95 | 0.00 | 0.20 | 0.00 | 11.12 |
|  | G3PD1ir | -0.01 | 5.83 | 0.00 | 0.20 | 0.00 | 5.82 | 0.00 | 0.15 | 0.00 | 8.16 |
|  | MTHFR2 | -0.01 | 4.13 | 0.00 | 0.14 | 0.00 | 4.13 | 0.00 | 0.10 | 0.00 | 5.78 |
|  | GLUSx | 0.00 | 0.00 | 0.00 | 0.00 | 0.00 | 0.00 | -0.37 | 38.42 | 0.00 | 22.81 |
| mixo | NDH1_2u | -0.16 | 44.84 | -0.98 | 97.97 | -0.02 | 44.84 | -0.55 | 39.81 | 0.00 | 0.00 |
|  | HSDxi | -0.08 | 21.52 | -0.01 | 0.79 | -0.01 | 21.52 | -0.01 | 0.57 | -0.01 | 30.11 |
|  | ALAD_L | -0.06 | 15.73 | -0.01 | 0.58 | -0.01 | 15.73 | -0.01 | 0.41 | -0.01 | 21.97 |
|  | P5CRx | -0.03 | 7.95 | 0.00 | 0.29 | 0.00 | 7.95 | 0.00 | 0.21 | 0.00 | 11.12 |
|  | G3PD1ir | -0.02 | 5.83 | 0.00 | 0.21 | 0.00 | 5.83 | 0.00 | 0.15 | 0.00 | 8.14 |
|  | MTHFR2 | -0.02 | 4.13 | 0.00 | 0.15 | 0.00 | 4.13 | 0.00 | 0.11 | 0.00 | 5.78 |
|  | GLUSx | 0.00 | 0.00 | 0.00 | 0.00 | 0.00 | 0.00 | -0.82 | 58.74 | -0.01 | 22.81 |

\*Presented are percentage (%) and the corresponding flux (mmol/gDW/h) of reactions contributing to the consumption of NADH. Zero flux indicates that the reaction carries no flux or less than 0.00 mmol/gDW/h. For convenience, presented are only reactions of interest to compare. The complete list of reactions, including their abbreviations, subsystem, lower and upper bounds, and stoichiometry can be found in Supplementary Data 1-10.

Additional reaction rates that contributed to the flow of NADH during isoprene biosynthesis were L-homoserine dehydrogenase (HSDxi, 21.52%), L-alanine dehydrogenase (ALAD_L, 15.73%), Δ1-pyrroline-5-carboxylate reductase (P5CRx, 7.95%), glycerol-3-phosphate dehydrogenase (G3PD1ir, 5.83%) and 5,10-methylenetetrahydrofolate reductase (MTHFR2, 4.13%). Similarly, HSDxi (30.11%), followed by ALAD_L (22.01%), P5CRx (11.12%) and MTHFR2 (5.78%) were dominant for 1-undecene synthesis.

A two-step process, including L-aspartate kinase (ASPK) and aspartate-semialdehyde dehydrogenase (ASAD), results in the generation of L-aspartate 4-semialdehyde from L-aspartate. L-aspartate 4-semialdehyde is then converted into L-homoserine by HSDxi. This biosynthesis pathway competes with the TCA cycle for carbon flux through a shared intermediate, oxaloacetate. Therefore, the overexpression of genes in the L-homoserine and L-threonine biosynthesis pathways could result in higher carbon flow through the TCA cycle. This, in turn, enhances the coenzyme A (CoA) availability, generated by citrate synthase (CS). CoA could be subsequently utilized by PTAr for generating acetyl-CoA (Supplementary Data 1-10, Supplementary Data 21-30).

The canonical L-proline biosynthesis pathway encompasses the conversion of L-glutamate into L-proline by sequential catalyzation steps, involving L-glutamate 5-kinase (GLU5K), L-glutamate-5-semialdehyde dehydrogenase (G5SD), L-glutamate 5-semialdehyde dehydratase (G5SADs) and Δ1-pyrroline-5-carboxylate reductase (P5CRx). For ethylene overproduction, the flux analysis showed that the product of the G5SADs reaction - Δ1-pyrroline-5-carboxylate - undergoes either a reduction by Δ1-pyrroline-5-carboxylate dehydrogenase (P5CD) to regenerate L-glutamate or converted into L-proline (Fig. 4B).

The model also predicted an increase in flux for glutamine:2-oxoglutarate aminotransferase (GOGAT), known as L-glutamate synthase (GLUSx in the model) reaction. GLUSx catalyzes the formation of L-glutamate from glutamine and 2-oxoglutarate, and is an important enzyme in ammonia assimilation. L-glutamate can be converted into L-proline by Δ1-pyrroline-5-carboxylate reductase (P5CRx). The upregulation of the P5CRx biosynthetic gene was determined during TAG accumulation in the oleaginous fungus *Mortierella alpina*^84^ and the diatom *Phaeodactylum tricornutum*.^85^ In plants, the degradation of L-proline has been considered to serve as a redox shuttle.^86^ Previous work has attributed the increased GLUSx to support lipogenesis in tumor cells.^87^ Interestingly, the model revealed that by unconstraining the NADTRHD, the GLUSx flux is diminished and the consumption of NADH is replaced by NADTRHD. Taken together, our results suggest that, besides adaptation to a stressful environment, nitrogen metabolism via L-glutamate and L-proline metabolites contributes stoichiometrically for balancing redox potential in a plant-like manner, during fatty acids and ethylene overproduction.

Our data additionally showed that *Synechocystis* exhibits a residual respiratory activity under light conditions, which is agreement with earlier studies.^40^ Cyanobacteria have the capacity to utilize both NADPH and NADH as electron donor for the respiratory electron transport chain.^88^ Low respiration rate has been identified as a possible limiting factor of ethylene production via EFE in Saccharomyces cerevisiae,^89^ and could explain the low ethylene production found experimentally in *Synechocystis*.^44^ Hence, it can be speculated that an enhanced respiration capacity could be beneficial for ethylene bioproduction. Re-oxidizing the surplus of NADH generated by MDH in the TCA cycle and P5CD, as well as converting NAD(P)H to ATP via oxidative phosphorylation, could be valid means to meet the demanded ATP/NADPH ratio. In addition, the ATP consumed by the CBB cycle could be coupled to the NAD(P)H consumption and ATP production by the respiratory pathways.

L-alanine dehydrogenase (ALAD_L) recycles L-alanine using ammonium and NADH, while releasing NAD^+^ as redox cofactor. In a recent study,^90^ the introduction of ALAD_L for nitrogen assimilation, instead of the native ferredoxin-dependent nitrite/nitrate reductases, resulted in increased branched-chain alcohol production. This facilitates the regeneration of the NAD(H) pool in the system, where NADH-producing reactions carry a high flux. However, such a high flux of ALAD_L was not observed for isobutene or ethylene overproduction. This is probably due the fact that the current computational analysis did not consider a growth-coupled engineering strategy or reaction knockouts. In the filamentous cyanobacterium *Anabaena* sp. PCC 7120, a stain deficient in active ALAD_L exhibited an impaired rate of diazotrophic growth.^91^ Therefore, we conclude that the high flux determined for ALAD_L is a consequence of the need for NAD(H) and nitrogen regeneration.

Glycerol-3-phosphate dehydrogenase (G3PD1ir) exhibited a relatively higher flux for 1-undecene production, than other biosynthesis pathways. This is in accordance with G3PD1ir being involved in the de novo synthesis of glycerolipids, catalyzing the redox conversion of dihydroxyacetone phosphate (DHAP) into glycerol 3-phosphate. A recent work showed that lipid biosynthesis imposes a substantial NAD^+^ consumption in proliferating cancer cells^87^.

An important aspect of balancing the redox cofactors in the cell is the phosphorylation of NAD^+^ to NADP^+^ by NAD kinase (NADK),^14^ and the reversal dephosphorylation of NADP^+^ into NAD^+^ by NADP phosphatase. NADP^+^ serves as the final electron acceptor from photosystem I to form NADPH. Hence, it was surprising to observe that this reaction had a negligible contribution to the overall flux (Supplementary Data 1-10). The role of NADK only begins to be elucidated,^92,93^ and the functionality of NADP^+^ phosphatase remains to be discovered, which are not covered in the model.

#### Serine and glycine metabolism represents a novel pathway for cellular energy and redox homeostasis in Synechocystis

The reaction-centric analysis uncovered that, reactions participating in L-serine synthesis, one-carbon metabolism and the glycine cleavage system - SOG pathway - contribute to the energy requirements of the cell. In this pathway, 3-PGA, branching from glycolysis and the CBB, is successively catalyzed by PGCD, phosphoserine aminotransferase (PSTA) and phosphoserine phosphatase (PSP) to form L-Serine, through the intermediate 3-phosphohydroxypyruvate. 3-phosphohydroxypyruvate is trans-aminated by phosphoserine aminotransferase to from O-phospho-L-serine. L-glutamate serves as the nitrogen donor for this reaction. L-glutamate can by supplied by L-glutamate dehydrogenase (GLUDy) and L-glutamate synthase (GLUSx). L-serine is catalyzed by glycine hydroxymethyltransferase (GHMT2r) into glycine, which is then used by the glycine cleavage system (GLYCL) to from 5,10-methylenetetrahydrofolate. NADPH can be produced when methylenetetrahydrofolate is oxidized to 5,10-methenyltetrahydrofolate and 10-formyltetrahydrofolate, by methylenetetrahydrofolate dehydrogenase (MTHFD) and methenyltetrahydrofolate cyclohydrolase, respectively. Additionally, 10-formyltetrahydrofolate, together with glycine, are used for the de novo biosynthesis of purines (i.e., ATP) (Fig. 3). These non-zero fluxes appeared when maximizing the production of alkenes, and were further emphasized when maximizing biomass production (Fig. 3 and Supplementary Figs 12-20). Previous work revealed that cancer cells can rewire their metabolism to rely on SOG pathway for regeneration of redox agents.^94–96^ The gene encoding folate reductase, which converts folate into 7,8-dihydrofolate, is not annotated in the genome of *Synechocystis* sp. PCC 6803, in contrast to the phototrophs *P. tricornutum* and *C. reinhardtii*. However, the knowledge on *Synechocystis* metabolism is yet incomplete,^97^ and it is recognized that this microorganism contains enzymes with similarity to folate transporters.^98^ Taken together, it is plausible that the SOG pathway can function in *Synechocystis* as an auxiliary pathway to regenerate cofactors such as ATP, NADPH and NADH.

## Conclusions

We applied a comprehensive computational fluxomic analysis to study the systemic properties of *Synechocystis* sp. PCC 6803, able to simultaneously utilize organic carbons and CO_2_. This work is the first to report the use of combined modelling algorithms, including pFBA, FVA and flux-sum, to study the intracellular energy and redox balance in *Synechocystis*. Balancing ATP and NAD(P)H metabolism is important to achieve efficient and robust cyanobacterial strains and high level of production; however, often neglected in metabolic engineering strategies. The comparative genome-scale modelling revealed that different routes or flux extent enable the balancing mechanism and avoidance of excessive energy during overproduction of end-products. Through a system-wide analysis, we were able to identify to which extent each of the cofactors are demanded and imposed the most metabolic burden on the cell. We could also recognize to which extent each the cofactor-dependent reactions contribute to the observed phenotypes. It was apparent that the overall activity of the *Synechocystis* metabolism is governed by several high-flux core reactions that involve both carbon and cofactor regeneration (e.g. PGK, GAPDi_nadp). This probably confers metabolic stability and allows the cell to adapt rapidly to changes in growth conditions. We conclude that the assessment of ATP and NAD(P)H balance can neither be done in isolation from each other and their precursors, nor from carbon and nitrogen metabolism. Even though the predictions presented in this study are yet to be validated experimentally, the new knowledge generated in this work serves as a direction for future research and has potential implications for metabolic engineering. While our analysis used alkenes as products of choice, we are here providing hints for strain design strategies for the efficient bioproduction of other valuable compounds, deriving from the core metabolic pathways in *Synechocystis*.

## Data availability

The authors declare that all the data supporting the work are available within the paper, its supplementary information, and in the GitHub repository https://github.com/amitkugler/CBA.

## Code availability

The models and code used for this study can be found in the GitHub repository https://github.com/amitkugler/CBA.

## Acknowledgments

This work was supported by Formas—A Swedish Research Council for Sustainable Development” (project no. 2021-01669), Swedish Energy Agency (project no. 44728-1) and the NordForsk Nordic Center of Excellence ‘NordAqua’ (project no. 82845). The authors would like to thank Kiyan Shabestary and Pia Lindberg for valuable discussions.

## CRediT author contributions

AK: Conceptualization, Methodology, Software, Data curation, Investigation, Formal analysis, Validation, Visualization, Writing-Original Draft, Writing-Review & Editing. KS: Conceptualization, Supervision, Funding Acquisition, Writing-Review & Editing.

## References

1. Baebprasert, W., Jantaro, S., Khetkorn, W., Lindblad, P. & Incharoensakdi, A. Increased H_2_ production in the cyanobacterium Synechocystis sp. strain PCC 6803 by redirecting the electron supply via genetic engineering of the nitrate assimilation pathway. Metab. Eng. 13, 610–616 (2011).

2. Angermayr, S. A. et al. Exploring metabolic engineering design principles for the photosynthetic production of lactic acid by Synechocystis sp. PCC6803. Biotechnol. Biofuels 7, 99 (2014).

3. Gao, Z., Zhao, H., Li, Z., Tan, X. & Lu, X. Photosynthetic production of ethanol from carbon dioxide in genetically engineered cyanobacteria. Energy Environ. Sci. 5, 9857–9865 (2012).

4. Angermayr, S. A., Gorchs Rovira, A. & Hellingwerf, K. J. Metabolic engineering of cyanobacteria for the synthesis of commodity products. Trends Biotechnol. 33, 352–361 (2015).

5. Matson, M. M. & Atsumi, S. Photomixotrophic chemical production in cyanobacteria. Curr. Opin. Biotechnol. 50, 65–71 (2018).

6. Liu, N., Santala, S. & Stephanopoulos, G. Mixed carbon substrates: a necessary nuisance or a missed opportunity? Curr. Opin. Biotechnol. 62, 15–21 (2020).

7. Yang, C., Hua, Q. & Shimizu, K. Metabolic Flux Analysis in Synechocystis Using Isotope Distribution from 13C-Labeled Glucose. Metab. Eng. 4, 202–216 (2002).

8. Lee, T.-C. et al. Engineered xylose utilization enhances bio-products productivity in the cyanobacterium Synechocystis sp. PCC 6803. Metab. Eng. 30, 179–189 (2015).

9. Kanno, M., Carroll, A. L. & Atsumi, S. Global metabolic rewiring for improved CO_2_ fixation and chemical production in cyanobacteria. Nat. Commun. 8, 14724 (2017).

10. Cruz, J. A. Plasticity in light reactions of photosynthesis for energy production and photoprotection. J. Exp. Bot. 56, 395–406 (2004).

11. Kämäräinen, J. et al. Pyridine nucleotide transhydrogenase PntAB is essential for optimal growth and photosynthetic integrity under low-light mixotrophic conditions in Synechocystis sp. PCC 6803. New Phytol. 214, 194–204 (2017).

12. Kramer, D. M. & Evans, J. R. The Importance of Energy Balance in Improving Photosynthetic Productivity. Plant Physiol. 155, 70–78 (2011).

13. Joliot, P. & Joliot, A. Cyclic electron transfer in plant leaf. Proc. Natl. Acad. Sci. 99, 10209–10214 (2002).

14. Gao, H. & Xu, X. The Cyanobacterial NAD Kinase Gene sll1415 Is Required for Photoheterotrophic Growth and Cellular Redox Homeostasis in Synechocystis sp. Strain PCC 6803. J. Bacteriol. 194, 218–224 (2012).

15. Chen, X., Li, S. & Liu, L. Engineering redox balance through cofactor systems. Trends Biotechnol. 32, 337–343 (2014).

16. Yang, H., Jia, X. & Han, Y. Microbial redox coenzyme engineering and applications in biosynthesis. Trends Microbiol. 30, 318–321 (2022).

17. Tamoi, M., Miyazaki, T., Fukamizo, T. & Shigeoka, S. The Calvin cycle in cyanobacteria is regulated by CP12 via the NAD(H)/NADP(H) ratio under light/dark conditions. Plant J. 42, 504–513 (2005).

18. Takahashi, H., Uchimiya, H. & Hihara, Y. Difference in metabolite levels between photoautotrophic and photomixotrophic cultures of Synechocystis sp. PCC 6803 examined by capillary electrophoresis electrospray ionization mass spectrometry. J. Exp. Bot. 59, 3009–3018 (2008).

19. Meyer, D., Buhler, B. & Schmid, A. Process and Catalyst Design Objectives for Specific Redox Biocatalysis. in 53–91 (2006). doi:10.1016/S0065-2164(06)59003-3.

20. Orth, J. D., Thiele, I. & Palsson, B. Ø. What is flux balance analysis? Nat. Biotechnol. 28, 245–248 (2010).

21. Hendry, J. I., Bandyopadhyay, A., Srinivasan, S., Pakrasi, H. B. & Maranas, C. D. Metabolic model guided strain design of cyanobacteria. Curr. Opin. Biotechnol. 64, 17–23 (2020).

22. Wilson, J., Gering, S., Pinard, J., Lucas, R. & Briggs, B. R. Bio-production of gaseous alkenes: ethylene, isoprene, isobutene. Biotechnol. Biofuels 11, 234 (2018).

23. Friedlingstein, P. et al. Global Carbon Budget 2020. Earth Syst. Sci. Data 12, 3269–3340 (2020).

24. Ögmundarson, Ó., Herrgård, M. J., Forster, J., Hauschild, M. Z. & Fantke, P. Addressing environmental sustainability of biochemicals. Nat. Sustain. 3, 167–174 (2020).

25. Kang, M.-K. & Nielsen, J. Biobased production of alkanes and alkenes through metabolic engineering of microorganisms. J. Ind. Microbiol. Biotechnol. 44, 613–622 (2017).

26. He, G. et al. Rapid cost decrease of renewables and storage accelerates the decarbonization of China’s power system. Nat. Commun. 11, 2486 (2020).

27. Lewis, N. E. et al. Omic data from evolved E. coli are consistent with computed optimal growth from genome-scale models. Mol. Syst. Biol. 6, 390 (2010).

28. Ebrahim, A., Lerman, J. A., Palsson, B. O. & Hyduke, D. R. COBRApy: COnstraints-Based Reconstruction and Analysis for Python. BMC Syst. Biol. 7, 74 (2013).

29. Nogales, J., Gudmundsson, S., Knight, E. M., Palsson, B. O. & Thiele, I. Detailing the optimality of photosynthesis in cyanobacteria through systems biology analysis. Proc. Natl. Acad. Sci. 109, 2678–2683 (2012).

30. King, Z. A. et al. BiGG Models: A platform for integrating, standardizing and sharing genome-scale models. Nucleic Acids Res. 44, D515–D522 (2016).

31. Zhang, S. & Bryant, D. A. The Tricarboxylic Acid Cycle in Cyanobacteria. Science (80-.). 334, 1551–1553 (2011).

32. Steinhauser, D., Fernie, A. R. & Aryaújo, W. L. Unusual cyanobacterial TCA cycles: not broken just different. Trends Plant Sci. 17, 503–509 (2012).

33. Xiong, W., Brune, D. & Vermaas, W. F. J. The γ-aminobutyric acid shunt contributes to closing the tricarboxylic acid cycle in Synechocystis sp. PCC 6803. Mol. Microbiol. 93, 786–796 (2014).

34. Xiong, W. et al. Phosphoketolase pathway contributes to carbon metabolism in cyanobacteria. Nat. Plants 2, 15187 (2016).

35. Bachhar, A. & Jablonsky, J. A new insight into role of phosphoketolase pathway in Synechocystis sp. PCC 6803. Sci. Rep. 10, 22018 (2020).

36. Chen, X. et al. The Entner–Doudoroff pathway is an overlooked glycolytic route in cyanobacteria and plants. Proc. Natl. Acad. Sci. 113, 5441–5446 (2016).

37. Klemke, F. et al. Identification of the light-independent phosphoserine pathway as an additional source of serine in the cyanobacterium Synechocystis sp. PCC 6803. Microbiology 161, 1050–1060 (2015).

38. Bonner, C. A., Jensen, R. A., Gander, J. E. & Keyhani, N. O. A core catalytic domain of the TyrA protein family: arogenate dehydrogenase from Synechocystis. Biochem. J. 382, 279–291 (2004).

39. Lea-Smith, D. J., Bombelli, P., Vasudevan, R. & Howe, C. J. Photosynthetic, respiratory and extracellular electron transport pathways in cyanobacteria. Biochim. Biophys. Acta - Bioenerg. 1857, 247–255 (2016).

40. Cooley, J. W. & Vermaas, W. F. J. Succinate Dehydrogenase and Other Respiratory Pathways in Thylakoid Membranes of Synechocystis sp. Strain PCC 6803: Capacity Comparisons and Physiological Function. J. Bacteriol. 183, 4251–4258 (2001).

41. Varma, A., Boesch, B. W. & Palsson, B. O. Stoichiometric interpretation of Escherichia coli glucose catabolism under various oxygenation rates. Appl. Environ. Microbiol. 59, 2465–2473 (1993).

42. Yunus, I. S. et al. Synthetic metabolic pathways for photobiological conversion of CO_2_ into hydrocarbon fuel. Metab. Eng. 49, 201–211 (2018).

43. Mustila, H., Kugler, A. & Stensjö, K. Isobutene production in Synechocystis sp. PCC 6803 by introducing α-ketoisocaproate dioxygenase from Rattus norvegicus. Metab. Eng. Commun. 12, e00163 (2021).

44. Ungerer, J. et al. Sustained photosynthetic conversion of CO_2_ to ethylene in recombinant cyanobacterium Synechocystis 6803. Energy Environ. Sci. 5, 8998 (2012).

45. Lindberg, P., Park, S. & Melis, A. Engineering a platform for photosynthetic isoprene production in cyanobacteria, using Synechocystis as the model organism. Metab. Eng. 12, 70–79 (2010).

46. Baldwin, J. E. et al. 4-Hydroxyphenylpyruvate dioxygenase appears to display α-ketoisocaproate dioxygenase activity in rat liver. Bioorg. Med. Chem. Lett. 5, 1255–1260 (1995).

47. Rossoni, L., Hall, S. J., Eastham, G., Licence, P. & Stephens, G. The Putative Mevalonate Diphosphate Decarboxylase from Picrophilus torridus Is in Reality a Mevalonate-3-Kinase with High Potential for Bioproduction of Isobutene. Appl. Environ. Microbiol. 81, 2625–2634 (2015).

48. Feist, A. M. & Palsson, B. O. The biomass objective function. Curr. Opin. Microbiol. 13, 344–349 (2010).

49. King, Z. A. et al. Escher: A Web Application for Building, Sharing, and Embedding Data-Rich Visualizations of Biological Pathways. PLOS Comput. Biol. 11, e1004321 (2015).

50. Mahadevan, R. & Schilling, C. H. The effects of alternate optimal solutions in constraint-based genome-scale metabolic models. Metab. Eng. 5, 264–276 (2003).

51. Chung, B. K. S. & Lee, D.-Y. Flux-sum analysis: a metabolite-centric approach for understanding the metabolic network. BMC Syst. Biol. 3, 117 (2009).

52. Santos-Merino, M. et al. Improved photosynthetic capacity and photosystem I oxidation via heterologous metabolism engineering in cyanobacteria. Proc. Natl. Acad. Sci. 118, (2021).

53. Yunus, I. S. et al. Synthetic metabolic pathways for conversion of CO_2_ into secreted short-to medium-chain hydrocarbons using cyanobacteria. Metab. Eng. 72, 14–23 (2022).

54. Kämäräinen, J. et al. Physiological tolerance and stoichiometric potential of cyanobacteria for hydrocarbon fuel production. J. Biotechnol. 162, 67–74 (2012).

55. Knoop, H. & Steuer, R. A Computational Analysis of Stoichiometric Constraints and Trade-Offs in Cyanobacterial Biofuel Production. Front. Bioeng. Biotechnol. 3, (2015).

56. Forti, G., Furia, A., Bombelli, P. & Finazzi, G. In Vivo Changes of the Oxidation-Reduction State of NADP and of the ATP/ADP Cellular Ratio Linked to the Photosynthetic Activity in Chlamydomonas reinhardtii. Plant Physiol. 132, 1464–1474 (2003).

57. Reznik, E., Mehta, P. & Segrè, D. Flux Imbalance Analysis and the Sensitivity of Cellular Growth to Changes in Metabolite Pools. PLoS Comput. Biol. 9, e1003195 (2013).

58. Brinkmann-Chen, S., Cahn, J. K. B. & Arnold, F. H. Uncovering rare NADH-preferring ketol-acid reductoisomerases. Metab. Eng. 26, 17–22 (2014).

59. Brinkmann-Chen, S. et al. General approach to reversing ketol-acid reductoisomerase cofactor dependence from NADPH to NADH. Proc. Natl. Acad. Sci. 110, 10946–10951 (2013).

60. Jackson, J. B. Proton translocation by transhydrogenase. FEBS Lett. 545, 18–24 (2003).

61. Gopalakrishnan, S. & Maranas, C. D. 13C metabolic flux analysis at a genome-scale. Metab. Eng. 32, 12–22 (2015).

62. Young, J. D., Shastri, A. A., Stephanopoulos, G. & Morgan, J. A. Mapping photoautotrophic metabolism with isotopically nonstationary (13)C flux analysis. Metab. Eng. 13, 656–665 (2011).

63. You, L., Berla, B., He, L., Pakrasi, H. B. & Tang, Y. J. 13C-MFA delineates the photomixotrophic metabolism of Synechocystis sp. PCC 6803 under light-and carbon-sufficient conditions. Biotechnol. J. 9, 684–692 (2014).

64. Jazmin, L. J. et al. Isotopically nonstationary 13C flux analysis of cyanobacterial isobutyraldehyde production. Metab. Eng. 42, 9–18 (2017).

65. Nishiguchi, H. et al. Transomics data-driven, ensemble kinetic modeling for system-level understanding and engineering of the cyanobacteria central metabolism. Metab. Eng. 52, 273–283 (2019).

66. Hu, G. et al. Engineering synergetic CO_2_-fixing pathways for malate production. Metab. Eng. 47, 496–504 (2018).

67. Chuang, D. S.-W. & Liao, J. C. Role of cyanobacterial phosphoketolase in energy regulation and glucose secretion under dark anaerobic and osmotic stress conditions. Metab. Eng. 65, 255–262 (2021).

68. Kramer, D. M., Avenson, T. J. & Edwards, G. E. Dynamic flexibility in the light reactions of photosynthesis governed by both electron and proton transfer reactions. Trends Plant Sci. 9, 349–357 (2004).

69. Almaas, E., Kovács, B., Vicsek, T., Oltvai, Z. N. & Barabási, A.-L. Global organization of metabolic fluxes in the bacterium Escherichia coli. Nature 427, 839–843 (2004).

70. Noor, E., Eden, E., Milo, R. & Alon, U. Central Carbon Metabolism as a Minimal Biochemical Walk between Precursors for Biomass and Energy. Mol. Cell 39, 809–820 (2010).

71. Wan, N. et al. Cyanobacterial carbon metabolism: Fluxome plasticity and oxygen dependence. Biotechnol. Bioeng. 114, 1593–1602 (2017).

72. Xiong, W. et al. The plasticity of cyanobacterial metabolism supports direct CO_2_ conversion to ethylene. Nat. Plants 1, 15053 (2015).

73. Lynch, S., Eckert, C., Yu, J., Gill, R. & Maness, P.-C. Overcoming substrate limitations for improved production of ethylene in E. coli. Biotechnol. Biofuels 9, 3 (2016).

74. Durall, C., Lindberg, P., Yu, J. & Lindblad, P. Increased ethylene production by overexpressing phosphoenolpyruvate carboxylase in the cyanobacterium Synechocystis PCC 6803. Biotechnol. Biofuels 13, 16 (2020).

75. Kaplan, A. & Reinhold, L. CO_2_ concentrating mechanisms in photosynthetic microorganisms. Annu. Rev. Plant Physiol. Plant Mol. Biol. 50, 539–570 (1999).

76. Zhang, Y., Adams, I. P. & Ratledge, C. Malic enzyme: the controlling activity for lipid production? Overexpression of malic enzyme in Mucor circinelloides leads to a 2.5-fold increase in lipid accumulation. Microbiology 153, 2013–2025 (2007).

77. Wynn, J. P., Hamid, A. bin A. & Ratledge, C. The role of malic enzyme in the regulation of lipid accumulation in filamentous fungi. Microbiology 145, 1911–1917 (1999).

78. Wise, E. M. & Ball, E. G. Malic enzyme and lipogenesis. Proc. Natl. Acad. Sci. 52, 1255–1263 (1964).

79. Cano, M. et al. Glycogen Synthesis and Metabolite Overflow Contribute to Energy Balancing in Cyanobacteria. Cell Rep. 23, 667–672 (2018).

80. Holland, S. C. et al. Impacts of genetically engineered alterations in carbon sink pathways on photosynthetic performance. Algal Res. 20, 87–99 (2016).

81. Makowka, A. et al. Glycolytic Shunts Replenish the Calvin–Benson–Bassham Cycle as Anaplerotic Reactions in Cyanobacteria. Mol. Plant 13, 471–482 (2020).

82. Wang, Y. et al. Biosynthesis of platform chemical 3-hydroxypropionic acid (3-HP) directly from CO2 in cyanobacterium Synechocystis sp. PCC 6803. Metab. Eng. 34, 60–70 (2016).

83. Chin, T., Okuda, Y. & Ikeuchi, M. Sorbitol production and optimization of photosynthetic supply in the cyanobacterium Synechocystis PCC 6803. J. Biotechnol. 276–277, 25–33 (2018).

84. Lu, H. et al. Time-resolved multi-omics analysis reveals the role of nutrient stress-induced resource reallocation for TAG accumulation in oleaginous fungus Mortierella alpina. Biotechnol. Biofuels 13, 116 (2020).

85. Pan, Y. et al. Amino Acid Catabolism During Nitrogen Limitation in Phaeodactylum tricornutum. Front. Plant Sci. 11, (2020).

86. Giberti, S., Funck, D. & Forlani, G. Δ 1 -pyrroline-5-carboxylate reductase from Arabidopsis thaliana: stimulation or inhibition by chloride ions and feedback regulation by proline depend on whether NADPH or NADH acts as co-substrate. New Phytol. 202, 911–919 (2014).

87. Li, Z. et al. Cancer cells depend on environmental lipids for proliferation when electron acceptors are limited. Nat. Metab. 4, 711–723 (2022).

88. Ryu, J.-Y. et al. NADPH dehydrogenase-mediated respiratory electron transport in thylakoid membranes of the cyanobacterium Synechocystis sp. PCC 6803 is inactive in the light. Mol. Cells 15, 240–4 (2003).

89. Johansson, N., Quehl, P., Norbeck, J. & Larsson, C. Identification of factors for improved ethylene production via the ethylene forming enzyme in chemostat cultures of Saccharomyces cerevisiae. Microb. Cell Fact. 12, 89 (2013).

90. Purdy, H. M., Pfleger, B. F. & Reed, J. L. Introduction of NADH-dependent nitrate assimilation in Synechococcus sp. PCC 7002 improves photosynthetic production of 2-methyl-1-butanol and isobutanol. Metab. Eng. 69, 87–97 (2022).

91. Pernil, R., Herrero, A. & Flores, E. Catabolic Function of Compartmentalized Alanine Dehydrogenase in the Heterocyst-Forming Cyanobacterium Anabaena sp. Strain PCC 7120. J. Bacteriol. 192, 5165–5172 (2010).

92. Ishikawa, Y. et al. One of the NAD kinases, sll1415, is required for the glucose metabolism of Synechocystis sp. PCC 6803. Plant J. 98, 654–666 (2019).

93. Ishikawa, Y. et al. The NAD Kinase Slr0400 Functions as a Growth Repressor in Synechocystis sp. PCC 6803. Plant Cell Physiol. 62, 668–677 (2021).

94. Fan, J. et al. Quantitative flux analysis reveals folate-dependent NADPH production. Nature 510, 298–302 (2014).

95. Locasale, J. W. et al. Phosphoglycerate dehydrogenase diverts glycolytic flux and contributes to oncogenesis. Nat. Genet. 43, 869–874 (2011).

96. Possemato, R. et al. Functional genomics reveal that the serine synthesis pathway is essential in breast cancer. Nature 476, 346–350 (2011).

97. Mills, L. A., McCormick, A. J. & Lea-Smith, D. J. Current knowledge and recent advances in understanding metabolism of the model cyanobacterium Synechocystis sp. PCC 6803. Biosci. Rep. 40, (2020).

98. Klaus, S. M. J. et al. Higher Plant Plastids and Cyanobacteria Have Folate Carriers Related to Those of Trypanosomatids. J. Biol. Chem. 280, 38457–38463 (2005).

