## Supplementary material for "Optimal energy and redox metabolism in the cyanobacterium *Synechocystis* sp. PCC 6803"

**Supplementary Figure. 1**

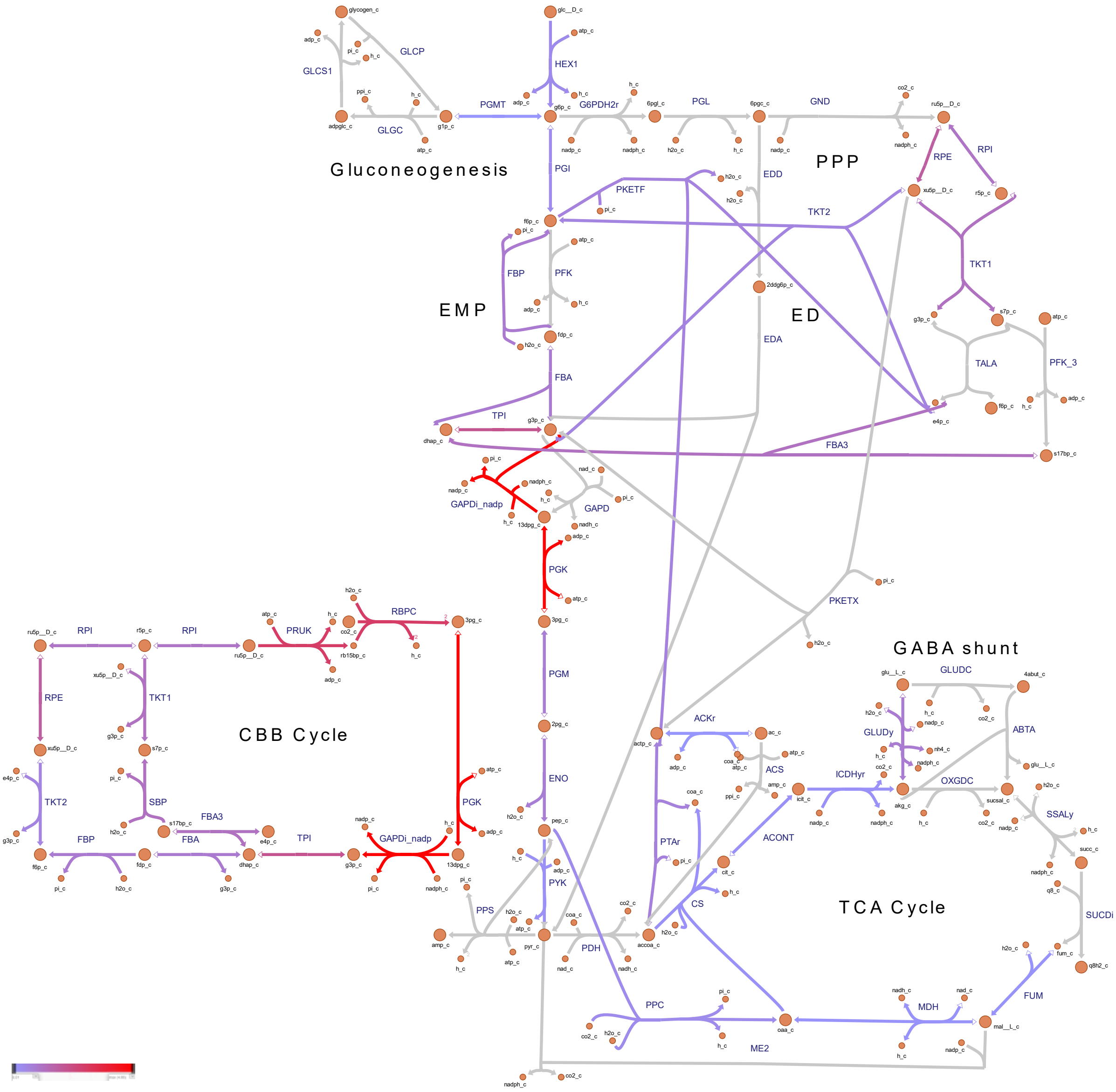


Supplementary Fig. 1 Metabolic flux map through central carbon metabolism for *Synechocystis* sp. PCC 6803 overproducing biomass, simulated to grow under mixotrophic conditions. Reaction fluxes (mmol/gDW/h) were predicted using pFBA. For interpretation of the references to color in this Fig legend, the reader is referred to the web version of this article.

**Supplementary Figure. 2**

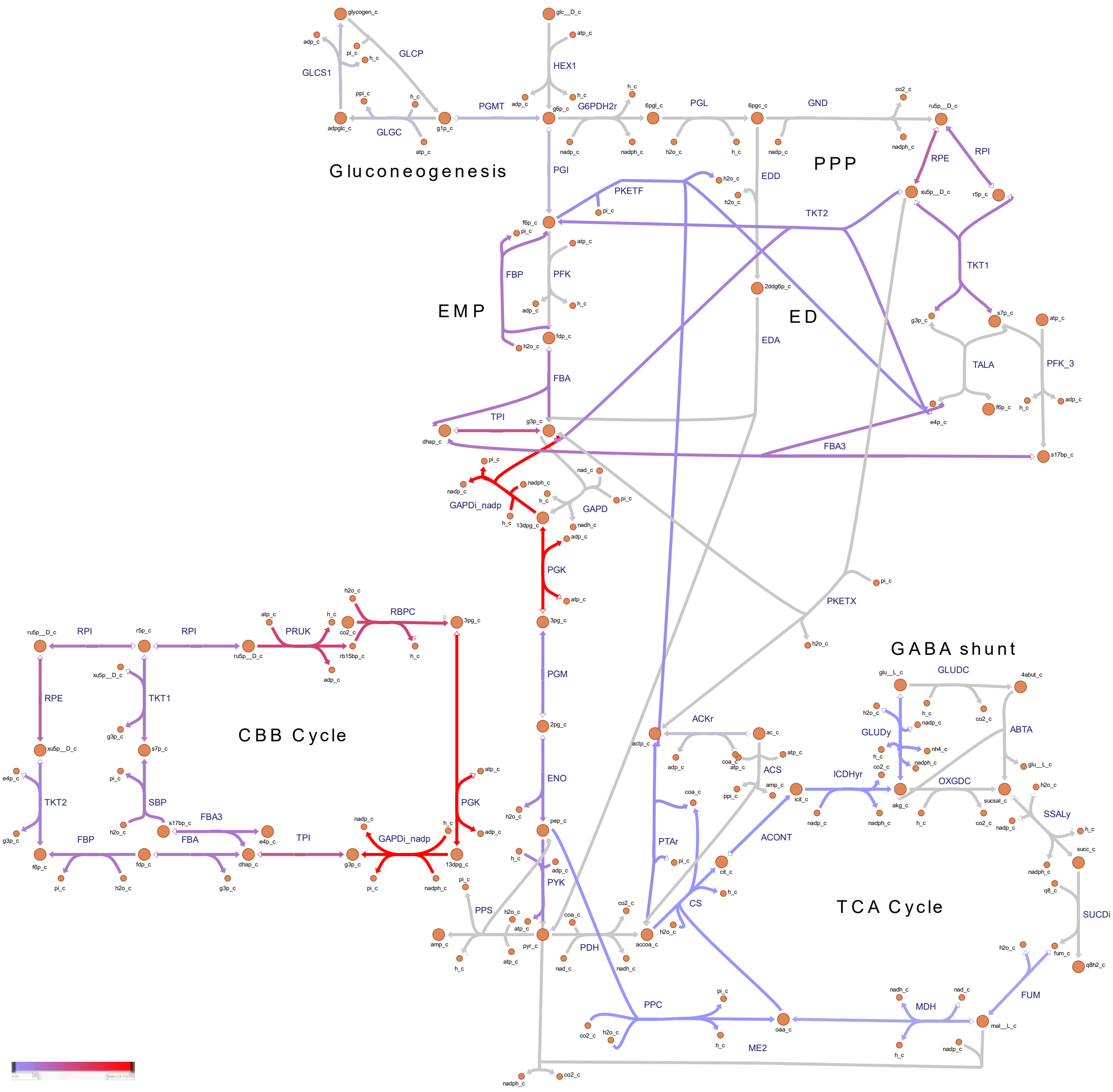


Supplementary Fig. 2 Metabolic flux map through central carbon metabolism for *Synechocystis* sp. PCC 6803 overproducing isobutene, simulated to grow under photoautotrophic conditions. Reaction fluxes (mmol/gDW/h) were predicted using pFBA. For interpretation of the references to color in this Fig legend, the reader is referred to the web version of this article.

**Supplementary Figure. 3**

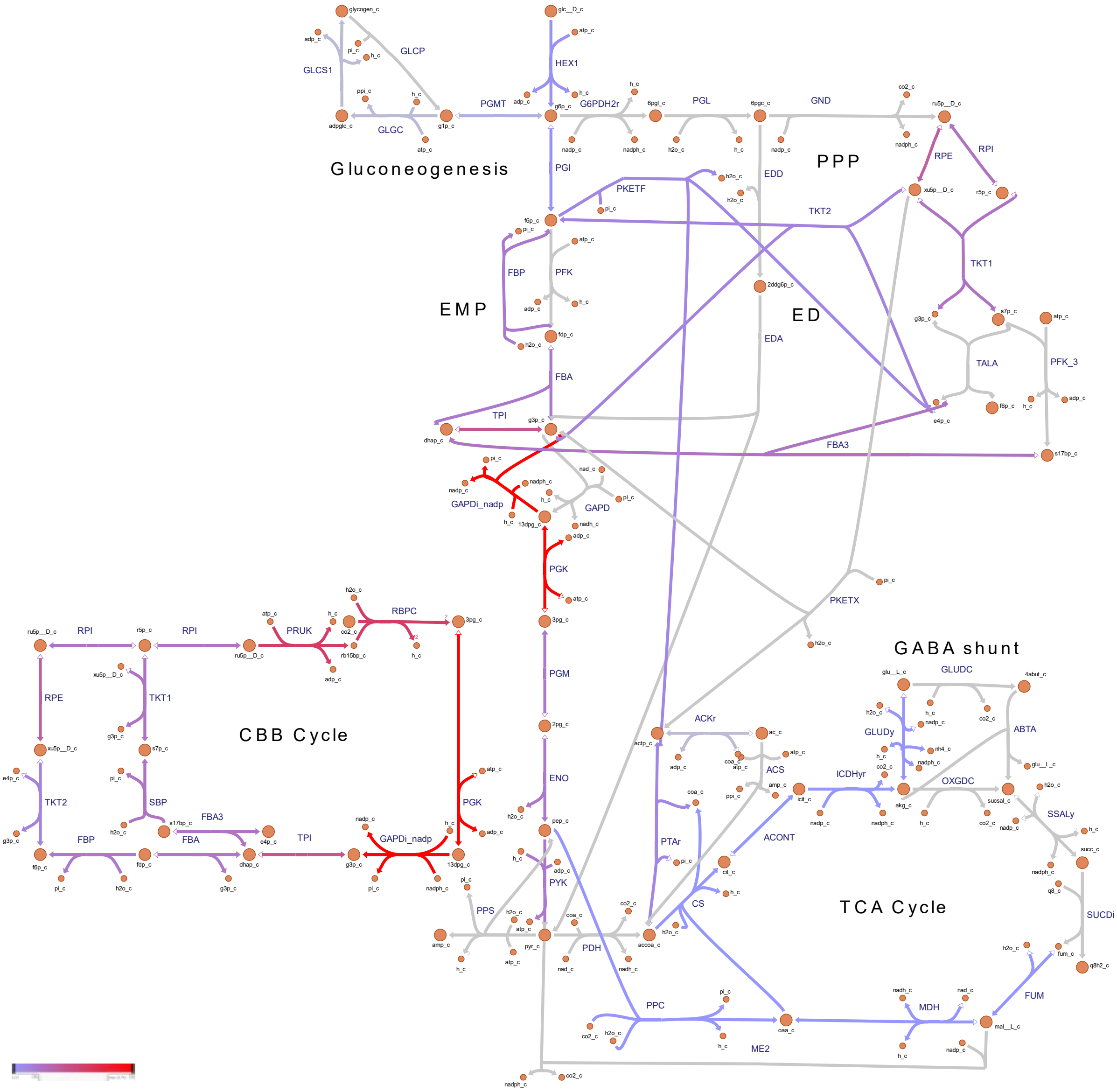


Supplementary Fig. 3 Metabolic flux map through central carbon metabolism for *Synechocystis* sp. PCC 6803 overproducing isobutene, simulated to grow under mixotrophic conditions. Reaction fluxes (mmol/gDW/h) were predicted using pFBA. For interpretation of the references to color in this Fig legend, the reader is referred to the web version of this article.

**Supplementary Figure. 4**

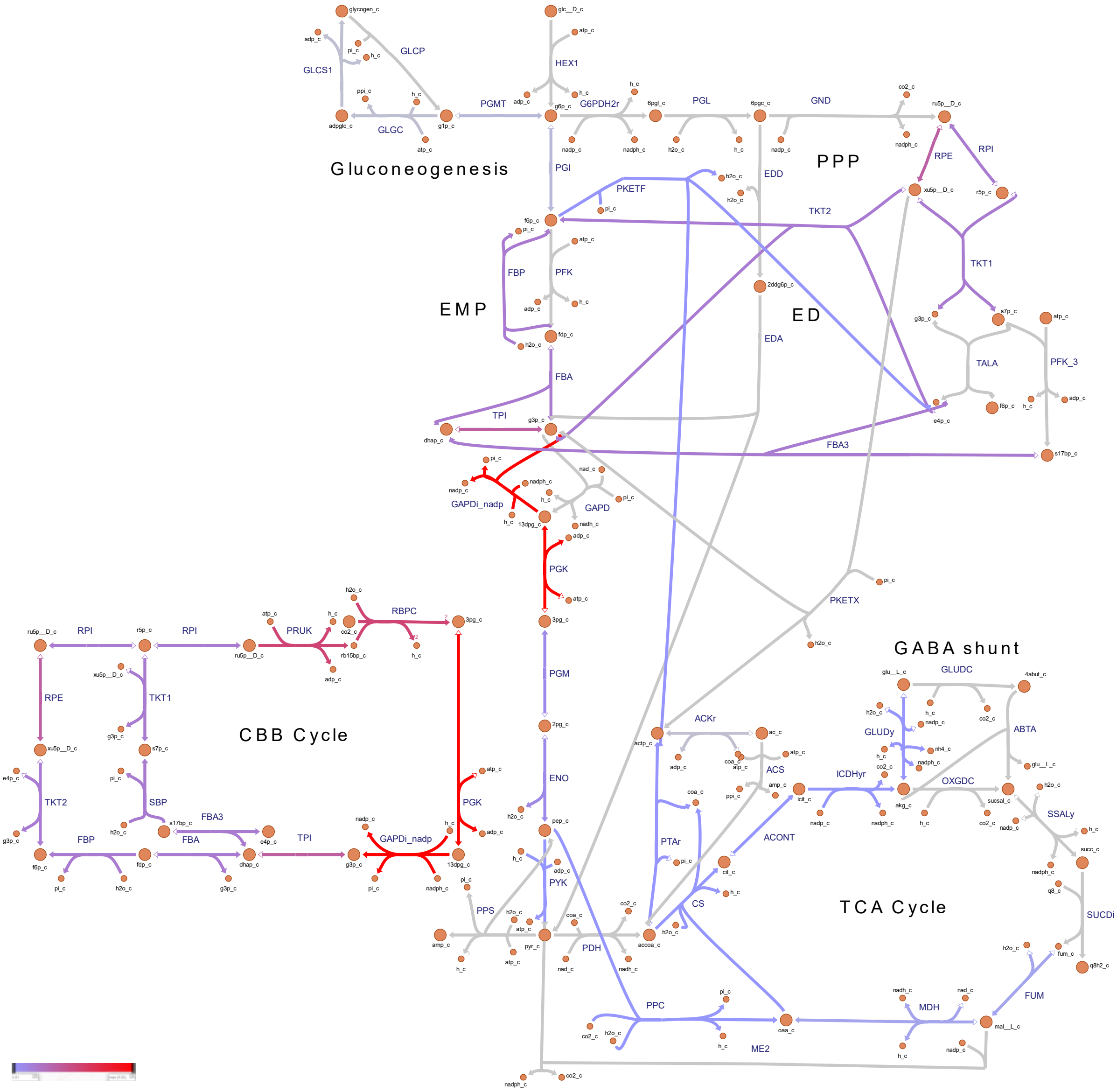


Supplementary Fig. 4 Metabolic flux map through central carbon metabolism for *Synechocystis* sp. PCC 6803 overproducing isoprene, simulated to grow under photoautotrophic conditions. Reaction fluxes (mmol/gDW/h) were predicted using pFBA. For interpretation of the references to color in this Fig legend, the reader is referred to the web version of this article.

**Supplementary Figure. 5**

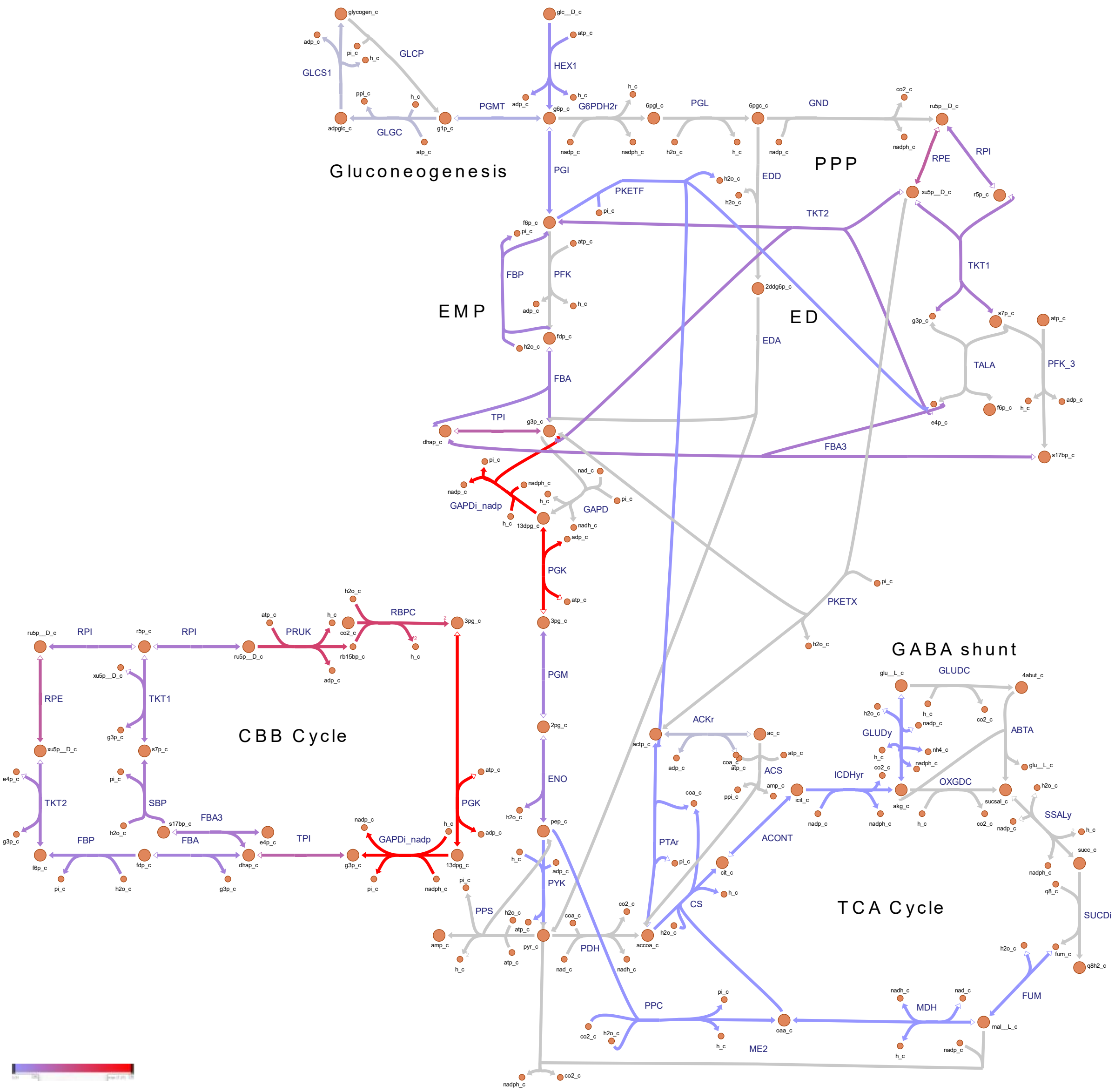


Supplementary Fig. 5 Metabolic flux map through central carbon metabolism for *Synechocystis* sp. PCC 6803 overproducing isoprene, simulated to grow under mixotrophic conditions. Reaction fluxes (mmol/gDW/h) were predicted using pFBA. For interpretation of the references to color in this Fig legend, the reader is referred to the web version of this article.

**Supplementary Figure. 6**

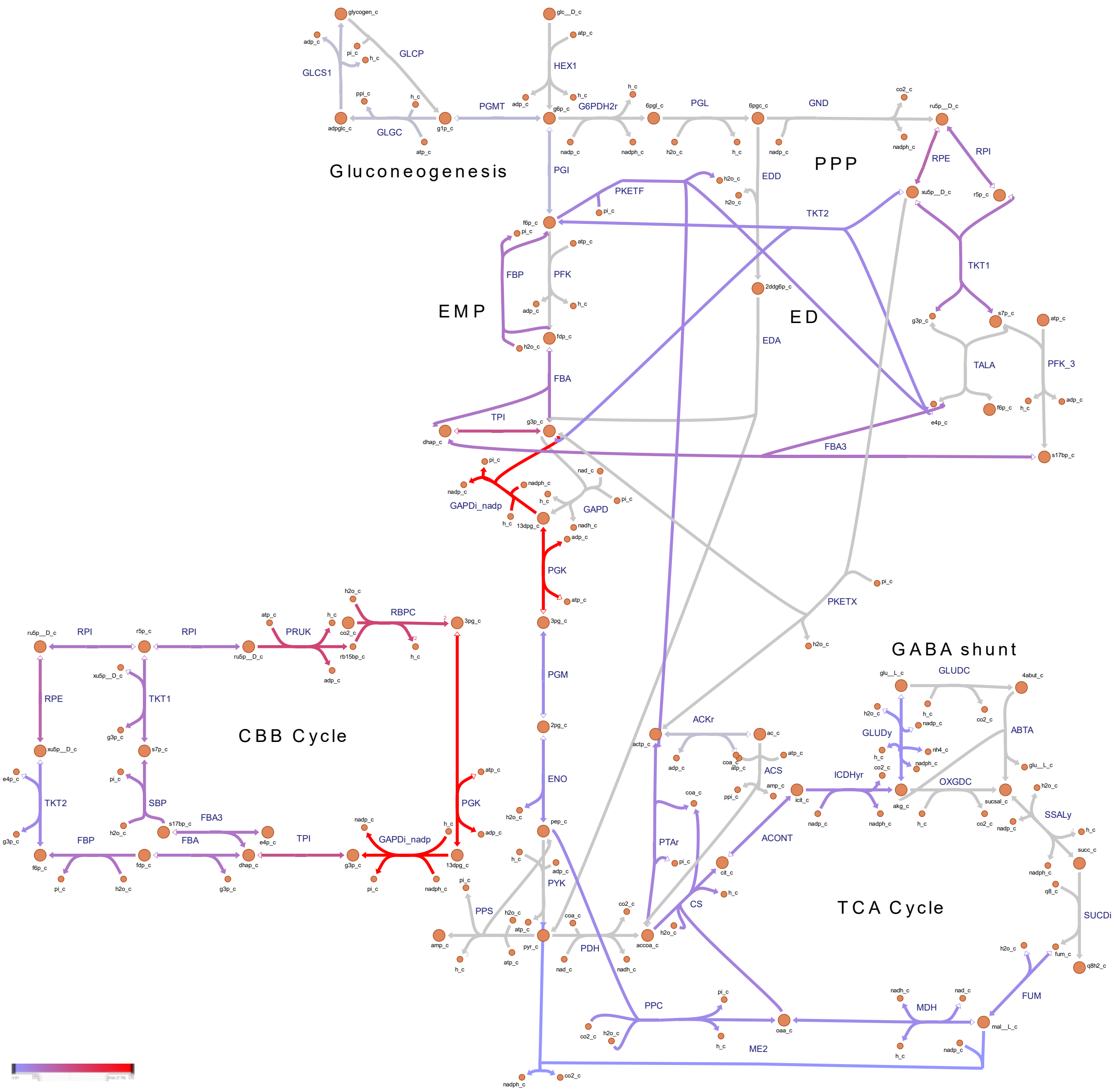


Supplementary Fig. 6 Metabolic flux map through central carbon metabolism for *Synechocystis* sp. PCC 6803 overproducing ethylene, simulated to grow under photoautotrophic conditions. Reaction fluxes (mmol/gDW/h) were predicted using pFBA. For interpretation of the references to color in this Fig legend, the reader is referred to the web version of this article.

**Supplementary Figure. 7**

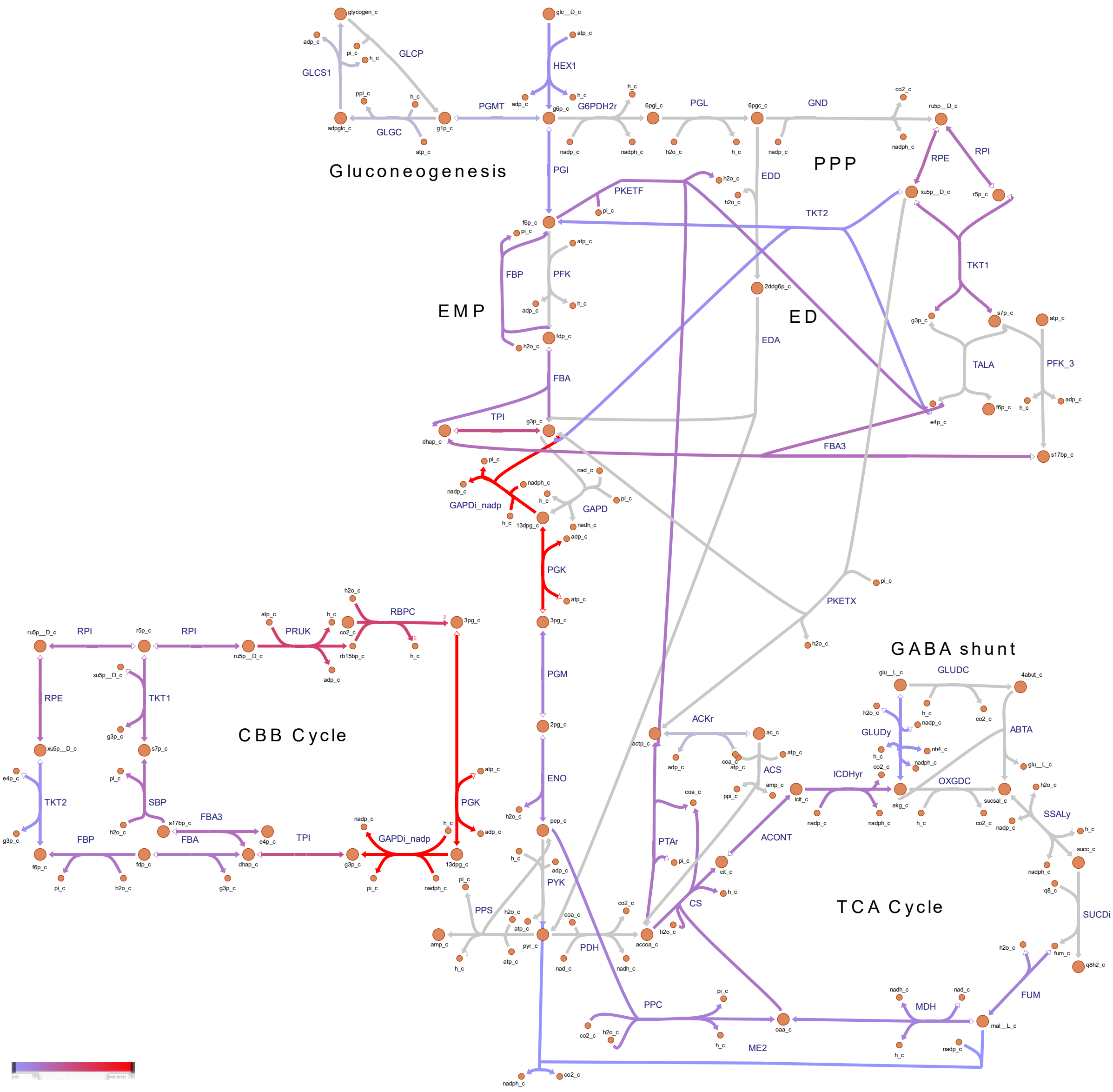


Supplementary Fig. 7 Metabolic flux map through central carbon metabolism for *Synechocystis* sp. PCC 6803 overproducing ethylene, simulated to grow under mixotrophic conditions. Reaction fluxes (mmol/gDW/h) were predicted using pFBA. For interpretation of the references to color in this Fig legend, the reader is referred to the web version of this article.

**Supplementary Figure. 8**

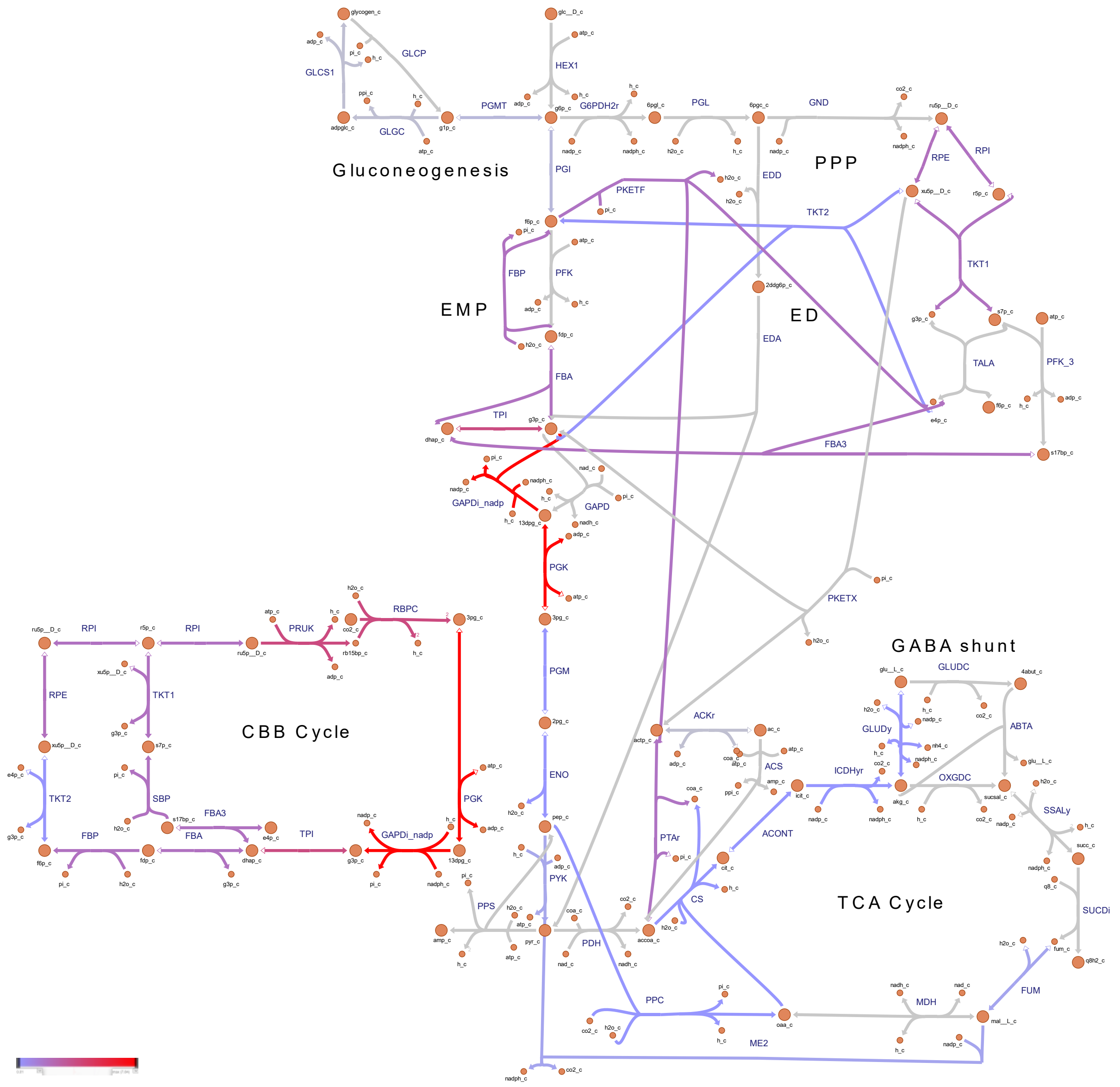


Supplementary Fig. 8 Metabolic flux map through central carbon metabolism for *Synechocystis* sp. PCC 6803 overproducing 1-undecene, simulated to grow under photoautotrophic conditions. Reaction fluxes (mmol/gDW/h) were predicted using pFBA. For interpretation of the references to color in this Fig legend, the reader is referred to the web version of this article.

**Supplementary Figure. 9**

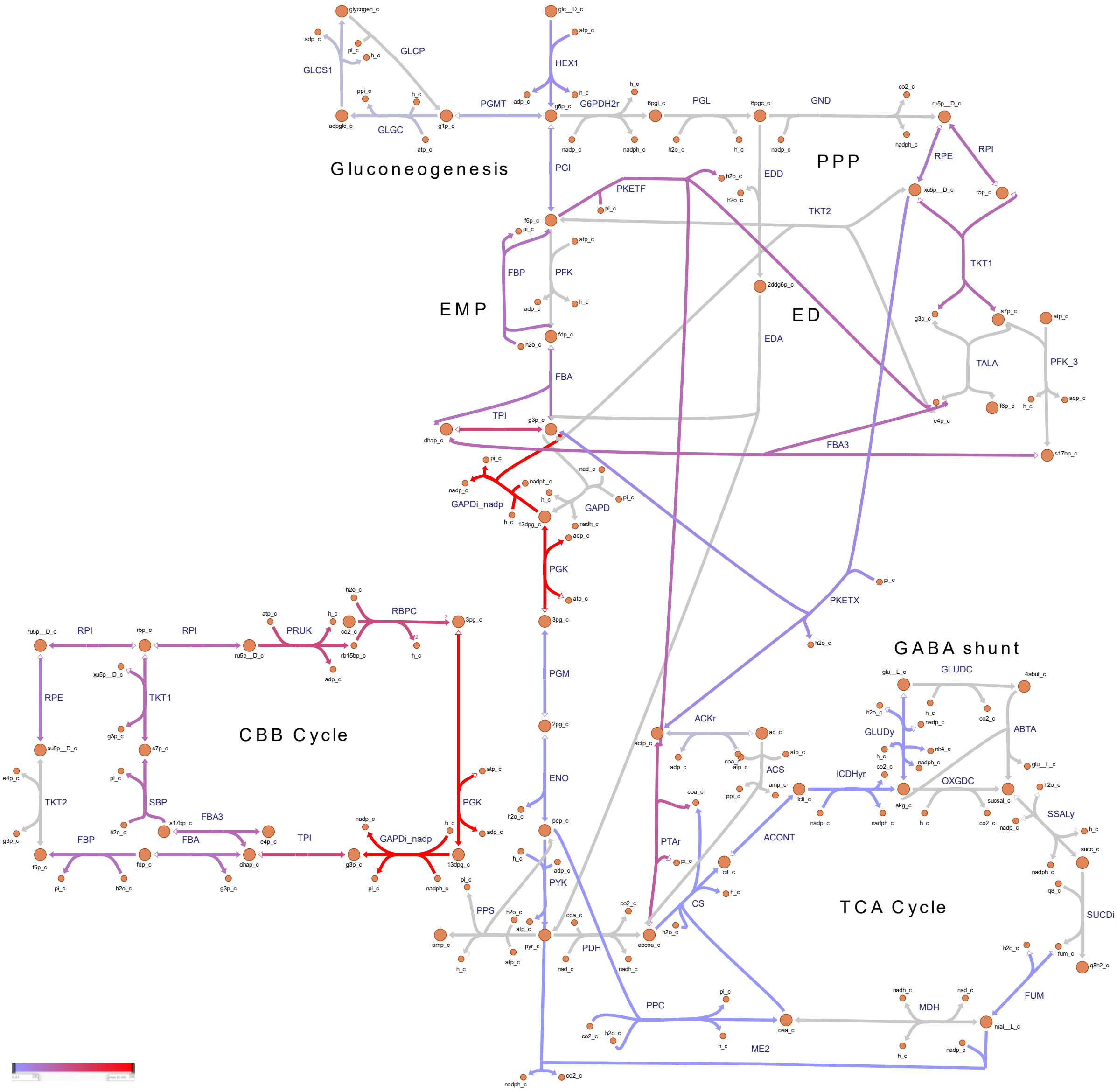


Supplementary Fig. 9 Metabolic flux map through central carbon metabolism for *Synechocystis* sp. PCC 6803 overproducing 1-undecene, simulated to grow under mixotrophic conditions. Reaction fluxes (mmol/gDW/h) were predicted using pFBA. For interpretation of the references to color in this Fig legend, the reader is referred to the web version of this article.

**Supplementary Figure. 11**

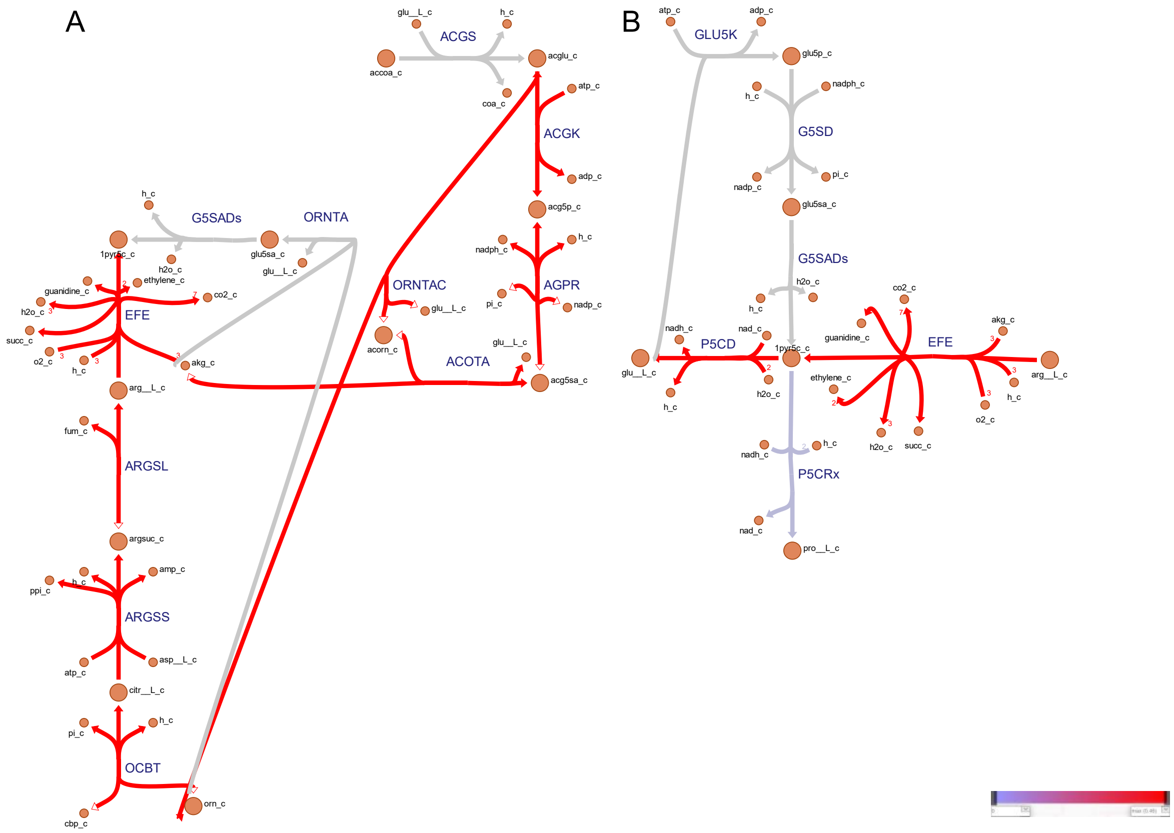


Supplementary Fig. 11 Metabolic flux map through reactions involved in balancing nitrogen metabolism for *Synechocystis* sp. PCC 6803 overproducing ethylene, simulated to grow under mixotrophic conditions. (A) L-glutamate and L-arginine regeneration through the urea cycle. B) L-glutamate and L-proline regeneration. Reaction fluxes (mmol/gDW/h) were predicted using pFBA. For interpretation of the references to color in this Fig legend, the reader is referred to the web version of this article.

**Supplementary Figure. 12**

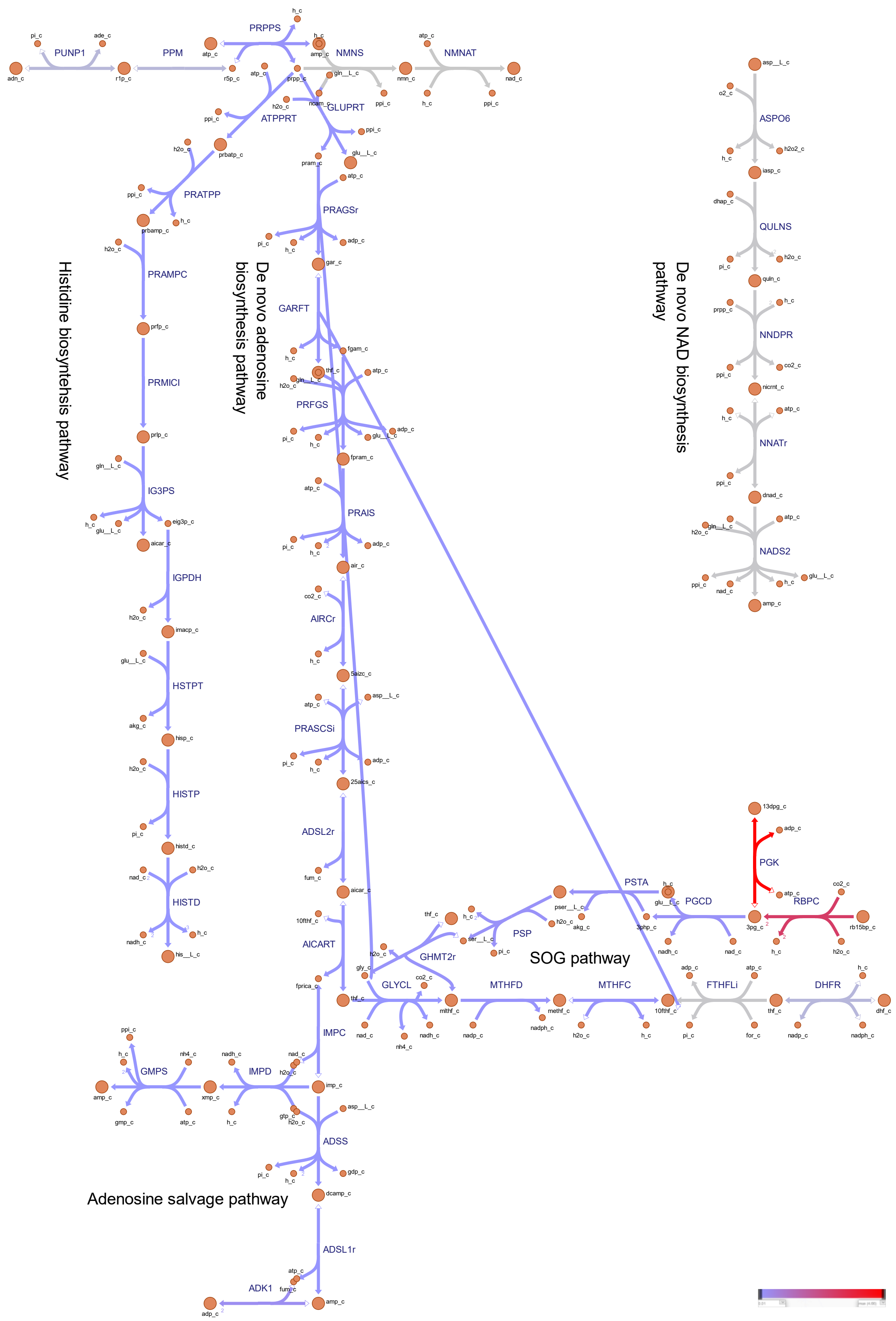


Supplementary Fig. 12 Metabolic flux map of auxiliary pathways enabling cellular energy and redox homeostasis. The auxiliary pathways are composed of the serine, one-carbon cycle, glycine synthesis (SOG) pathway and the biosynthesis of nucleotide precursors. *Synechocystis* sp. PCC 6803 was simulated to overproduce biomass, while grown under mixotrophic conditions. Reaction fluxes (mmol/gDW/h) were predicted using pFBA. For interpretation of the references to color in this Fig legend, the reader is referred to the web version of this article.

### Supplementary Figure. 13

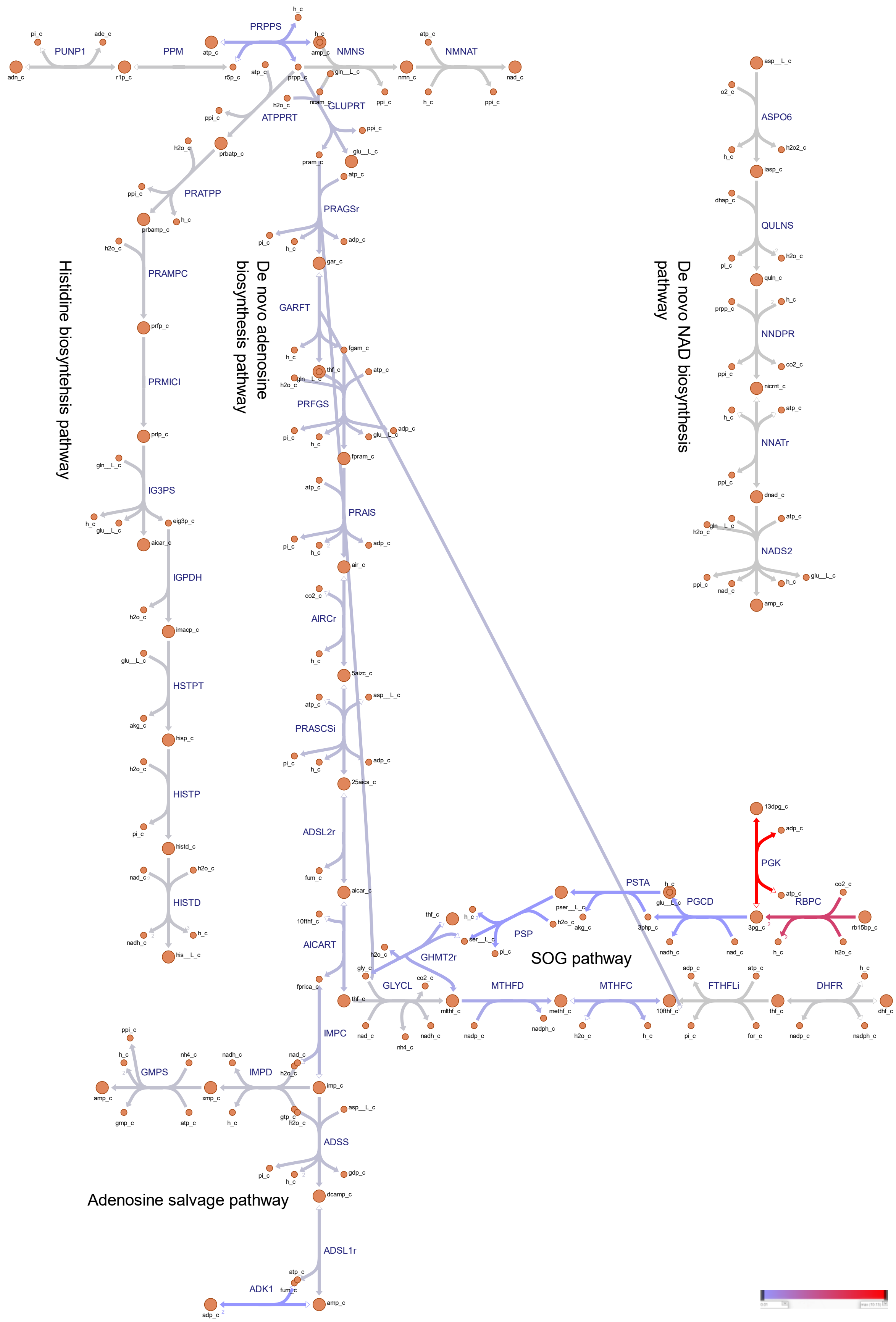


Supplementary Fig. 13 Metabolic flux map of auxiliary pathways enabling cellular energy and redox

homeostasis. The auxiliary pathways are composed of the serine, one-carbon cycle, glycine synthesis (SOG) pathway and the biosynthesis of nucleotide precursors. *Synechocystis* sp. PCC 6803 was simulated to overproduce isobutene, while grown under photoautotrophic conditions. Reaction fluxes (mmol/gDW/h) were predicted using pFBA. For interpretation of the references to color in this Fig legend, the reader is referred to the web version of this article.

### Supplementary Figure. 14

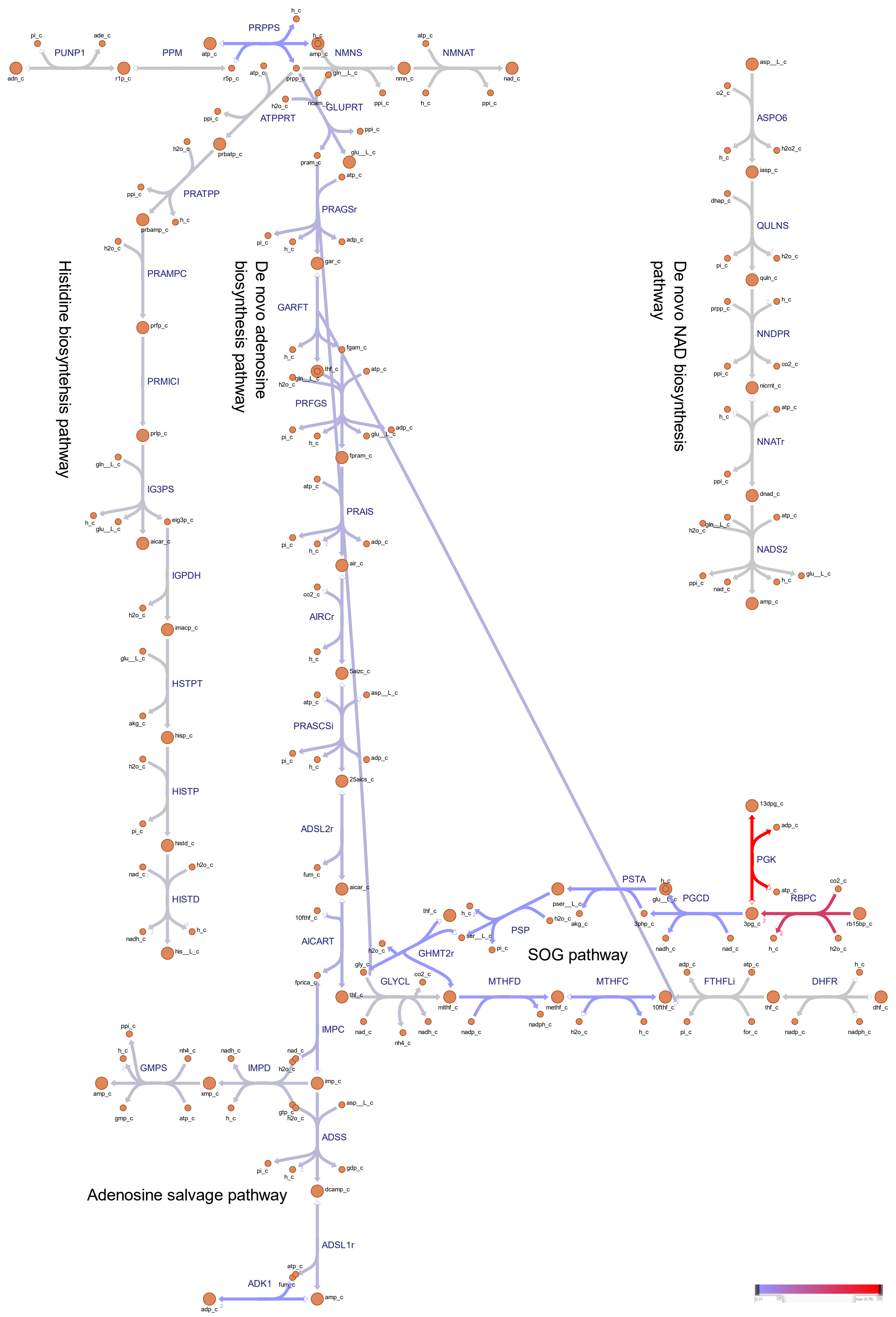


Supplementary Fig. 14 Metabolic flux map of auxiliary pathways enabling cellular energy and redox

homeostasis. The auxiliary pathways are composed of the serine, one-carbon cycle, glycine synthesis (SOG) pathway and the biosynthesis of nucleotide precursors. *Synechocystis* sp. PCC 6803 was simulated to overproduce isobutene, while grown under mixotrophic conditions. Reaction fluxes (mmol/gDW/h) were predicted using pFBA. For interpretation of the references to color in this Fig legend, the reader is referred to the web version of this article.

### Supplementary Figure. 15

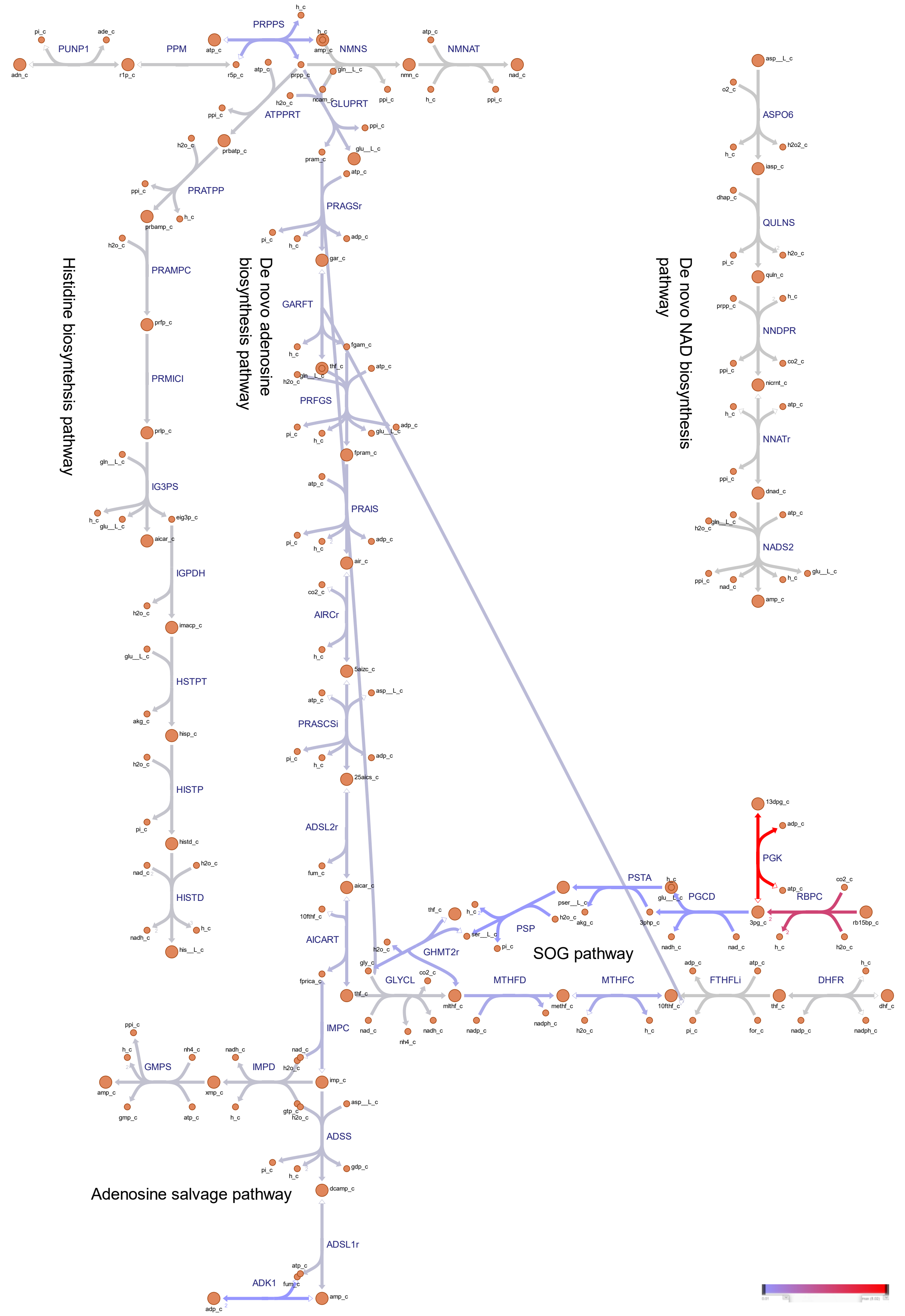


Supplementary Fig. 15 Metabolic flux map of auxiliary pathways enabling cellular energy and redox

homeostasis. The auxiliary pathways are composed of the serine, one-carbon cycle, glycine synthesis (SOG) pathway and the biosynthesis of nucleotide precursors. *Synechocystis* sp. PCC 6803 was simulated to overproduce isoprene, while grown under photoautotrophic conditions. Reaction fluxes (mmol/gDW/h) were predicted using pFBA. For interpretation of the references to color in this Fig legend, the reader is referred to the web version of this article.

**Supplementary Figure. 16**

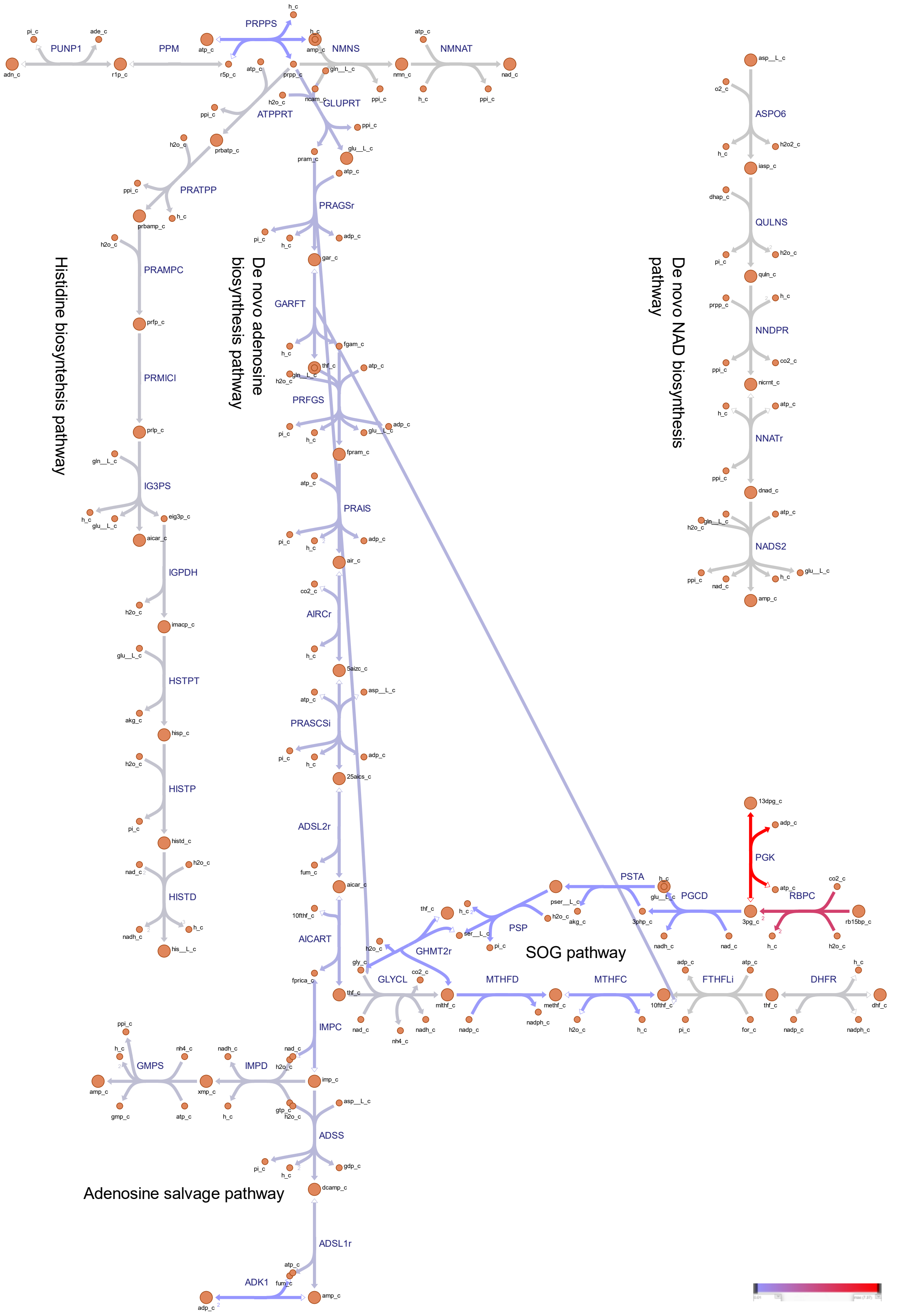


Supplementary Fig. 16 Metabolic flux map of auxiliary pathways enabling cellular energy and redox homeostasis. The auxiliary pathways are composed of the serine, one-carbon cycle, glycine synthesis (SOG) pathway and the biosynthesis of nucleotide precursors. *Synechocystis* sp. PCC 6803 was simulated to overproduce isoprene, while grown under mixotrophic conditions. Reaction fluxes (mmol/gDW/h) were predicted using pFBA. For interpretation of the references to color in this Fig legend, the reader is referred to the web version of this article.

**Supplementary Figure. 17**

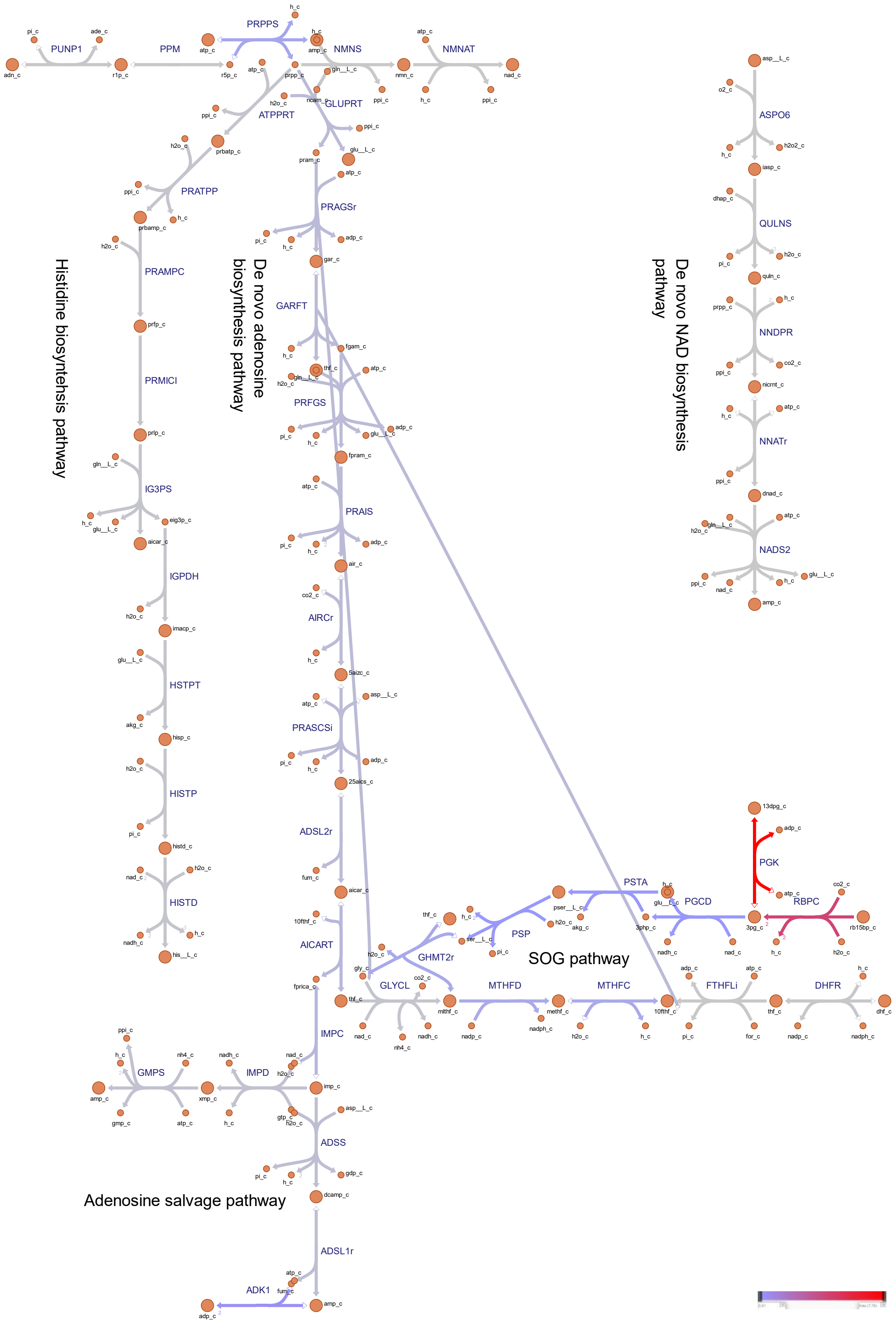


Supplementary Fig. 17 Metabolic flux map of auxiliary pathways enabling cellular energy and redox homeostasis. The auxiliary pathways are composed of the serine, one-carbon cycle, glycine synthesis (SOG) pathway and the biosynthesis of nucleotide precursors. *Synechocystis* sp. PCC 6803 was simulated to overproduce ethylene, while grown under photoautotrophic conditions. Reaction fluxes (mmol/gDW/h) were predicted using pFBA. For interpretation of the references to color in this Fig legend, the reader is referred to the web version of this article.

### Supplementary Figure. 18

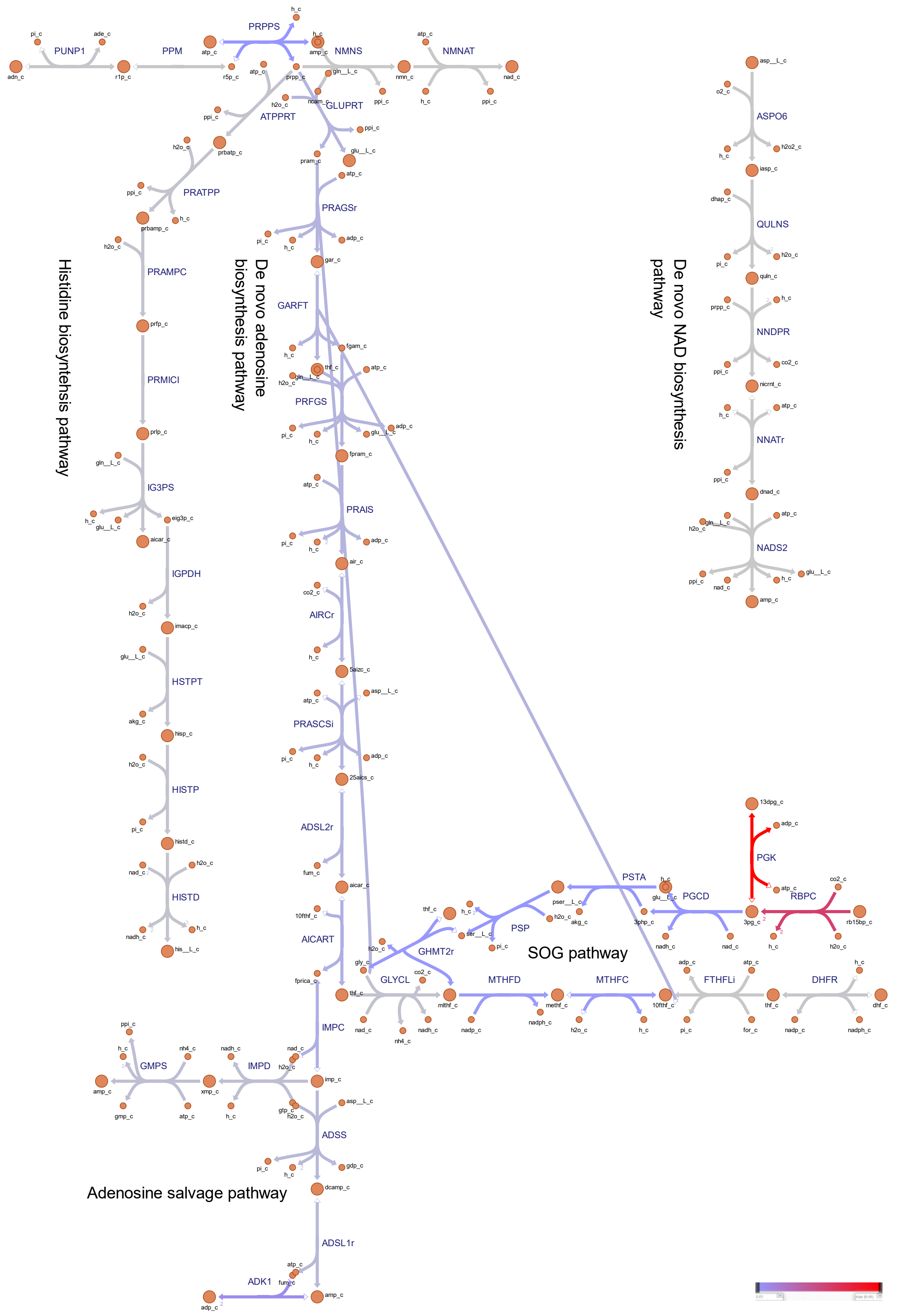


Supplementary Fig. 18 Metabolic flux map of auxiliary pathways enabling cellular energy and redox

homeostasis. The auxiliary pathways are composed of the serine, one-carbon cycle, glycine synthesis (SOG) pathway and the biosynthesis of nucleotide precursors. *Synechocystis* sp. PCC 6803 was simulated to overproduce ethylene, while grown under mixotrophic conditions. Reaction fluxes (mmol/gDW/h) were predicted using pFBA. For interpretation of the references to color in this Fig legend, the reader is referred to the web version of this article.

Supplementary Fig. 19

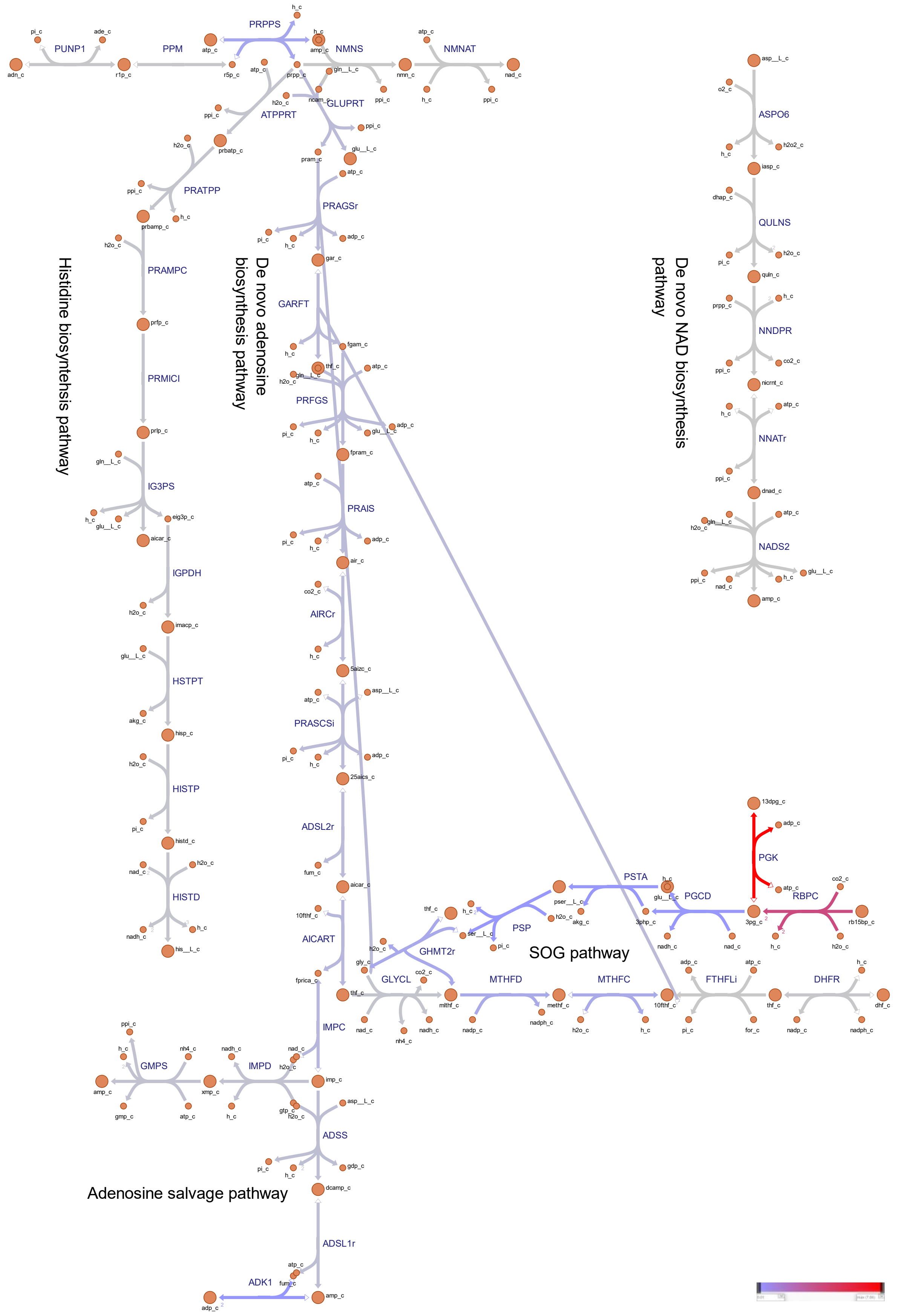


Supplementary Fig. 19 Metabolic flux map of auxiliary pathways enabling cellular energy and redox

homeostasis. The auxiliary pathways are composed of the serine, one-carbon cycle, glycine synthesis (SOG) pathway and the biosynthesis of nucleotide precursors. *Synechocystis* sp. PCC 6803 was simulated to overproduce 1-undecene, while grown under photoautotrophic conditions. Reaction fluxes (mmol/gDW/h) were predicted using pFBA. For interpretation of the references to color in this Fig legend, the reader is referred to the web version of this article.

**Supplementary Fig. 20**

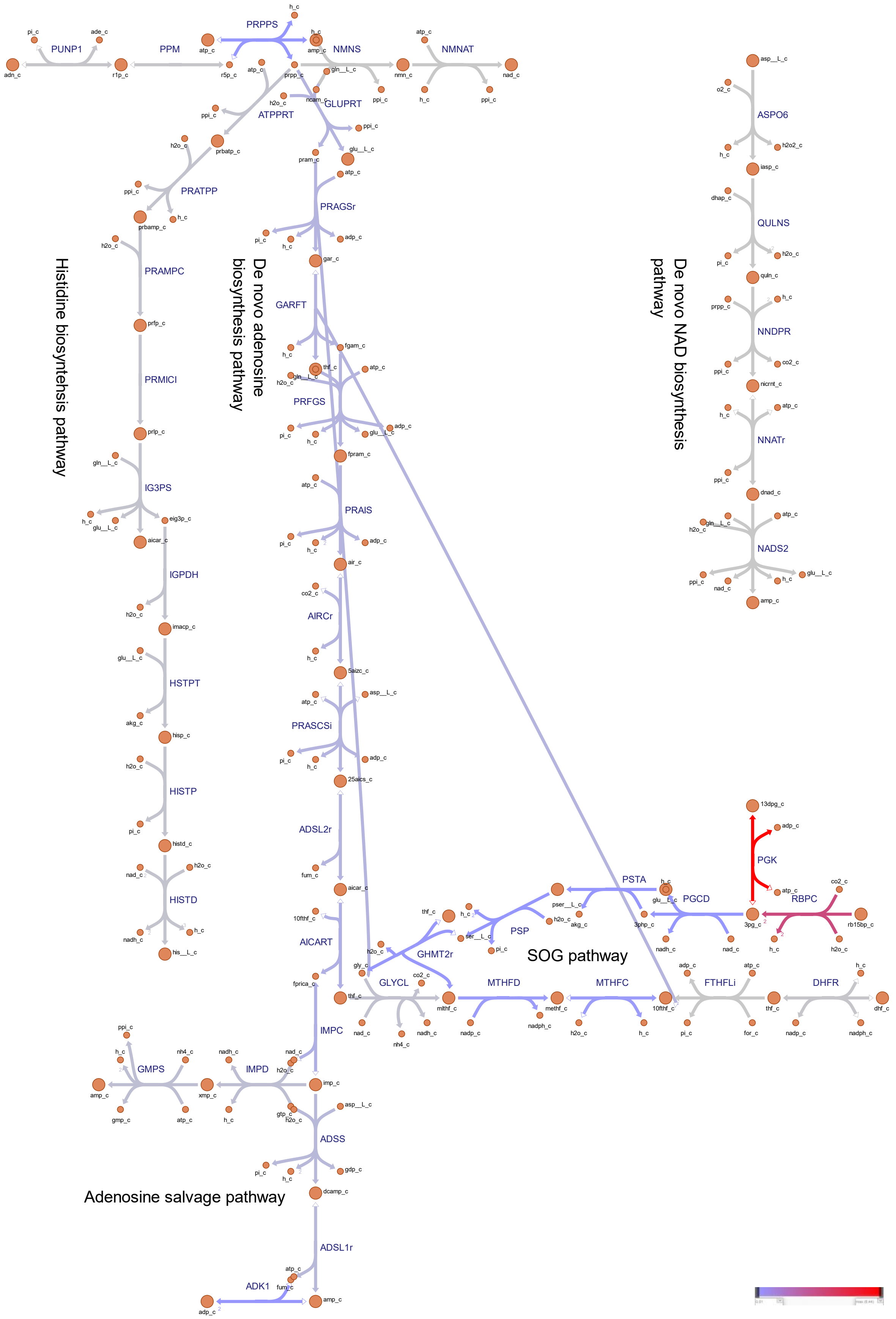


Supplementary Fig. 20 Metabolic flux map of auxiliary pathways enabling cellular energy and redox

homeostasis. The auxiliary pathways are composed of the serine, one-carbon cycle, glycine synthesis (SOG) pathway and the biosynthesis of nucleotide precursors. *Synechocystis* sp. PCC 6803 was simulated to overproduce 1-undecene, while grown under mixotrophic conditions. Reaction fluxes (mmol/gDW/h) were predicted using pFBA. For interpretation of the references to color in this Fig legend, the reader is referred to the web version of this article.

**Supplementary Table 1. Reactions added to the iJN678 model.**

| Name | Description | Stoichiometry | Reference |
|---|---|---|---|
| OXGDC | 2-oxoglutarate decarboxylase | akg_c + h_c --> co2_c + sucsal_c | [1–3] |
| PKETF | Phosphoketolase (fructose-6-phosphate utilizing) | f6p_c + pi_c --> actp_c + e4p_c + h2o_c | [4,5] |
| PKETX | Phosphoketolase (xylulose-5-phosphate utilizing) | pi_c + xu5p__D_c --> actp_c + g3p_c + h2o_c | [4,5] |
| GDH | Glucose dehydrogenase | glc__bD_c + h2o_c + nadp_c --> glcn_c + 2 h_c + nadph_c | [6] |
| GNK | Gluconokinase | atp_c + glcn_c --> 6pgc_c + adp_c + h_c | [6] |
| EDD | 6-phosphogluconate dehydratase | 6pgc_c --> 2ddg6p_c + h2o_c | [6] |
| EDA | 2-dehydro-3-deoxy-phosphogluconate aldolase | 2ddg6p_c --> g3p_c + pyr_c | [6] |
| PSTA | Phosphoserine transaminase | "3php_c + glu__L_c --> akg_c + pser__L_c | [7] |
| PSP | Phosphoserine phosphatase | h2o_c + pser__L_c --> 2 h_c + pi_c + ser__L_c | [7] |
| PAT1 | Prephenate transaminase (aspartate donor) | asp__L_c + h_c + pphn_c --> Largn_c + oaa_c | [8] |
| PAT2 | Prephenate transaminase (glutamate donor) | glu__L_c + h_c + pphn_c --> Largn_c + akg_c | [8] |
| PHEA | Prephenate dehydratase | Largn_c --> co2_c + h2o_c + phe__L_c | [8] |
| TYRA | Arogenate dehydrogenase | Largn_c + nadp_c --> co2_c + h_c + nadph_c + tyr__L_c | [8] |
| NADTRHD | NAD transhydrogenase | nad_c + nadph_c ⇌ nadh_c + nadp_c | [9] |
| NDH2_1p | NADH dehydrogenase 2 | h_c + nadh_c + pq_p --> nad_c + pqh2_p | [10,11] |
| ARTO | Alternative respiratory terminal oxidase | 2 h_c + 0.5 o2_p + pqh2_p --> h2o_p + pq_p | [10,11] |
| Flv2/4 | Flavodiiron 2/4 | 2 h_c + 0.5 o2_u + pqh2_u --> h2o_u + pq_u | [10,11] |
| CBFCpp | Cytochrome b6/f complex periplasm | Constrained to zero | [10,11] |
| CBFC2pp | Cytochrome b6/f complex periplasm | Constrained to zero | [10,11] |
| CYO1b2pp_syn | Cytochrome c oxidase | Constrained to zero | [10,11] |
| CYO1bpp_syn | Cytochrome c oxidase | Constrained to zero | [10,11] |
| CYO1b2_syn | Cytochrome c oxidase | Constrained to zero | [10,11] |
| NDH1_2p | NAD(P)H dehydrogenase | Constrained to zero | [10,11] |
| NDH1_1p | NAD(P)H dehydrogenase | Constrained to zero | [10,11] |
| NGAM | Non-growth associated ATP maintenance | atp_c + h2o_c --> adp_c + h_c + pi_c | |

**Supplementary Table 2. Reactions added to iJN678 to create iJN678_isobutene**

| Name | Description | Stoichiometry | Reference |
|---|---|---|---|
| KICD | alpha-ketoisocaproate dioxygenase/decarboxylase | 4mop_c + o2_c --> co2_c + m_3hivac_c | [12–14] |
| M3K | mevalonate-3-kinase | atp_c + m_3hivac_c --> adp_c + m_3piv_c | [12–14] |
| Isobutene_spontaneous | Isobutene spontaneous | m_3piv_c --> co2_c + isb_c + pi_c | |
| ISBt | Isobutene transport via diffusion (cytoplasm to extracellular)) | isb_c --> isb_e | |
| EX_isb_e | Isobutene exchange | isb_e --> | |

**Supplementary Table 3. Reactions added to iJN678 to create iJN678_isoprene**

| Name | Description | Stoichiometry | Reference |
|---|---|---|---|
| ISPs | Isoprene synthase | dmpp_c --> isp_c + ppi_c | [15] |
| ISPt | Isoprene transport via diffusion (cytoplasm to extracellular) | isp_c --> isp_e | |
| EX_isp_e | Isoprene exchange | isp_e --> | |

**Supplementary Table 4. Reactions added to iJN678 to create iJN678_ethylene**

| Name | Description | Stoichiometry | Reference |
|---|---|---|---|
| EFE | Ethylene-forming enzyme | 3 akg_c + arg__L_c + 3 h_c + 3 o2_c --> 1pyr5c_c + 7 co2_c + 2 ethylene_c + guanidine_c + 3 h2o_c + succ_c | [16] |
| GUANIDINEt | Guanidine transport via diffusion (cytoplasm to extracellular) | guanidine_c --> guanidine_e | |
| ETHYLENEt | Ethylene transport via diffusion (cytoplasm to extracellular) | ethylene_c --> eth_e | |
| EX_gua_e | Guanidine exchange | guanidine_e --> | |
| EX_eth_e | Ethylene exchange | eth_e --> | |

**Supplementary Table 5. Reactions added to iJN678 to create iJN678_1-undecene**

| Name | Description | Stoichiometry | Reference |
|---|---|---|---|
| tesA | Acyl-ACP thioesterase | ddcaACP_c + h2o_c --> ACP_c + dodecacid_c + h_c | [17] |
| UndB | Desaturase-like | dodecacid_c --> co2_c + h_c + und_c | [17] |
| UNDt | 1-undecene transport via diffusion (cytoplasm to extracellular) | und_c --> und_e | |
| EX_und_e | 1-undecene exchange | und_e --> | |

**Supplementary Table 6. Release of $CO_2$ in Synechocystis grown photoautotrophically and mixotrophically, when maximizing biomass and alkenes production.**

| Objective | auto | mixo |
|---|---|---|
| biomass | 3.81 | 3.26 |
| isobutene | 5.80 | 6.01 |
| isoprene | 4.41 | 4.35 |
| ethylene | 5.28 | 5.19 |
| 1-undecene | 3.99 | 3.32 |

*Estimated values are based on the flux-sum analysis using iJN678 metabolic model of *Synechocystis*, grown under light intensity of 45 mmol/gDW/h and bicarbonate uptake rate of 3.7 mmol/gDW/h, where the minimum cellular growth was subjected to 10% of the maximum growth rate. For mixotrophic condition, glucose uptake rate was constrained to 0.38 mmol/gDW/h. Values are presented in units of mmol/gDW/h. Auto, photoautotrophic; mixo, mixotrophic.

**Supplementary Table 7. Shadow prices and weighted costs of cofactor metabolites (ATP, NADPH and NADH) in biomass and alkene productivity, under photoautotrophic conditions.**

| Objective | ATP Shadow prices | NADPH Shadow prices | NADH Shadow prices | ATP Weighted costs | NADPH Weighted costs | NADH Weighted costs |
|---|---|---|---|---|---|---|
| **biomass** | -0.26 | -0.48 | -0.48 | -4.21 | -3.75 | -0.12 |
| **isobutene** | -2.36 | -4.43 | -4.39 | -40.68 | -48.50 | -3.08 |
| **isoprene** | -2.00 | -4.20 | -4.20 | -28.76 | -42.71 | -0.10 |
| **ethylene** | -2.29 | -4.27 | -4.24 | -35.16 | -36.41 | -4.13 |
| **1-undecene** | -1.04 | -1.94 | -1.93 | -15.01 | -21.32 | -0.03 |

**Supplementary Table 8. Shadow prices and weighted costs of cofactor metabolites (ATP, NADPH and NADH) in biomass and alkene productivity, under mixotrophic conditions.**

| Objective | ATP Shadow prices | NADPH Shadow prices | NADH Shadow prices | ATP Weighted costs | NADPH Weighted costs | NADH Weighted costs |
|---|---|---|---|---|---|---|
| **biomass** | -0.27 | -0.51 | -0.51 | -4.25 | -3.44 | -0.19 |
| **isobutene** | -2.35 | -4.41 | -4.36 | -41.84 | -48.62 | -4.35 |
| **isoprene** | -2.45 | -4.61 | -4.56 | -36.33 | -49.19 | -0.17 |
| **ethylene** | -2.29 | -4.27 | -4.24 | -35.25 | -30.85 | -5.89 |
| **1-undecene** | -1.03 | -1.94 | -1.94 | -15.01 | -21.02 | -0.05 |

**Supplementary Table 9. Predicted flux distributions of ADP-producing reactions in Synechocystis grown photoautotrophically and mixotrophically, when maximizing biomass and alkenes production.**

| ADP | | biomass | | isobutene | | isoprene | | ethylene | | 1-undecence | |
|---|---|---|---|---|---|---|---|---|---|---|---|
| | reaction | flux | percent | flux | percent | flux | percent | flux | percent | flux | percent |
| **auto** | PGK | 6.37 | 39.08 | 10.13 | 58.81 | 8.02 | 55.76 | 7.78 | 50.77 | 7.84 | 54.15 |
| | BIOMASS_Ec_SynAuto | 4.40 | 26.96 | 0.44 | 2.55 | 0.44 | 3.06 | 0.44 | 2.87 | 0.44 | 3.04 |
| | PRUK | 3.50 | 21.45 | 5.77 | 33.52 | 4.38 | 30.43 | 4.25 | 27.71 | 3.95 | 27.29 |
| | NO3abcpp | 0.74 | 4.56 | 0.07 | 0.43 | 0.07 | 0.52 | 1.05 | 6.84 | 0.07 | 0.51 |
| | ADK1 | 0.35 | 2.12 | 0.03 | 0.20 | 0.03 | 0.24 | 0.68 | 4.46 | 0.03 | 0.24 |
| | ACCOAC | 0.24 | 1.49 | 0.02 | 0.14 | 0.02 | 0.17 | 0.02 | 0.16 | 1.52 | 10.53 |
| | GLNS | 0.11 | 0.68 | 0.01 | 0.06 | 0.01 | 0.08 | 0.39 | 2.51 | 0.02 | 0.10 |
| | ASPK | 0.07 | 0.43 | 0.01 | 0.04 | 0.01 | 0.05 | 0.01 | 0.05 | 0.01 | 0.05 |
| | M3K | 0.00 | 0.00 | 0.68 | 3.94 | 0.00 | 0.00 | 0.00 | 0.00 | 0.00 | 0.00 |
| | CYTK1 | 0.02 | 0.10 | 0.00 | 0.01 | 0.67 | 4.68 | 0.00 | 0.01 | 0.00 | 0.01 |
| | CDPMEK | 0.01 | 0.09 | 0.00 | 0.01 | 0.67 | 4.68 | 0.00 | 0.01 | 0.00 | 0.01 |
| | CBMKr | 0.05 | 0.30 | 0.00 | 0.03 | 0.00 | 0.03 | 0.33 | 2.15 | 0.00 | 0.03 |
| | ACGK | 0.02 | 0.14 | 0.00 | 0.01 | 0.02 | 0.00 | 0.33 | 2.13 | 0.00 | 0.02 |
| | BCT1_syn | 0.00 | 0.00 | 0.00 | 0.00 | 0.00 | 0.00 | 0.00 | 0.00 | 0.54 | 3.71 |
| **mixo** | PGK | 4.66 | 29.89 | 9.81 | 54.99 | 7.40 | 49.97 | 6.46 | 42.00 | 6.43 | 44.13 |
| | BIOMASS_Ec_SynAuto | 4.73 | 30.34 | 0.47 | 2.65 | 0.47 | 3.19 | 0.47 | 3.08 | 0.47 | 3.25 |
| | PRUK | 2.80 | 17.93 | 5.91 | 33.14 | 4.25 | 28.68 | 3.73 | 24.29 | 3.26 | 22.39 |
| | NO3abcpp | 1.11 | 7.12 | 0.11 | 0.62 | 0.11 | 0.75 | 1.49 | 9.68 | 0.11 | 0.76 |
| | ADK1 | 0.52 | 3.31 | 0.05 | 0.29 | 0.05 | 0.35 | 0.97 | 6.31 | 0.05 | 0.35 |
| | ACCOAC | 0.36 | 2.33 | 0.04 | 0.20 | 0.04 | 0.25 | 0.04 | 0.24 | 2.13 | 14.64 |
| | GLNS | 0.17 | 1.07 | 0.02 | 0.09 | 0.02 | 0.11 | 0.83 | 5.42 | 0.02 | 0.16 |
| | ASPK | 0.11 | 0.68 | 0.01 | 0.06 | 0.01 | 0.07 | 0.01 | 0.07 | 0.01 | 0.07 |
| | HEX1 | 0.38 | 2.44 | 0.38 | 2.13 | 0.38 | 2.56 | 0.38 | 2.47 | 0.38 | 2.61 |
| | M3K | 0.00 | 0.00 | 0.96 | 5.38 | 0.00 | 0.00 | 0.00 | 0.00 | 0.00 | 0.00 |
| | CYTK1 | 0.02 | 0.15 | 0.00 | 0.01 | 1.01 | 6.79 | 0.00 | 0.02 | 0.00 | 0.02 |
| | CDPMEK | 0.02 | 0.14 | 0.00 | 0.01 | 1.01 | 6.79 | 0.00 | 0.01 | 0.01 | 0.05 |
| | CBMKr | 0.07 | 0.46 | 0.01 | 0.04 | 0.01 | 0.05 | 0.47 | 3.03 | 0.01 | 0.05 |
| | ACGK | 0.03 | 0.21 | 0.00 | 0.02 | 0.00 | 0.02 | 0.46 | 3.01 | 0.00 | 0.02 |
| | BCT1_syn | 0.00 | 0.00 | 0.00 | 0.00 | 0.00 | 0.00 | 0.00 | 0.00 | 1.62 | 11.12 |

*Presented are percentage (%) and the corresponding flux (mmol/gDW/h) of reactions contributing to the generation of ADP. For convenience, presented are only reactions whose flux consists of XXX percent or higher. The complete list of reactions, including their abbreviations, subsystem, minimum and maximum bounds, and stoichiometry can be found in table SXXX.

**Supplementary Table 10. Predicted flux distributions of ADP-consuming reactions in Synechocystis grown photoautotrophically and mixotrophically, when maximizing biomass and alkenes production.**

| ADP | - | biomass | | isobutene | | isoprene | | ethylene | | 1-undecence | |
|---|---|---|---|---|---|---|---|---|---|---|---|
| | reaction | flux | percent | flux | percent | flux | percent | flux | percent | flux | percent |
| **auto** | ATPSu | -16.03 | 98.34 | -15.84 | 91.97 | -14.35 | 99.81 | -15.04 | 98.09 | -14.46 | 99.95 |
| | PYK | -0.13 | 0.82 | -1.37 | 7.95 | -0.01 | 0.09 | 0.00 | 0.00 | -0.01 | 0.04 |
| | ATPS4rpp_1 | -0.13 | 0.81 | -0.01 | 0.08 | -0.01 | 0.09 | -0.29 | 1.90 | 0.00 | 0.00 |
| | URIDK2r | 0.00 | 0.01 | 0.00 | 0.00 | 0.00 | 0.00 | 0.00 | 0.00 | 0.00 | 0.00 |
| | RNDR1 | 0.00 | 0.01 | 0.00 | 0.00 | 0.00 | 0.00 | 0.00 | 0.00 | 0.00 | 0.00 |
| | PPK2 | 0.00 | 0.00 | 0.00 | 0.00 | 0.00 | 0.00 | 0.00 | 0.00 | 0.00 | 0.00 |
| **mixo** | ATPSu | -15.28 | 97.93 | -15.89 | 89.11 | -14.80 | 99.86 | -15.04 | 97.84 | -14.46 | 99.20 |
| | PYK | -0.20 | 1.28 | -1.94 | 10.88 | -0.02 | 0.14 | 0.00 | 0.00 | -0.01 | 0.07 |
| | ATPS4rpp_1 | -0.12 | 0.74 | 0.00 | 0.00 | 0.00 | 0.00 | -0.33 | 2.16 | -0.11 | 0.73 |
| | URIDK2r | 0.00 | 0.02 | 0.00 | 0.00 | 0.00 | 0.00 | 0.00 | 0.00 | 0.00 | 0.00 |
| | RNDR1 | 0.00 | 0.02 | 0.00 | 0.00 | 0.00 | 0.00 | 0.00 | 0.00 | 0.00 | 0.00 |
| | PPK2 | 0.00 | 0.00 | 0.00 | 0.00 | 0.00 | 0.00 | 0.00 | 0.00 | 0.00 | 0.00 |

*Presented are percentage (%) and the corresponding flux (mmol/gDW/h) of reactions contributing to the consumption of ADP. The complete list of reactions, including their abbreviations, subsystem, minimum and maximum bounds, and stoichiometry can be found in table SXXX.

**Supplementary Table 11. Predicted flux distributions of AMP-producing reactions in Synechocystis grown photoautotrophically and mixotrophically, when maximizing biomass and alkenes production.**

| AMP | | biomass | | isobutene | | isoprene | | ethylene | | 1-undecence | |
|---|---|---|---|---|---|---|---|---|---|---|---|
| | reaction | flux | percent | flux | percent | flux | percent | flux | percent | flux | percent |
| **auto** | PRPPS | 0.07 | 39.47 | 0.01 | 39.47 | 0.01 | 39.47 | 0.01 | 1.99 | 0.01 | 39.47 |
| | ARGSS | 0.02 | 12.98 | 0.00 | 12.98 | 0.00 | 12.98 | 0.33 | 95.61 | 0.00 | 12.98 |
| | ADSL1r | 0.02 | 12.01 | 0.00 | 12.01 | 0.00 | 12.01 | 0.00 | 0.61 | 0.00 | 12.01 |
| | ASNS1 | 0.02 | 8.90 | 0.00 | 8.90 | 0.00 | 8.90 | 0.00 | 0.45 | 0.00 | 8.90 |
| | BPNT | 0.02 | 8.87 | 0.00 | 8.87 | 0.00 | 8.87 | 0.00 | 0.45 | 0.00 | 8.87 |
| | GLUTRS | 0.01 | 8.43 | 0.00 | 8.43 | 0.00 | 8.43 | 0.00 | 0.43 | 0.00 | 8.43 |
| | GMPS | 0.01 | 7.54 | 0.00 | 7.54 | 0.00 | 7.54 | 0.00 | 0.38 | 0.00 | 7.54 |
| | ADPT | 0.00 | 1.52 | 0.00 | 1.52 | 0.00 | 1.52 | 0.00 | 0.08 | 0.00 | 1.52 |
| | NADS2 | 0.00 | 0.11 | 0.00 | 0.11 | 0.00 | 0.11 | 0.00 | 0.01 | 0.00 | 0.11 |
| | SUCBZL | 0.00 | 0.08 | 0.00 | 0.08 | 0.00 | 0.08 | 0.00 | 0.00 | 0.00 | 0.08 |
| | HPPK | 0.00 | 0.04 | 0.00 | 0.04 | 0.00 | 0.04 | 0.00 | 0.00 | 0.00 | 0.04 |
| | PANTS | 0.00 | 0.03 | 0.00 | 0.03 | 0.00 | 0.03 | 0.00 | 0.00 | 0.00 | 0.03 |
| **mixo** | PRPPS | 0.10 | 39.47 | 0.01 | 39.47 | 0.01 | 39.47 | 0.01 | 2.10 | 0.01 | 39.47 |
| | ARGSS | 0.03 | 12.98 | 0.00 | 12.98 | 0.00 | 12.98 | 0.46 | 95.36 | 0.00 | 12.98 |
| | ADSL1r | 0.03 | 12.01 | 0.00 | 12.01 | 0.00 | 12.01 | 0.00 | 0.64 | 0.00 | 12.01 |
| | ASNS1 | 0.02 | 8.90 | 0.00 | 8.90 | 0.00 | 8.90 | 0.00 | 0.47 | 0.00 | 8.90 |
| | BPNT | 0.02 | 8.87 | 0.00 | 8.87 | 0.00 | 8.87 | 0.00 | 0.47 | 0.00 | 8.87 |
| | GLUTRS | 0.02 | 8.43 | 0.00 | 8.43 | 0.00 | 8.43 | 0.00 | 0.45 | 0.00 | 8.43 |
| | GMPS | 0.02 | 7.54 | 0.00 | 7.54 | 0.00 | 7.54 | 0.00 | 0.40 | 0.00 | 7.54 |
| | ADPT | 0.00 | 1.52 | 0.00 | 1.52 | 0.00 | 1.52 | 0.00 | 0.08 | 0.00 | 1.52 |
| | NADS2 | 0.00 | 0.11 | 0.00 | 0.11 | 0.00 | 0.11 | 0.00 | 0.01 | 0.00 | 0.11 |
| | SUCBZL | 0.00 | 0.08 | 0.00 | 0.08 | 0.00 | 0.08 | 0.00 | 0.00 | 0.00 | 0.08 |
| | HPPK | 0.00 | 0.04 | 0.00 | 0.04 | 0.00 | 0.04 | 0.00 | 0.00 | 0.00 | 0.04 |
| | PANTS | 0.00 | 0.03 | 0.00 | 0.03 | 0.00 | 0.03 | 0.00 | 0.00 | 0.00 | 0.03 |

*Presented are percentage (%) and the corresponding flux (mmol/gDW/h) of reactions contributing to the generation of AMP. The complete list of reactions, including their abbreviations, subsystem, minimum and maximum bounds, and stoichiometry can be found in table SXXX.

**Supplementary Table 12. Predicted flux distributions of AMP-consuming reactions in Synechocystis grown photoautotrophically and mixotrophically, when maximizing biomass and alkenes production.**

| AMP | - | biomass | | isobutene | | isoprene | | ethylene | | 1-undecence | |
|---|---|---|---|---|---|---|---|---|---|---|---|
| | reaction | flux | percent | flux | percent | flux | percent | flux | percent | flux | percent |
| auto | ADK1 | -0.17 | 100.00 | -0.02 | 100.00 | -0.02 | 100.00 | -0.34 | 100.00 | -0.02 | 100.00 |
| mixo | ADK1 | -0.26 | 100.00 | -0.03 | 100.00 | -0.03 | 100.00 | -0.48 | 100.00 | -0.03 | 100.00 |

*Presented are percentage (%) and the corresponding flux (mmol/gDW/h) of reactions contributing to the consumption of AMP. The complete list of reactions, including their abbreviations, subsystem, minimum and maximum bounds, and stoichiometry can be found in Supplementary Data.

**Supplementary Table 13. Predicted flux distributions of Pi-producing reactions in Synechocystis grown photoautotrophically and mixotrophically, when maximizing biomass and alkenes production.**

| Pi | | biomass | - | isobutene | - | isoprene | - | ethylene | - | 1-undecence | - |
|---|---|---|---|---|---|---|---|---|---|---|---|
| | reaction | flux | percent | flux | percent | flux | percent | flux | percent | flux | percent |
| **auto** | GAPDi_nadp | 6.37 | 38.39 | 10.13 | 61.11 | 8.02 | 55.65 | 7.78 | 47.61 | 7.84 | 48.06 |
| | BIOMASS_Ec_SynAuto | 4.40 | 26.49 | 0.44 | 2.65 | 0.44 | 3.05 | 0.44 | 2.69 | 0.44 | 2.70 |
| | FBP | 1.39 | 8.39 | 2.17 | 13.11 | 1.48 | 10.28 | 1.76 | 10.79 | 1.94 | 11.89 |
| | SBP | 1.32 | 7.95 | 2.17 | 13.07 | 1.47 | 10.23 | 1.76 | 10.74 | 1.93 | 11.85 |
| | NO3abcpp | 0.74 | 4.48 | 0.07 | 0.45 | 0.07 | 0.52 | 1.05 | 6.42 | 0.07 | 0.46 |
| | PPA | 0.57 | 3.41 | 0.06 | 0.34 | 2.74 | 19.02 | 0.71 | 4.32 | 0.06 | 0.35 |
| | PTAr | 0.44 | 2.67 | 0.72 | 4.36 | 0.04 | 0.31 | 1.02 | 6.23 | 1.84 | 11.31 |
| | ACCOAC | 0.24 | 1.46 | 0.02 | 0.15 | 0.02 | 0.17 | 0.02 | 0.15 | 1.52 | 9.35 |
| | PPC | 0.23 | 1.41 | 0.02 | 0.14 | 0.02 | 0.16 | 0.70 | 4.26 | 0.03 | 0.19 |
| | PIuabcpp | 0.11 | 0.69 | 0.01 | 0.07 | 0.01 | 0.08 | 0.01 | 0.07 | 0.01 | 0.07 |
| | GLNS | 0.11 | 0.67 | 0.01 | 0.07 | 0.01 | 0.08 | 0.39 | 2.36 | 0.02 | 0.09 |
| | PSP | 0.10 | 0.57 | 0.01 | 0.06 | 0.01 | 0.07 | 0.01 | 0.06 | 0.01 | 0.06 |
| | ASAD | 0.07 | 0.43 | 0.01 | 0.04 | 0.01 | 0.05 | 0.01 | 0.04 | 0.01 | 0.04 |
| | Isobutene_spontaneous | 0.00 | 0.00 | 0.68 | 4.09 | 0.00 | 0.00 | 0.00 | 0.00 | 0.00 | 0.00 |
| | AGPR | 0.02 | 0.14 | 0.00 | 0.01 | 0.00 | 0.02 | 0.33 | 2.00 | 0.00 | 0.01 |
| | OCBT | 0.02 | 0.14 | 0.00 | 0.01 | 0.00 | 0.02 | 0.33 | 2.00 | 0.00 | 0.01 |
| | BCT1_syn | 0.00 | 0.00 | 0.00 | 0.00 | 0.00 | 0.00 | 0.00 | 0.00 | 0.54 | 3.30 |
| **mixo** | GAPDi_nadp | 4.66 | 29.08 | 9.81 | 57.97 | 7.40 | 49.82 | 6.46 | 38.40 | 6.43 | 37.51 |
| | BIOMASS_Ec_SynAuto | 4.73 | 29.51 | 0.47 | 2.80 | 0.47 | 3.19 | 0.47 | 2.82 | 0.47 | 2.76 |
| | FBP | 0.87 | 5.40 | 1.94 | 11.48 | 1.07 | 7.19 | 1.35 | 8.06 | 1.58 | 9.20 |
| | SBP | 1.16 | 7.25 | 2.31 | 13.67 | 1.44 | 9.69 | 1.73 | 10.27 | 1.95 | 11.37 |
| | NO3abcpp | 1.11 | 6.93 | 0.11 | 0.66 | 0.11 | 0.75 | 1.49 | 8.85 | 0.11 | 0.65 |
| | PPA | 0.79 | 4.96 | 0.08 | 0.47 | 4.09 | 27.55 | 1.00 | 5.93 | 0.08 | 0.46 |
| | PTAr | 0.66 | 4.13 | 1.03 | 6.07 | 0.07 | 0.45 | 1.44 | 8.58 | 2.58 | 15.07 |
| | ACCOAC | 0.36 | 2.26 | 0.04 | 0.21 | 0.04 | 0.24 | 0.04 | 0.22 | 2.13 | 12.45 |
| | PPC | 0.35 | 2.18 | 0.03 | 0.21 | 0.03 | 0.24 | 0.97 | 5.79 | 0.05 | 0.26 |
| | PIuabcpp | 0.17 | 1.06 | 0.02 | 0.10 | 0.02 | 0.11 | 0.02 | 0.10 | 0.02 | 0.10 |
| | GLNS | 0.17 | 1.04 | 0.02 | 0.10 | 0.02 | 0.11 | 0.83 | 4.96 | 0.02 | 0.13 |
| | PSP | 0.14 | 0.89 | 0.01 | 0.08 | 0.01 | 0.10 | 0.01 | 0.08 | 0.01 | 0.08 |
| | ASAD | 0.11 | 0.66 | 0.01 | 0.06 | 0.01 | 0.07 | 0.01 | 0.06 | 0.01 | 0.06 |
| | Isobutene_spontaneous | 0.00 | 0.00 | 0.96 | 5.68 | 0.00 | 0.00 | 0.00 | 0.00 | 0.00 | 0.00 |
| | AGPR | 0.03 | 0.21 | 0.00 | 0.02 | 0.00 | 0.02 | 0.46 | 2.75 | 0.00 | 0.02 |
| | OCBT | 0.03 | 0.21 | 0.00 | 0.02 | 0.00 | 0.02 | 0.46 | 2.75 | 0.00 | 0.02 |
| | BCT1_syn | 0.00 | 0.00 | 0.00 | 0.00 | 0.00 | 0.00 | 0.00 | 0.00 | 1.62 | 9.45 |

*Presented are percentage (%) and the corresponding flux (mmol/gDW/h) of reactions contributing to the generation of Pi. The complete list of reactions, including their abbreviations, subsystem, minimum and maximum bounds, and stoichiometry can be found in table SXXX.

**Supplementary Table 14. Predicted flux distributions of Pi-consuming reactions in Synechocystis grown photoautotrophically and mixotrophically, when maximizing biomass and alkenes production.**

| Pi | - | biomass | | isobutene | | isoprene | | ethylene | | 1-undecence | |
|---|---|---|---|---|---|---|---|---|---|---|---|
| | reaction | flux | percent | flux | percent | flux | percent | flux | percent | flux | percent |
| **auto** | ATPSu | -16.03 | 96.62 | -15.84 | 95.57 | -14.35 | 99.61 | -15.04 | 91.99 | -14.46 | 88.70 |
| | PKETF | -0.43 | 2.57 | -0.72 | 4.35 | -0.04 | 0.30 | -1.02 | 6.22 | -1.84 | 11.30 |
| | ATPS4rpp_1 | -0.13 | 0.79 | -0.01 | 0.08 | -0.01 | 0.09 | -0.29 | 1.78 | 0.00 | 0.00 |
| | PUNP1 | 0.00 | 0.01 | 0.00 | 0.00 | 0.00 | 0.00 | 0.00 | 0.00 | 0.00 | 0.00 |
| | MTAP | 0.00 | 0.00 | 0.00 | 0.00 | 0.00 | 0.00 | 0.00 | 0.00 | 0.00 | 0.00 |
| **mixo** | ATPSu | -15.28 | 95.27 | -15.89 | 93.94 | -14.80 | 99.57 | -15.04 | 89.46 | -14.46 | 84.33 |
| | PKETF | -0.64 | 3.99 | -1.02 | 6.05 | -0.06 | 0.43 | -1.44 | 8.57 | -1.96 | 11.40 |
| | ATPS4rpp_1 | -0.12 | 0.72 | 0.00 | 0.00 | 0.00 | 0.00 | -0.33 | 1.97 | -0.11 | 0.62 |
| | PUNP1 | 0.00 | 0.02 | 0.00 | 0.00 | 0.00 | 0.00 | 0.00 | 0.00 | 0.00 | 0.00 |
| | MTAP | 0.00 | 0.01 | 0.00 | 0.00 | 0.00 | 0.00 | 0.00 | 0.00 | 0.00 | 0.00 |
| | PKETX | 0.00 | 0.00 | 0.00 | 0.00 | 0.00 | 0.00 | 0.00 | 0.00 | -0.63 | 3.65 |

### References


1. Zhang, S. & Bryant, D. A. The Tricarboxylic Acid Cycle in Cyanobacteria. *Science (80-. ).* **334**, 1551–1553 (2011).
2. Steinhauser, D., Fernie, A. R. & Araújo, W. L. Unusual cyanobacterial TCA cycles: not broken just different. *Trends Plant Sci.* **17**, 503–509 (2012).
3. Xiong, W., Brune, D. & Vermaas, W. F. J. The γ-aminobutyric acid shunt contributes to closing the tricarboxylic acid cycle in S *ynechocystis* sp. PCC 6803. *Mol. Microbiol.* **93**, 786–796 (2014).
4. Xiong, W. *et al.* Phosphoketolase pathway contributes to carbon metabolism in cyanobacteria. *Nat. Plants* **2**, 15187 (2016).
5. Bachhar, A. & Jablonsky, J. A new insight into role of phosphoketolase pathway in *Synechocystis* sp. PCC 6803. *Sci. Rep.* **10**, 22018 (2020).
6. Chen, X. *et al.* The Entner–Doudoroff pathway is an overlooked glycolytic route in cyanobacteria and plants. *Proc. Natl. Acad. Sci.* **113**, 5441–5446 (2016).
7. Klemke, F. *et al.* Identification of the light-independent phosphoserine pathway as an additional source of serine in the cyanobacterium *Synechocystis* sp. PCC 6803. *Microbiology* **161**, 1050–1060 (2015).
8. Bonner, C. A., Jensen, R. A., Gander, J. E. & Keyhani, N. O. A core catalytic domain of the TyrA protein family: arogenate dehydrogenase from Synechocystis. *Biochem. J.* **382**, 279–291 (2004).
9. Kämäräinen, J. *et al.* Pyridine nucleotide transhydrogenase Pnt AB is essential for optimal growth and photosynthetic integrity under low-light mixotrophic conditions in *Synechocystis* sp. PCC 6803. *New Phytol.* **214**, 194–204 (2017).

10. Lea-Smith, D. J., Bombelli, P., Vasudevan, R. & Howe, C. J. Photosynthetic, respiratory and extracellular electron transport pathways in cyanobacteria. *Biochim. Biophys. Acta - Bioenerg.* **1857**, 247–255 (2016).

11. Cooley, J. W. & Vermaas, W. F. J. Succinate Dehydrogenase and Other Respiratory Pathways in Thylakoid Membranes of *Synechocystis* sp. Strain PCC 6803: Capacity Comparisons and Physiological Function. *J. Bacteriol.* **183**, 4251–4258 (2001).

12. Mustila, H., Kugler, A. & Stensjö, K. Isobutene production in *Synechocystis* sp. PCC 6803 by introducing α-ketoisocaproate dioxygenase from Rattus norvegicus. *Metab. Eng. Commun.* **12**, e00163 (2021).

13. Baldwin, J. E. *et al.* 4-Hydroxyphenylpyruvate dioxygenase appears to display α-ketoisocaproate dioxygenase activity in rat liver. *Bioorg. Med. Chem. Lett.* **5**, 1255–1260 (1995).

14. Rossoni, L., Hall, S. J., Eastham, G., Licence, P. & Stephens, G. The Putative Mevalonate Diphosphate Decarboxylase from Picrophilus torridus Is in Reality a Mevalonate-3-Kinase with High Potential for Bioproduction of Isobutene. *Appl. Environ. Microbiol.* **81**, 2625–2634 (2015).

15. Lindberg, P., Park, S. & Melis, A. Engineering a platform for photosynthetic isoprene production in cyanobacteria, using Synechocystis as the model organism. *Metab. Eng.* **12**, 70–79 (2010).

16. Ungerer, J. *et al.* Sustained photosynthetic conversion of $CO_2$ to ethylene in recombinant cyanobacterium *Synechocystis* 6803. *Energy Environ. Sci.* **5**, 8998 (2012).

17. Yunus, I. S. *et al.* Synthetic metabolic pathways for photobiological conversion of $CO_2$ into hydrocarbon fuel. *Metab. Eng.* **49**, 201–211 (2018).
